# Does Bodily Action Shape Spatial Representation? Evidence from Virtual Reality, Sensory Augmentation and Map Learning

**DOI:** 10.1101/2023.10.15.562402

**Authors:** Nicolas Kuske, Viviane Clay

## Abstract

Spatial relations can be defined with respect to the body (egocentric) or among environmental objects only (allocentric). Egocentric relations are necessarily transformed through bodily action. To what extent allocentric cognitive representations are shaped by the body remains unclear. In our study, participants navigate a virtual-reality (VR) city over multiple days in one of three embodiment conditions. In two VR conditions, the participants sit on a swivel chair actively changing navigation direction through bodily rotation. In one of these groups the VR participants wear a sensory augmentation belt which indicates the cardinal direction of north through vibration. The third group of participants navigates a two-dimensional map of the city. After each exploration session, participants complete tasks asking for allocentric spatial relations. We find that the performance in the spatial tasks interacts with the duration of exploration time and the embodiment condition. These findings indicate allocentric spatial representations to be structured by bodily action.

---

Starting around the second half of the 20th century, specialists across disciplines from philosophy to robotics sought to reevaluate the role of bodily action for cognition. Motoric bodily action, those scientists argued, is not an uninteresting end product of cognitive processes. Rather, it should be considered a key player at the very core of cognition itself. Changes in sensory activity, for example, are not only structured by the agent’s environment but also by the agent’s bodily shape and motor actions (Lenay, Canu, & Villon, 1997; Merleau-Ponty, 1945; Nagel, Christine, Kringe, Märtin & König, 2005; O’Regan & Noë, 2001; Prinz, 1997; Schumann & O’Regan, 2017; White, Saunders, Scadden, Bach-y-Rita & Collins, 1970). As early as 1950, James Gibson explored the importance of the structure of sensorimotor relations for general cognition. He thought these relations can help explain the perceived relative immobility of the space surrounding us.

Gibson (1979) later coined the term “affordances,” by which he claims that our perception of an object depends on what we can do with that object (cf. Koffka, 1935). In recent years, a wide body of research suggests that perception and the planning of bodily action share a common neural code (Bonner & Epstein, 2017; Cisek & Kalaska, 2010; Etzel, Gazzola, & Keysers, 2008; Melnik, Hairston, Ferris, & König, 2017; Prinz, 1997). Even the concept of memory might be best understood as “the encoding of patterns of possible physical interaction with a three-dimensional world” (Glenberg, 1997, p. 1). Building upon these ideas, Engel, Maye, Kurthen and König (2013) present evidence from diverse areas of research, culminating in their conjecture that even abstract reasoning is fundamentally grounded in bodily action (cf. Lakoff & Johnson, 1980).

The aforementioned research falls into the broad category of embodied enactivism. While there is general consensus that some basic cognitive processes are well described in this framework, critics warn against an exaggerated embrace of these ideas (Clark, 1999; Goldinger, Papesh, Barnhart, Hansen, & Hout, 2016; Schlicht & Starzak, 2019). One major point of contention for Goldinger et al. (2016) is the vagueness of many embodiment claims. The researchers ask how to express embodiment in a formal model. Secondly, Goldinger and colleagues claim that the embodied framework does not help to illuminate the majority of findings cognitive science tries to explain. They list, for example, Stroop interference, the attentional blink and short-term memory scanning (in the same order, Stroop, 1935; Shapiro, Raymond & Arnell, 1997; Sternberg, 1966). In a similar vein, cognitive reasoning processes seem to involve computations over mental representations (Clark, 1999; Wilson, 2002). Proponents of embodiment still lack a detailed account of how these reasoning processes fit into a framework relying heavily on the body (Schlicht & Starzak, 2019).

Our work tries to shed new light on the relationship of representation and embodiment in spatial cognition. We focus on spatial cognition because the ability to navigate in a large-scale environment can be considered a paradigmatic example of a “representation-hungry” problem (Clark, 1999). One seldom starts walking into an unfamiliar part of town without first making a (mental) plan of how to reach a certain goal location. Recent developments in cognitive and neuroscience further bolster the importance of this example. Numerous researchers have argued that the processes involving successful spatial navigation lie at the very heart of cognitive reasoning itself (Bellmund, Gärdenfors, Moser & Doeller, 2018; Buzsáki & Moser, 2013; Constantinescu, O’Reilly & Behrens, 2016; Epstein, Patai, Julian & Spiers, 2017; Schiller et al., 2015). Indeed, the idea of a strong link between navigation and general problem solving in cognition is not new (Siegel & White, 1975; Tolman, 1948). Finally, spatial exploration naturally involves our complete body. Spatial cognition seems well suited to elucidate the relationship of representation and embodiment.

## Egocentric Versus Allocentric

Contemporary research on spatial cognition commonly distinguishes between two categories of spatial relations, namely egocentric and allocentric (Burgess, 2007; Feigenbaum & Rolls, 1991; Klatzky, 1998; Wolbers & Wiener, 2014). Spatial relations among parts of our extracranial body or among objects in the environment in relation to parts of our body are called egocentric (or deictic, idiothetic, body centered). For egocentric representations our body, literally, is the frame of reference. Spatial relations exclusively among objects which are not part of our body are called allocentric (or extrinsic, exocentric, geocentric, world centered). Consequently, only objects, or directions defined through objects in the environment can be reference frames for allocentric code.

While the difference in the spatial relations can be easily defined, how it maps onto neuronal circuits is debated. There is overwhelming evidence that spatial relations which can be overseen from one location, also referred to as a vista space, can be represented in both the egocentric and the allocentric frame of reference (Burgess, 2007). Klaus Gramann (2013) emphasizes differences in individual proclivities toward a preferred reference frame during navigation. Furthermore, the neural bases of both representations overlap to a large extent (Ekstrom et al., 2003; Ekstrom, Arnold, & Laria, 2014; Feigenbaum & Rolls, 1991). These difficulties are further illustrated in the ongoing debate over how to interpret findings in numerous studies investigating egocentric or allocentric processing (Burgess, 2007; Banta Lavenex & Lavenex, 2009; Wolbers & Wiener, 2014). In fact, even the representational distinction itself, is disputed (Bennett, 1996; Filimon, 2015). The terminological distinction is, nevertheless, adopted in our study to emphasize that we investigate cognitive processes which involve spatial relations among environmental objects. Tasks which involve mostly egocentric spatial relations include finding a small object, like an apple, in a cluttered environment, like an office, and picking it up. The fluidity and speed of humans in such simple picking challenges is still unmatched by modern-day robots (Lawrence, 2017; Leitner, 2018). We can conclude that the involved representations have the property of allowing quick, goal-directed action. In contrast, tasks which involve allocentric spatial relations are often time consuming. Spatial decision latency, for example, has been reported to correlate with angular difference when participants had to point to objects located within a large environment which cannot be overseen from one point of view (Evans & Pezdek, 1980; Sholl, Kenny & DellaPorta, 2006).

The most remarkable property of representations involving allocentric relations is revealed in large environments which consist of multiple separated vista spaces, like a town. Some animals including humans are able to point in the direction of their current goal location despite never having seen or walked this path before. More often than not, the result of the pointing gesture is only a very rough approximation of how the crow would fly (Warren, 2019). Nevertheless, this cognitive ability might imply that representations of allocentric spatial relations are dissociable from embodied action. Spatial navigation in large environments might not fit in an enacted framework.

## Allocentric Spatial Knowledge

It is generally agreed upon that knowledge about spatial relations in a large environment is built up through, at least, three processes. Namely, memorizing landmarks, acquiring route knowledge, and connecting those to attain allocentric survey knowledge (McNaughton, Battaglia, Jensen, Moser & Moser, 2006; Siegel & White, 1975).

A landmark, here, does not need to be exceptionally large. Any immobile object or place which left an impression on the explorer such that she will remember it on subsequent appearance, or can represent it mentally even in its absence, can be considered a landmark. Representations underlying route knowledge combine landmarks and directional information. It is assumed that this directional information comes from bodily action and/or visual flow experienced at the landmarks (Chrastil, 2013). Route knowledge is the ability to sequentially represent (already experienced) landmark-action pairs in the correct order to reach the goal landmark (Siegel & White, 1975). When the animal has acquired enough experience in the new environment to mentally deduce paths to the goal location which it never took before, we speak of a global, survey-like representation, also known as cognitive map (Tolman, 1948). It remains unclear, however, to what extent the processes on the way to survey knowledge are actually separate and if these are the only processes involved (Chrastil, 2013; Ishikawa & Montello, 2006; Montello, 1998).

On a neuronal level, certain activity patterns have been associated with survey knowledge. After sufficient exploration time in a new environment some cells begin to increase their firing rate whenever an animal moves across a certain location. These cells are known as place cells (Ekstrom et al., 2003; O’Keefe & Dostrovsky, 1971). There are also cells the firing rate of which, integrated over time, makes up a regular hexagonal grid-like pattern covering the traversed space (Doeller,Barry, & Burgess, 2010; Hafting, Fyhn, Molden, Moser, M.B., & Moser, E. I., 2005). These cells are called grid cells. The emergence of place and grid cells is generally interpreted as evidence that a cognitive map, or survey knowledge, of the new environment has been acquired (Moser,E. I., Kropff, & Moser M.B., 2008; O’Keefe & Nadel, 1978; Wills, Cacucci, Burgess, & O’Keefe, 2010).

One factor that influences the allocentric spatial knowledge we acquire in large environments is the purpose of our experience. If we intent to learn spatial relations, for example to navigate to a goal more efficiently, we gather more comprehensive knowledge about our surroundings than when we do not care where we are. While this might seem trivial, the intention related difference in spatial knowledge assessed with behavioral measures like estimation of goal direction is remarkably low (Burte & Montello, 2017; Chrastil & Warren, 2011). Thus, contrary to intuition, allocentric spatial representations seem to form largely automatic, without the need for intention.

Research on large-scale spatial cognition has repeatedly found evidence for subgroups of spatial learners. Some people are very adept at acquiring survey knowledge of a new environment. Others are remarkably poor. These individual differences are not statistical noise but remain consistent over longitudinal studies. In one well known study by Ishikawa and Montello (2006), participants were driven along separate roads through the outskirts of a town. Even after ten such drives a considerable number of participants still did not acquire comprehensive survey knowledge. Using large virtual environments Weisberg and colleagues (2014, 2016) could repeatedly identify three distinct groups of spatial aptitude. They further found that questionnaire data on self-assessed spatial ability, or “sense of direction,” correlates well with these groups. Burte and Montello (2017) as well as He, McNamara and Brown (2019) also report self-assessed spatial aptitude to inform different groups of spatial learners. The ability to succeed at tasks involving allocentric spatial relations differs consistently in between individuals.

## Allocentric Knowledge and Body

Proprioception in connection with motor activity can help to increase the quality of spatial knowledge acquired while exploring an environment (Chrastil & Warren, 2013; Ruddle, Volkova, Mohler & Buelthoff, 2011; Waller, Loomis & Haun 2004). The degree to which this is the case however is still debated. For instance, a recent study shows participants who change movement direction through whole body rotation, but only need to lean forward to increase speed, acquire spatial knowledge of a similar quality as free walking participants. Only bodily rotation as motor action, on the other hand, left participants performing worse (Nguyen-Vo, Riecke, Stuerzlinger, Pham & Kruijff, 2019). Evidently, even small proprioceptive and vestibular activity changes in combination with motor activity are sufficient to significantly increase the quality of allocentric spatial knowledge. An unexpected finding, it still corroborates the role of embodied processing. Generally, integrating information from different modalities into one coherent structure is a highly complex task (Cao, Summerfield, Park, Giordano, & Kayser, 2019; Chen, McNamara, Kelly, & Wolbers, 2017; Ernst & Bülthoff, 2004). If the brain is able to integrate many bodily cues simultaneously to create a more accurate and quicker accessible representation, it hints at neural dynamics shaped by the body in a nontrivial manner.

Ecology of representation also corroborates the claim of spatial processing being grounded in bodily action. Sensory substitution or augmentation devices, the usability of which relies on lawful sensorimotor relations, have been shown to have significant effects on both spatial perception and cognition (Bach-y-Rita, 1972; Lenay et al., 1997; Nagel et al., 2005; Schumann & O’Regan, 2017). Spatial judgements improve in accuracy for participants wearing a belt that provides constant sensory information about north. However, the participants’ spatial reasoning processes seem to rely less on allocentric representational structures (Kaspar, König, Schwandt & König, 2014; König et al., 2016). Thus, representations might only be built if necessary to guide efficient action.

Recent results, both theoretical and experimental provide further evidence for a representational structure which is intimately linked to bodily action. Bicanski and Burgess (2018), for example, developed a model for spatial memory and imagery which features pseudo motor signals as an important factor. Bonner and Epstein (2017) reveal a bottom-up mechanism involving the visual cortex for perceiving potential paths for movement in one’s immediate surroundings.

We want to experimentally investigate the effect of bodily action on allocentric representational structure using behavioral measures. Large-scale studies of spatial cognition have mostly been asking participants to estimate distances or judge directions. Participants that point from landmark A to landmark B, however, already believe to have found the shortest distance between both landmarks (at least implicitly). The shortest distance implies a certain form of movement. It implies sensorimotor relations experienced during straight line travel. Therefore, these measures cannot be considered independent with respect to their relation to bodily action. We need to consider allocentric representations that clearly differ in their relation to action. A second hurdle to overcome is that, in general, the distinction between egocentric and allocentric is not clear-cut, as described above.

## Pilot Study

Both hurdles have been largely overcome in a recent pilot study, the precursor to our current project. The “Osnabrück study” inquired into the character of allocentric spatial knowledge which participants had acquired while living in a real town (König, Goeke, Meilinger, & König, 2019). As outlined above a town is especially well suited to investigate the relationship of embodiment and allocentric representations because it is a large-scale environment made up of visually separated vista spaces.

In the Osnabrück study the researchers exclusively investigated judgement of direction. For this, three tasks were set up. Two tasks were of foremost importance because they involve spatial concepts related to largely independent forms of bodily movement. In the Absolute task participants judged the orientation of a housefront relative to cardinal north. In the Relative task participants judged the orientation of two houses relative to each other. The researchers argued that the spatial concept of cardinal direction is categorically different from the spatial concept of house orientation concerning their relation to embodied action. While house orientation affords local action, like rotation, north informs global action, like long-distance travel.

The researchers additionally set up the Pointing task in which participants had to judge the shortest direction from one house to another. Pointing from landmark to landmark is most familiar to the participant and a standard measure throughout literature on spatial cognition. Similar to knowledge about cardinal direction, it also enables the participant to move globally on a straight line. It is, however, the only one of the three tasks that requires knowledge of relative location to be solved. Since its familiarity might additionally confound embodiment effects, the focus of analysis in the Osnabrück study was the comparison of Absolute and Relative tasks.

As discussed earlier, we generally need more time to plan travel along the shortest route in a large environment than to identify and pick-up an object in front of us. The difference in time until bodily action makes it interesting to compare task performance under time pressure to performance when the participant is free to reason. Consequently, in the Osnabrück study the spatial tasks were implemented for both time conditions.

The experimenters found that, under time pressure, participants were better at judging the angle between two housefronts, than judging the angle between house and cardinal direction of north. The analysis also revealed that without time pressure the relation reversed. When participants were allowed to reason indefinitely they excelled at the Absolute task. The interaction indicates two different forms of allocentric spatial representations. A possible explanation for the two forms could be the distinct actions afforded by the spatial concepts of house orientation and north, respectively.

## Present Study

One of the current study’s goals was to reproduce the results of the Osnabrück study in a more controlled environment. Consequently, again, we needed to find an experimental environment of a size and structure sufficient to ensure minimal egocentric influence on task performance.

In the last two decades virtual reality (VR) has become widely used among researchers across disciplines (Bohil, Alicea, & Biocca, 2011, Parsons, Gaggioli & Riva, 2017), also in spatial cognition (Hardless, Meilinger, & Mallot, 2015). Modern equipment combines stereoscopic 3D vision and largely unrestrained, full-body action possibilities with considerably low setup costs. Another advantage of a VR setup is that it allows access to coordinates of bodily movement in real time. Furthermore, VR allows for a feeling of “presence” of participants in the virtual environment, a reason why it is also used in research on emotion and therapy (Diemer, Alpers, Peperkorn, Shiban, & Mühlberger, 2015; Price & Anderson, 2007). Taking all of the above into consideration, we decided to repeat the pilot study by letting participants explore a complete city in VR. Our custom-made VR city contains roughly 200 houses spanning about 450 by 500 meters, which allows for extensive embodied exploration.

The main goal of the present study was to reproduce evidence for the two distinct forms of allocentric encodings found in the pilot study. As mentioned above, these distinct representations seem to result from the respective relation of the represented spatial concept to embodied action. We, thus, sought to not only reproduce but also expand the findings by investigating spatial representations acquired through different means of embodied exploration. Consequently, we set up three experimental embodiment conditions. Participants in each condition navigate through the same town, only the bodily action possibilities differ. After city exploration participants complete the same three spatial tasks with and without time pressure as in the pilot study. To observe the development of the spatial encodings we employ a longitudinal in-between design. Exploration in each embodiment condition is repeated three times. In summary, the setup allows us to compare the performances in three spatial tasks, times two decision-time conditions, times three exploration sessions, for each of the three modes of embodiment, respectively.

### VR Embodiment: Hypothesis One

In the VR condition participants explore the city sitting on a swivel chair, controlling movement speed with a handheld controller. Participants change direction through head movements, which are unrestrained, as are torso rotations around their longitudinal axis.

The VR embodiment condition was mainly set up to reproduce, or call into question, the findings of the Osnabrück study. Our hypothesis (H1) for the results after VR exploration reads as follows: Performance of Absolute and Relative tasks interact with time pressure. With unlimited decision time judging house alignment to north is more accurate than judging relative orientation of two houses and vice versa under time pressure. Houses can become decision points that afford local action involving rotation. Access of cardinal direction allows the navigator to move globally along one direction. Thus, finding the purported interaction after VR exploration suggests two forms of allocentric representations, which develop because they involve spatial concepts that allow for different bodily actions.

Compared to the other two exploration conditions we will introduce below, VR most closely resembles embodied navigation without augmenting devices (e.g., a map or smartphone). Thus, the VR results might also shed light on the general development of allocentric knowledge in large environments. While we have no concrete hypothesis, we are interested to see whether performance in the separate spatial tasks develops in parallel or shows interactions with an increasing number of sessions.

### VR with Belt Embodiment: Hypothesis Two

The VR condition with belt resembles the VR condition except that the participant is additionally equipped with a sensory augmentation device. The device is a belt worn around the waist with equally spaced vibrators of which always only the one closest to north is active. That means, if participants rotate clockwise on the swivel chair, the vibration switches to an element in the counter-clockwise rotation direction such that the location of vibration on the body remains closest to north.

Participants in the present study were not trained with the belt prior to VR city exploration and, thus, accustoming effects might confound results. Nevertheless, our hypothesis (H2) for the results after VR with belt exploration reads as follows: Performance in the Relative task is better than in the Absolute task. We assume that the belt reduces the ecological value of building a representation of north since the belt is delivering this information online. Consequently, the representation of north is lacking in the absence of the belt’s signal. Reasoning processes involving representations of the angle between house fronts, on the other hand, will not be negatively affected by the sensory augmentation device since the belt’s information is not directly helpful for this relative task.

### Map Embodiment: Hypothesis Three

For the map condition we created an interactive city map. Participants sit on an immobile chair in front of a setup of computer monitors. The monitor screens together display one large, birds-eye-view map of the VR city, with north being on top. Using a mouse cursor, participants can click on an individual house on the map, then a picture appears of the housefront view. In this setup, exploration of the VR city involves only a minimum of bodily movements.

Our hypothesis (H3) for the results after map exploration reads as follows: Performance in the Absolute task is better than in the Relative task. The map follows a north-up layout. The participant’s upright body is aligned with that north-up axis of the map for the entire exploration period. We, therefore, expect the structure of the acquired representation to favor performance in the Absolute task in both time conditions.

### General Expectations: Hypotheses Four to Six

We set out to design a study which incorporates three different embodiment conditions each allowing us to answer a different question, i.e. hypotheses one (H1) to three (H3). Consequently, we have no explicit hypotheses purporting effects in between those three conditions. Indeed, comparing hypotheses in-between different exploration conditions can come with a reduction in power due to possible in-between factors (e.g. getting accustomed to the belt might lower performance in all tasks) which are not the primary concern of this study. We do, however, expect three results independent of the means of embodiment during exploration. First (H4), we do expect to find better performance in Pointing than in the Relative and Absolute tasks. Pointing from one location to another represents direct knowledge about the (theoretically) shortest path to a goal. This kind of knowledge seems to be most important for navigation. In addition, of the three spatial tasks the Pointing task is probably by far the most familiar to our participants.

We already mentioned that a considerable part of people are poor spatial learners. Despite the apparent difficulty in generating largescale survey knowledge however, we nevertheless expect the following hypothesis (H5) to hold for the results after exploration in any of the three embodiment conditions: Performance increases with session. The intervals between sessions range from one to seven days. Consequently, even for participants which are not adept at spatial navigation and acquired little knowledge during the first exploration, we believe long-term changes in neuronal structure and activity will facilitate learning during subsequent sessions.

The final hypothesis (H6) we believe to hold for the task performance results after exploration in all of the three embodiment conditions reads as follows: Performance under time pressure is worse than without. Egocentric representations have a structure that allows quick action, both mental and bodily. Allocentric representations require more time to maximize accuracy of spatial judgements.

## Method

### Participants

Overall, 259 young and healthy adults took part in our study. They were randomly assigned to three differently embodied spatial exploration groups: virtual reality (VR), VR with sensory augmentation belt and map. We had to discard a total of 33 participants (8 VR, 18 VR with belt, 7 map) due to physiological issues like motion sickness in VR, as well as technical problems with data collection. The remaining 226 participants were distributed as follows: In the VR condition without belt we counted 82 participants (51 female; mean age 23 ± 3 years standard deviation). Exploring the VR city while wearing a sensory augmentation belt were 70 participants (40 female; age 23 ± 4). A total of 74 participants (50 female; age 24 ± 4) only explored a map of the city.

These numbers were informed by a power analysis based on the results of the pilot study mentioned in the introduction (König, et al., 2019). We calculated a sample size of about 60 participants to find the smallest effect in between two embodiment groups with 80% probability (G*Power (Faul, Erdfelder, Buchner & Lang, 2009): Two groups, one tailed, planned-power 80%, Cohen’s d=0.46 (μ1=19.4, μ2=18, σ=3)). As a sidenote, while we did not have explicit hypotheses comparing embodiment conditions, we still believed the power reasonable in order not to miss effects during exploratory data analysis.

The number of participants necessary to only miss an effect in 20% of experiments was performed for one experimental session. Initially, we had planned one session per participant per embodiment condition. Unexpectedly, performance remained close to chance in most tasks across conditions. Considering constraints on time and resources we settled for a design with three sessions for each experimental group.

Of the VR participants 28 (16 female; age 23 ± 3) took part in the longitudinal study. Of these 22 showed complete data sets in each of the three sessions. The remaining six had to discard one or two sessions each due to technical error. For example, a participant’s complete data set of the first session in the VR embodiment condition had to be discarded but her data was recorded correctly in the second and third session. We included all 28 longitudinal VR participants as well as the 54 VR participants who only explored the city once, for statistical analysis.

For the map condition again 28 participants (15 female; age 24 ± 3) took part in the longitudinal study. Here 26 had complete data sets after three sessions. We included all 28 longitudinal as well as 46 single-session participants for analysis. For VR with belt 27 participants (13 female; age 24 ± 4) returned for multiple experimental sessions. We found 23 of these to have complete data sets. All 27 longitudinal as well as 43 single-session participants were included in the analysis. Table 1 gives an overview of the total number of participants in each experimental session. The detailed distribution of participants and data is shown in the Appendix A.

**Table 1.** Number of participants per exploration condition per session.

| # Participants | First Session | Second | Third |
| --- | --- | --- | --- |
| VR | 82 | 28 | 28 |
| Map | 74 | 28 | 28 |
| VR with belt | 70 | 27 | 27 |

The participants could choose between either monetary compensation (9 Euros per hour) or ‘‘participant hours,’’ which are a requirement in most students’ study programs. All participants gave written informed consent. Also, only healthy participants (questionnaire data) were admitted for the experiment. The study was approved by the ethics committee of the Osnabrück University.

### Virtual Reality City

We built a VR city using Unity (version 5.6.3 and 2017.2.0), a software development environment to build video games. We named the city Seahaven. It makes up an island spanning 450 times 500 m^2^. Seahaven contains 213 houses ranging from wooden huts, garages and supermarkets, to villas, churches and office buildings.

To prevent participants from relying on simple navigation strategies Seahaven does not contain a building which is visible from all parts of town. In addition, there are no specific city districts, and the street system does not follow an ordered grid (i.e., “Manhattan style”). Finally, the angle between housefronts and north is roughly equally distributed in steps of 30°.

The VR environment includes a moving sun to provide natural lighting conditions and an indication of the direction of north. The sky over Seahaven is always cloudless, so the sun is clearly visible as well as its movement from sunrise to sunset (East to West) during the course of the day. Each participant experienced one such VR day in her 30-minute exploration period in the city. The only available cues to infer cardinal direction in Seahaven, that is most importantly the cardinal direction of north, are the sun’s position as well as the shadows cast by objects in the city. Figure 1 depicts the city from different perspectives.

**Figure 1.**
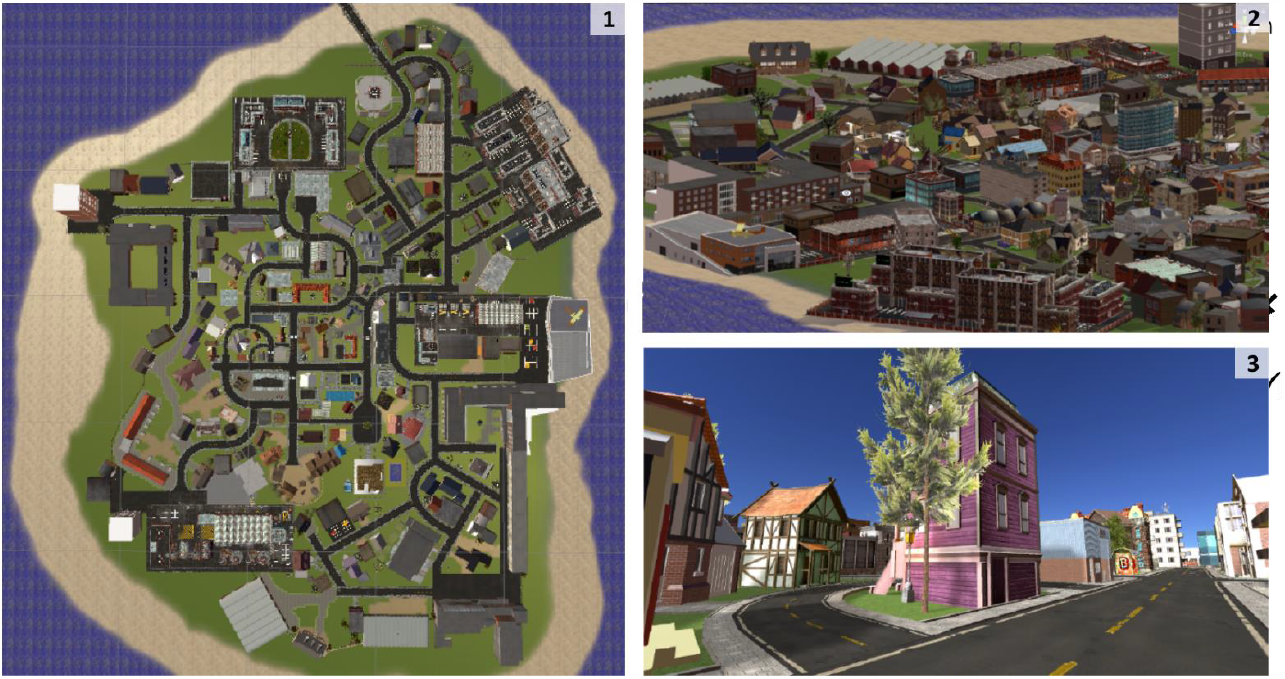
Virtual Reality City Seahaven. The figure depicts the VR city from different points of view. *1*.*1* Seahaven from a bird’s-eye perspective. North is up. *1*.*2* Panoramic view of the eastern part of the city. *1*.*3* 2D example of how a VR participant experiences Seahaven.

### Tasks

The task design is very similar to the setup used in the Osnabrück pilot study (König et al., 2019). Each participant is tested on three judgement-of-relative-direction tasks, once with and once without time pressure. The tasks follow a two-alternative-forced-choice (2AFC) design with 36 trials for each of the three kinds of spatial relations with each time condition. That makes for six task blocks each holding 36 trials (altogether 216 trials). The six task blocks were presented in random order. Trials per block were also randomized anew for each experimental session.

#### Absolute Task

In the Absolute task we assessed the participants’ ability to estimate the orientation of single houses in relation to north. In a 2AFC setup the participant was presented two pictures of the same house. Both pictures were overlaid with an arrow inside an ellipse. Only one of these arrows pointed to the cardinal north (as defined inside the VR environment) direction. The other arrow pointed in a direction that diverged from north by some amount between 30° and 330° (in steps of 30°). The pictures were presented at the same time and one above the other on separate screens (details on task setup below). The participant had to choose the picture with the arrow pointing towards north. She could choose the upper image or the lower image respectively by pressing the upper or lower button on a response box. Figure 2 depicts the task schematically.

**Figure 2.**
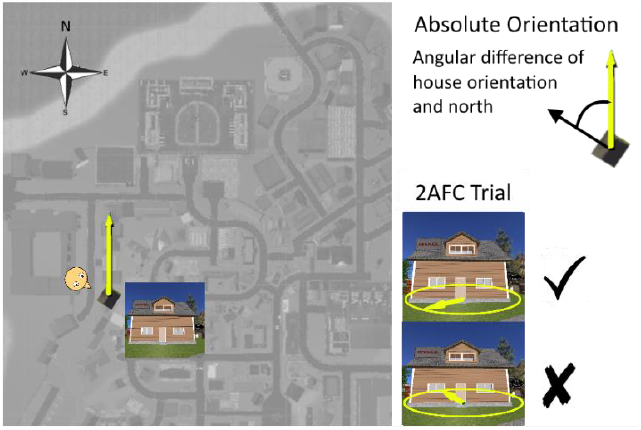
Absolute Task. In an Absolute task trial the participant had to choose the picture on which the arrow correctly pointed in the cardinal direction of north. Not depicted are the grey screens shown in between Absolute task trials. We presented 36 trials which had to be answered in three seconds and 36 trials without decision time limit.

Altogether, 36 2AFC trials with five second breaks (gray screens) in between each trial were presented sequentially in a block. There were two Absolute task blocks. In one block the participant had unlimited decision time. In another block the decision had to be made within three seconds, amounting to a total of 72 Absolute task trials.

#### Relative Task

In the Relative task, we assessed the participant’s ability to estimate the orientation of houses in relation to each other. We employed a 2AFC setup again, similar to the setup for the Absolute task but for the Relative task the participant saw three houses per trial. First the lower and upper screen both showed the same house (without an arrow). Then the upper and lower screen both showed two different houses. These are called target houses. One of these target houses faced in the same direction as the prime house. The other house of the target-house-pair faced in a direction that diverged from the prime house’s orientation by some amount between 30° and 330° (in steps of 30°). The participant’s task was to select the target house with the same orientation as the prime house. As in the Absolute task the participant chose for the upper image by pressing the upper button and for the lower image by pressing the lower button on the response box. Figure 3 depicts the task schematically.

**Figure 3.**
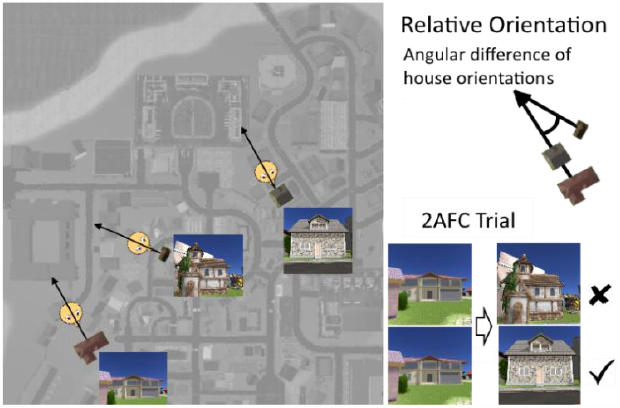
Relative Task. In a Relative task trial the participant first saw one house. She then saw two different houses and had to decide which of these was oriented in the same direction as the first house. We presented 36 trials which had to be answered in three seconds and 36 trials without decision time limit.

We presented 36 2AFC trials, each trial consisting of the prime house being shown for five seconds and then followed by the target houses, sequentially in a block. There were two Relative task blocks. In one block the participant had unlimited time to decide for the correct target. In another block the decision had to be made within three seconds, amounting to a total of 72 Relative task trials.

#### Pointing Task

In the Pointing task, we assessed the participants’ ability to estimate the orientation of a house relative to the direction towards another house. We again employed a 2AFC setup, similar to the setup for the Absolute and the Relative task. Also similar to the Relative task, each trial started with the upper and lower screen both showing the same house (no arrow). This prime house which was shown for five seconds. Then, as in the Absolute task, the participant was presented two pictures of the same house. These pictures of the one target house were both overlaid with an arrow inside an ellipsoid. One of these arrows pointed from the target towards the prime house. The other arrow pointed in a direction that diverged by some amount between 30° and 330° (in steps of 30°). The participants’ task was to select the target house picture with the arrow that pointed towards the prime house. As in the Absolute and the Relative task the participant chose for the upper image by pressing the upper button and for the lower image by pressing the lower button on the response box. Figure 4 depicts the task schematically.

**Figure 4.**
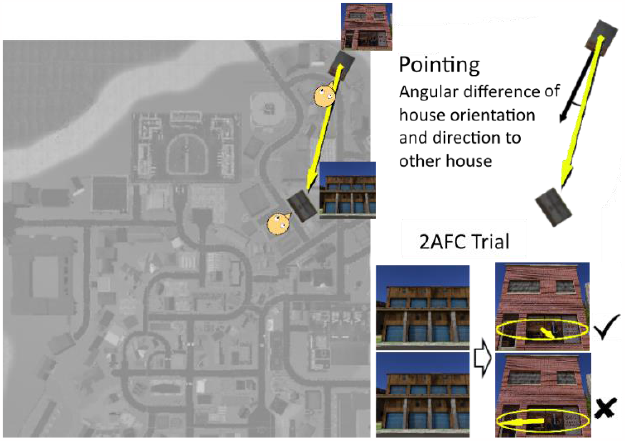
Pointing task. In a Pointing task trial the participant first saw one house. She then saw another house with two different arrows. The participant had to choose the arrow which pointed to the first house. We presented 36 trials which had to be answered in three seconds and 36 trials without decision time limit.

We presented each participant with 36 2AFC trials. Each trial consisted of the prime house being shown for five seconds, followed by the target house with a different arrow on the upper and the lower screen. There were two Pointing task blocks. In one 36 trial block the participant had unlimited time to decide for the correctly pointing arrow. In another block the decision had to be made within three seconds, amounting to a total of 72 Pointing task trials.

### Stimuli Preparation

The set of stimuli was entirely made up of screenshots of houses in the VR city. The screenshots were taken from the minimal distance each house could be wholly captured, with little else in the image that could serve as distractions. The pictures were taken during the course of a VR day. Since the sky in Seahaven is always cloudless each picture had good lighting conditions. Most importantly each screenshot was taken facing the building’s front. Therefore, the orientation of each house in the picture was towards the participant sitting in front of the monitors.

Most of the screenshots were overlaid with 3D stylized arrows lying inside an ellipse, similar to a compass. The ellipse and arrows were chosen bright green to increase salience and together took up the lower third of the picture (Ellipse and arrow in Figures 2 to 4 were slightly modified for better readability). Each picture had a resolution of 2160x1920 pixels.

So as not to create any bias for a certain direction, we wanted the orientation of stimuli used in the tasks to be equally distributed. Additionally, if the correct and wrong options in a 2AFC trial differ in their orientation by only 30°, it might be harder to judge correctly than if they are 60° apart. Consequently, we wanted the distribution of differences between choice options to be identical across all tasks. To not exhaust the participants too much after the already demanding exploration of the city we decided for 36 trials per task and time condition. Since Seahaven only contains 213 houses we need to allow houses to occur more than once across the three spatial tasks. We decided that within each of the three spatial tasks the stimuli which appeared in the condition with decision time limit should not appear again when the participant had unlimited decision time and vice versa. Only the prime houses, which are the first houses that appear before the actual decision of the participant in each trial in the Relative and Pointing task, can appear multiple times. We decided to use all together a set of 18 prime houses. These were chosen due to their high familiarity ranking in accordance with results from a preliminary study. They also had to fulfill the criteria of being distributed at different locations in the city and fit the constraints on equal orientation described above. The final set of stimuli was picked through an algorithm fulfilling the above constraints and thereby ensuring homogeneous stimuli properties for each task.

We used the algorithm to create one stimulus set which was employed after each exploration independent of embodiment condition. Consequently, within each task block all stimuli are identical across participants. The order of trials during each block as well as the screen on which the trial stimuli are presented (upper or lower screen) are newly randomized for each participant. The same holds for the order of the six task blocks themselves.

### Procedure

We partitioned each experimental session into three main parts. First the participant was introduced to the tasks. Then she explored the city. Finally, the participant completed the tasks. All three sessions over all three exploration conditions shared the same introduction and the same task blocks (task order was randomized). The conditions only differed in the embodied city exploration possibilities. A graphical overview is given in Figure 5.

**Figure 5.**
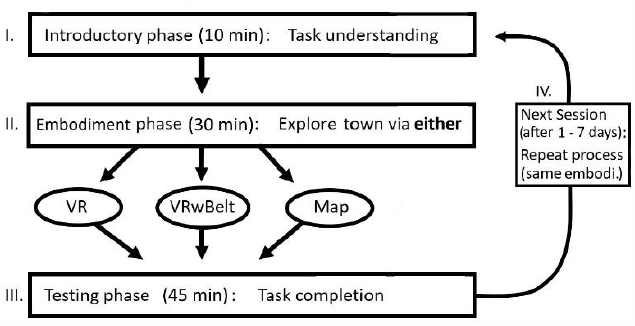
Overview of the complete study design. Each experimental session started with an introductory phase. Here the participant was (re-)introduced to the tasks and completed a time pressure response training. The introduction was followed by the embodiment phase. Here the participant explored the city for 30 minutes using one of three different bodily means for exploration. After the exploration phase the participant had to complete the six task blocks. Task completion was followed by questionnaires on navigational behavior and aptitude. Some participants returned for multiple sessions, always in one and the same bodily exploration condition.

#### Task Introduction

The introduction consisted of three parts. It started with the experimenter explaining the spatial tasks, in which one example picture for each of the spatial tasks was shown to the participant. These pictures were taken from houses of the city of Osnabrück. After this explanation the participant sat at about 80 cm distance in front of a six-screen monitor setup in a 2x3 arrangement. Only the middle column of monitors was turned on. Each of the two screens showed an arrow inside an ellipse. The participant had to choose the arrow which pointed more upwards. She chose for the arrow on the upper screen by pressing the upper button labelled “U” and for the arrow on the lower screen by pressing the lower button labelled “D” on a four-button response box (Black Box ToolKit USB response pad). The box was placed approximately 30 cm in front of her and left and right button were unmarked and had no function. If the participant did not decide after three seconds, the screens turned red. If she chose the wrong arrow the screens turned blue. Upon providing the correct answer within three seconds the screens turned green. Arrow pairs appeared sequentially until the participant correctly decided in 48 out of 50 trials. Following this response training the participant did one trial for each of the three spatial tasks, once with time limit and once without. For these six test trials, we again used houses from the city of Osnabrück. In these trials the participant, only received feedback if she did not answer in the time limit condition. Then the screens turned red. The complete introduction phase lasted approximately 10 minutes.

#### Embodied Exploration

Participants were randomly assigned into one of three groups, each defined through its specific means of embodiment of exploration. If the participant returned for repeated measurements, she stayed in her initial embodiment group.

##### VR Embodiment

The participant sat upright on a swivel chair wearing the VR headmounted display and headphones. The headphones played a sound loop of ocean surf to prevent external environmental sounds from distracting or confusing the participant. The participant also held a VR controller in one hand.

Each participant got accustomed to the VR experience on a small island set up only for training purposes. Touching a disc on the controller with her thumb the participant was able to regulate both speed (max. 3.3 m/s) and direction with which she moved across the island. The participant was specifically instructed not to change direction using the handheld controller. To decrease the risk of motion sickness, change of movement direction should happen exclusively by rotating the chair.

We then proceeded with calibrating the eye tracker implemented in the VR goggles. During calibration the island disappeared. The participant had to focus her gaze onto ten sequentially appearing targets spread over the VR goggles’ viewing field. This process was repeated for validation. Calibration and validation were repeated until gaze location could be detected with an error smaller 2°. The participant was then teleported to the VR town. A session always started at the same location, central in the city and facing east.

We instructed each participant that she could freely explore the city for half an hour and that this would amount to one virtual day, that is from virtual morning to evening. Every five minutes the validation process was repeated, but considerably shorter, with only one validation point. Only when the gaze location error became larger than 2° was the complete ten-point calibration repeated. At the end of the half-hour exploration time the participant was asked to turn towards north. Note that throughout exploration of the city, north could only be inferred from the sun’s position and the direction of shadows. The complete VR exploration procedure took approximately 45 minutes. The setup is depicted in Figures 6.1 and 6.2.

**Figure 6.**
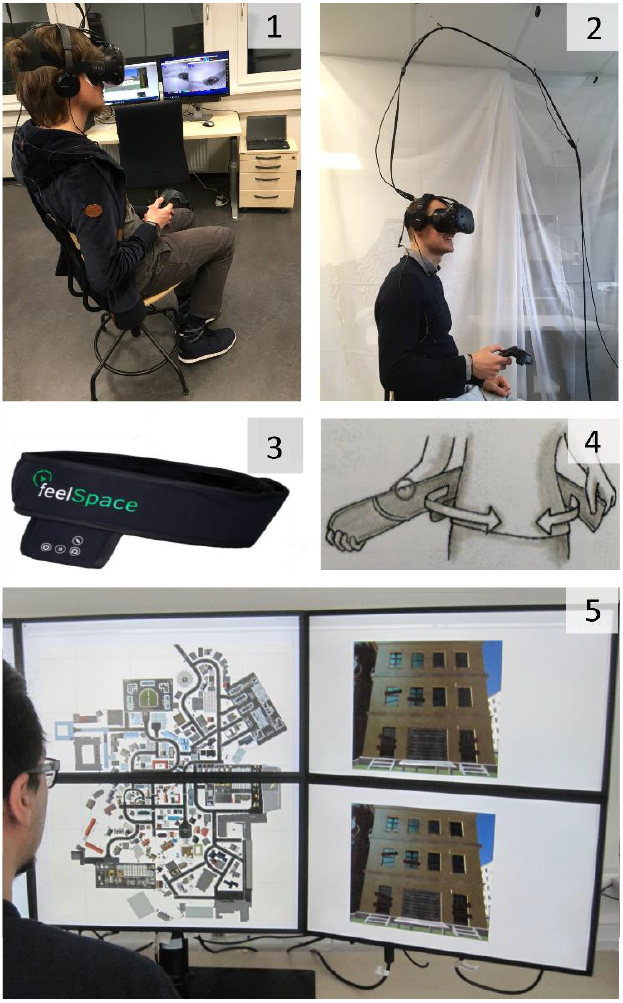
Experimental Setup. *6.1* Participant with head-mounted VR display, headphones and VR controller during an experimental session. He uses the controller to regulate his movement speed and rotates on the chair to change movement direction in the VR environment. In the background one sees the experimenter station. *6.2* Experimenter with head-mounted display, headphones and controller. The cable management allows for free rotation. *6.3* The sensory augmentation belt. Visible is the feelSpace logo and the control unit (small rectangular bag with four symbols). Not visible are the 16 evenly spaced vibration elements which are covered by the outer layer of cloth. During the experiment, only the northernmost (VR north) vibration element is active. Image provided courtesy of feelSpace GmbH. *6.4* The belt is strapped around the waist. The participant can then walk around or (for the experiment) sit down on the VR swivel chair (depicted in Figure 6.1). *6.5* Experimenter sitting in front of the interactive map setup as used during a regular experimental session. With the mouse (not visible) he clicked on a house in the map to see a picture of the house.

##### VR with Belt Embodiment

In this group the participant wore the sensory augmentation belt while exploring the city in VR. Before exploring the VR environment, the participant was introduced to the belt. She then walked around the lab with the activated belt vibrating north (real world direction) until she felt accustomed. The belt was then turned off and the participant introduced to the VR setup as described above.

Note that this is a very short introduction period, only sufficient to get familiar with the basic principle of the sensory augmentation device. Prior to this experiment, our participants did not have any experience with the belt. Different from past studies, they are not trained belt wearers (Nagel et al., 2005; Kaspar et al., 2014; König et al., 2016).

After the participant had been teleported into the VR town (to the same location as in the VR experiment), she was asked to turn towards a specific building. This way the participant directly faced in the direction of VR north. We then activated the belt again and asked the participant to hold her arm out in the direction of the vibration. We changed the belt’s vibration direction until it matched the VR city’s north, that is until the participant pointed directly forward. Each belt wearing participant then freely explored the city for half an hour, from virtual morning to evening.

As in the VR condition, we performed a validation on eye tracking accuracy every five minutes. Likewise, at the end of the exploration time the participant was asked to turn north. The complete VR with belt procedure took approximately 55 minutes. The belt is depicted in Figure 6.3 and 6.4.

##### Map Embodiment

For the exploration of the city map we used the same six-screen monitor setup in a 2x3 arrangement as during the introductory phase. The left column of monitors was turned off. We used a 2D map of the VR city (see Figure 1.1, only houses and streets with white background), which stretched across the middle column of monitors. The participant sat about 80 cm away from the monitors and could use a mouse to move a cursor across the city map.

When the cursor hovered over a house, the house showed a red dot (only for 193 out of all 213 houses). After clicking on a house with a red dot, the right column of monitors showed, on both screens, the same frontal view of the house. The position of the dot on the bird’s-eye-view house outline on the map indicated from which side the frontal-view picture of the house was taken. Thus, we ensured that participants knew the orientation of houses.

After the participant was made familiar with the interactive map design, we instructed her that she could freely explore the city for half an hour. After 15 minutes of exploration time had passed, we reminded the participant that 15 minutes were left. The complete map exploration procedure took 30 minutes. The setup is depicted in Figure 6.5.

#### Task completion

After the exploration period during which the participant became familiar with the spatial layout of the VR city, she did a series of spatial tasks with and without time pressure. The tasks were the same as introduced at the beginning of the experiment, but the spatial relations now asked for were among houses from the VR city.

The participant had to complete a block of 36 trials for the Absolute task with a time limit of three seconds for each trial. She also had to complete a block of 36 trials for the Absolute task with unlimited decision time for each trial. She did the same for the Relative task as well as the Pointing task. These six task blocks were presented in random order to each participant. Before each block a screen appeared explaining the task. As in the introductory phase, the participant sat approximately 80 cm away from the six-screen setup, and only the two middle screens were used. The task completion took around 45 minutes.

After the participant completed the tasks, she was asked to fill in the “Questionnaire of Spatial Strategies” asking about her own perceived everyday navigation strategies and skills (translated from German, “Fragebogen Räumlicher Strategien”, or FRS; Münzer & Hölscher, 2011). The complete experiment took approximately two hours.

### Materials

#### Task Setup

Each of the six screens had width 53cm times height 31cm and was used at a resolution of 1920x1080 pixels. The 2x3 setup was synchronized with Samsung SyncMaster supported by three NVIDIA Quadro K620 graphics cards. The PC processor was an Intel Xeon E5620 running at 2.4 GHz with 12 GB RAM.

#### Virtual Reality Setup

For our experimental setup we used the HTC Vive Virtual Reality System. We installed two base stations on the ceiling of the lab. These tracked the position of both the head-mounted display, which we will refer to as VR goggles, as well as the VR controller at a rate of 60 Hz. The VR goggles had an image-refresh rate of 90 Hz and a binocular field of view of about 110°. The device used two OLED panels, one per eye, each having a display resolution of 1080×1200 (technical specifications as provided by HTC).

Inside the VR goggles we installed the Vive Pro Binocular Add-on eye tracker from Pupil Labs (Kassner, Patera, & Bulling, 2014). This eye tracker has a gaze accuracy of 1° and updates at a maximal frequency of 120 Hz (technical specifications as provided by Pupil Labs). For performance reasons we only tracked the gaze location at a frequency of 30 Hz. For more details on how we implemented eye tracking in VR see Clay, König and König (2019).

The software interface linking the PC and the Vive VR System was the game development software Unity (version 5.6.3 and 2017.2.0). The PC processor was an Intel Xeon E5 running at 3.7 GHz with 16 GB RAM. The graphics card was a NVIDIA GeForce GTX TITAN X. During a typical experimental session in the VR city, the rate of frames sent from the PC to the VR goggles varied between 90 and 30 Hz. We used Sony MDR-ZX310B headphones.

#### Sensory Augmentation Belt

For our experimental setup we used the naviGürtel Beta’17.Q1 in sizes C, M and L from feelSpace. The sensory augmentation device provides directional information of magnetic north via vibration around the waist.

The belt’s control circuit board contains a compass module as well as 16 vibromotors uniformly distributed over its length. Each vibration unit covers an angle of 22.5°. The vibration frequency is 170 Hz. The belt is powered by a lithium-ion battery allowing for approximately eight hours of continuous usage. If the belt is used without an additional device (e.g. smartphone), it functions like a compass in that only the vibration unit that is closest to north vibrates.

The feelSpace belt can also be used in combination with a Bluetooth capable device, allowing users to recalibrate north. We used the belt in combination with the smartphone Samsung Galaxy J3 (2017), which has Bluetooth capabilities. The Android application Luftlinie beta then enabled us to gauge the belt’s control circuit onto the VR city’s internal north.

#### Spatial Strategy Questionnaire

The FRS questionnaire is a self-report measure to assess proclivity and ability of participants to use three main spatial navigation strategies (Münzer & Hölscher, 2011; Münzer, Fehringer & Kühl, 2016). The three strategies are differentiated in global-egocentric, survey-allocentric and cardinal-allocentric. The global-egocentric questions are about general navigational ability as well as knowledge of directions and knowledge of routes and usage thereof, it comprises ten items. The survey-allocentric factor includes items specifically asking about mental map formation and usage, it comprises seven items. The cardinal-allocentric scale finally only includes questions about the knowledge of cardinal directions, it comprises two items. Each question is a Likert item with a score ranging from 1 (“I disagree strongly.”) to 7 (“I agree strongly.”).

#### Data

The resulting data is available at https://osf.io/kyp83/.

### Method of Analysis

The experiment follows a longitudinal, within-subject design for each exploration group. Consequently, we decided for linear mixed-effect models with the participant as random effect grouping variable as our method of analysis.^1^

Note that our analysis includes all participants even though about 2/3 did not complete the repeated sessions. While we excluded some data due to technical error, the total imbalance is almost entirely due to a change in the original study design, which only called for one session instead of three sessions per participant. Session is therefore nested within subgroups of subjects: One subgroup only received one session and the other received three sessions. The experimental procedures were identical for each subgroup. Also, t-tests did not reveal significant differences between the one-session and three-session participants in their respective first sessions marginalized over tasks. Details of the comparisons including all single task conditions can be found in Appendix B. We thus decided to ignore subgroup as possible factor and treat the missing data as completely missing at random. Then, including all participants should result in the estimate closest to the population value (Ibrahim & Molenberghs, 2009; Rubin, 1976).

Former studies investigating spatial cognition abilities found that males reacted faster and performed better than females in a variety of spatial tasks (Masters & Sanders, 1993; Moffat, Hampson, & Hatzipantelis, 1998; Newhouse, Newhouse, & Astur, 2007). Preliminary analysis of the data collected in our study did not reveal evidence that gender differences affect our general claims. We therefore do not include gender as a factor in our models.

We do not include item (house stimulus) as a random effect since we specifically designed an algorithm to ensure homogeneous distribution of stimuli properties across tasks (see above). The house stimuli in each task are assumed to be an exhaustive sample of the underlying population, that is the town Seahaven, which makes item a fixed variable (Tukey & Green, 1960).

We differentiate in models for best data fit and models for plotting purposes. The best-fit models are selected following a stepdown procedure. First the maximum model is run. All possible fixed terms, that means also all possible interactions are included. Following Barr et al. (2013) but adjusting for sample size, we also include all these terms except interactions as random effects. Then a likelihood ratio test is run comparing the maximum model to a nested model with one less fixed term (random structure unchanged). The comparison is done for all fixed terms. Afterwards, the least likely coefficient is dropped from both the fixed and (given it is part of it) the random effect structure. This cleaning step leaves us with the maximum-1 model. The procedure is repeated until a maximum-k model is reached with only significant fixed effects. This is the best-fit model.

In order to use likelihood ratio tests, we employ a maximum likelihood fit algorithm (not restricted). Throughout the analysis, we fit a heterogeneous, but due to convergence issues, diagonal covariance matrix, i.e. possible covariance between random effects is not represented.^2^

The effects of a posteriori model selection on Type I and Type II errors are complex and no universal policy is agreed upon (Altman & Andersen, 1989; Heinze, Wallisch, & Dunkler, 2018; Murtaugh, 2009; Tibshirani, 1996). For mixed models, the situation becomes even more complicated (Müller, Scealy, & Welsh, 2013). To minimize potential model selection effects on alpha and beta errors, here we only interpret significant coefficients at odds with, or in favor of, our a priori hypotheses. We provide all model design matrices and results in Appendix D.

Model selection also makes the level at which to extract nonsignificant coefficients for plotting purposes somewhat arbitrary. If, for example, in the best-fit model the difference between some task conditions over sessions turns out not to be significant, do we depict the average as the best linear-fit line? To allow for qualitative trends to show in our plots, we decided to fit each plot line separately on the data of exclusively one of the six task conditions over sessions. For example, the Absolute task, three second decision time condition was fitted independently. This separate plot-fit approach is simple and systematic. The only exceptions were the plots depicting linear-fit lines where we marginalize over the three spatial tasks. Details are provided in Appendix C.

## Results

We define performance or accuracy in each task condition as the fraction of correct answers in percent. A performance of fifty percent corresponds to chance in the no-time-limit task condition. With time limit, if participants do not answer in the three seconds the trial is counted as wrong. Thus, here chance performance can be below fifty percent if the participant hesitates too long.

### VR Embodiment

Following hypothesis one (H1) we expect participants to perform better in the Relative than in the Absolute task under time pressure after having explored the city Seahaven in VR. Linear mixed-model (Appendix D, Model s-VR) likelihood comparison however does not provide evidence for such an effect. The best fit model does not differentiate between task performances. It only hints at an increase in mean performance over three sessions by two percent which, however, remains insignificant (coefficient session=2, standard error (SE)=1, confidence interval (CI)= -0.1, 4.2 (all in percent correct trials, that means performance); likelihood ratio statistics χ2(1)=3.4, p=0.065). For a qualitative comparison see Figure 7.5.

**Figure 7.**
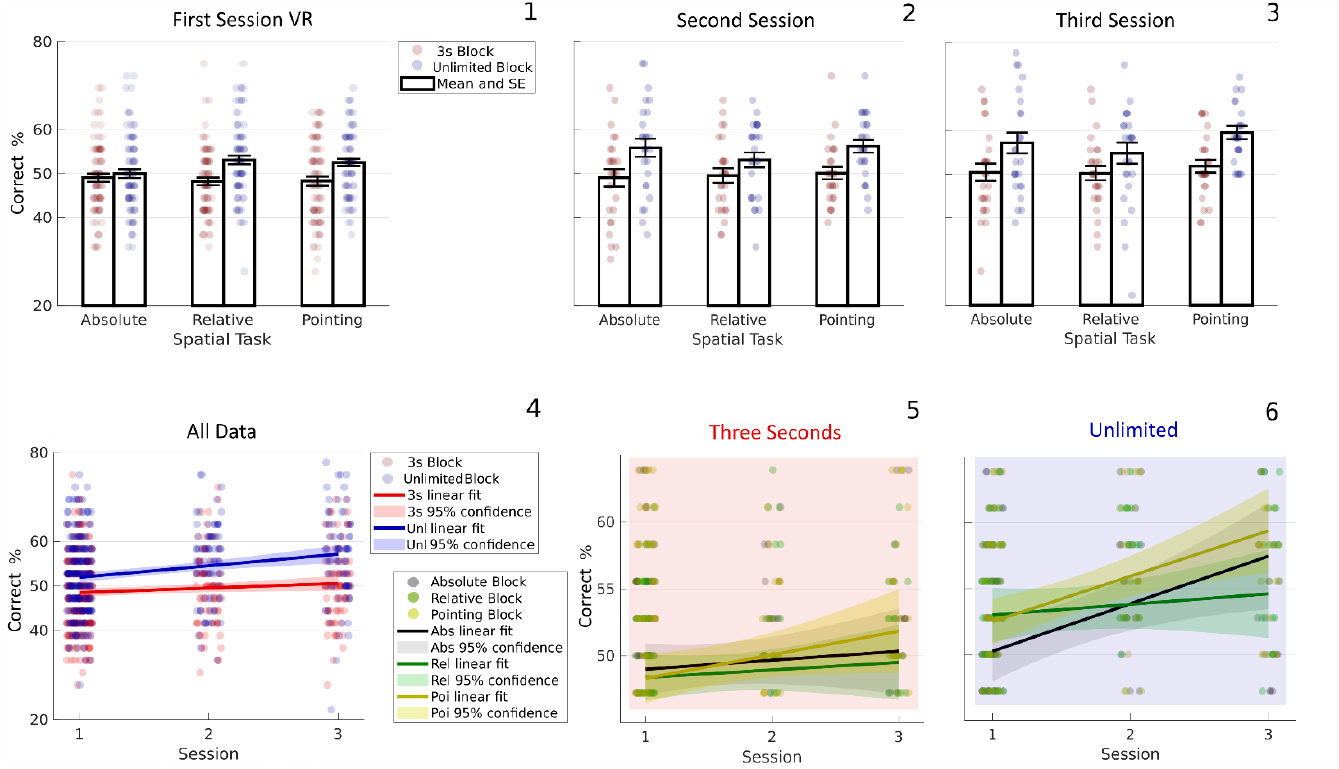
VR Embodiment. *7.1-3* (Black bars) Barplots of mean performance and standard error (SE) in all six trial blocks of VR participants in their first, second and third session, respectively. (Red dots) Single participant’s performance in block with three seconds decision time per trial. (Blue dots) Single participant’s performance in block with unlimited decision time. *7.4* Participants without time pressure are five percent more accurate than those with limited decision time (χ2(1)=41.6, p<0.001). Participants make four percent more accurate spatial judgements with increasing number of exploration session (χ2(1)=18, p<0.001). The increase in performance without time pressure becomes three percent stronger over the three exploration seesions (χ2(1)=4.5, p=0.035). (Red dots) Single participant’s performance in the three-second condition for each spatial task, respectively. (Blue dots) Single participant’s performance for unlimited decision time. (Red line) Linear mixed-effect penalized least squares fit with participant as grouping variable across all three-second blocks. (Blue line) Linear fit across all unlimited decision time blocks. (Red area) The 95% confidence interval for the three-second condition linear fit. That is, 95 of 100 intervals each from a different sample include the actual population fit. (Blue area) The 95% confidence interval for the unlimited condition. *7.5* With time pressure participants only show an insignificant performance increase of two percent over sessions (χ2(1)=3.4, p=0.065). (Black, Green, Yellow dots) Single participant’s performance in the Absolute, Relative and Pointing spatial task trial block, respectively, only for trials with time limit. (Black, Green, Yellow line) Linear fit across sessions in the condition with time limit, for Absolute, Relative and Pointing task, respectively. (Black, Green, Yellow area) The 95% confidence interval for the three-second condition linear fit for Absolute, Relative and Pointing, respectively. *7.6* Given unlimited decision time Pointing task performance is two percent above Absolute performance (χ2(1)=5.5, p=0.018). The difference between Absolute and Relative task performance reverses by six percent over the three sessions (χ2(1)=7.18, p=0.007). (Black, Green, Yellow dots) Single participant’s performance in the Absolute, Relative and Pointing spatial task trial block, respectively, only for trials without time pressure. (Black, Green, Yellow line) Linear fit across sessions in the free-reasoning condition, for Absolute, Relative and Pointing task, respectively. (Black, Green, Yellow area) The 95% confidence interval for the unlimited time condition linear fit for Absolute, Relative and Pointing, respectively.

A second result to be expected if H1 were to hold would be that, with unlimited decision time judging house alignment to north is more accurate than judging relative orientation of two houses. That is the opposite effect as expected with time pressure. The model fit (u-VR) does not corroborate this claim, at least not one-to-one. It rather seems that the difference in performance between Absolute and Relative task reverses with session. Absolute starts out worse than Relative but outperforms it with more sessions. The model reveals a significant change of performance difference from session one to session three of about six percent (interaction Abs. to Rel. with session= -5.6, SE=2.1, CI= -9.7, -1.5; χ2(1)=7.18, p=0.007). Extrapolating this interaction, we can surmise that H1 holds for large environments only for explorers with sufficient experience.

We do also find a main effect of task in the no-time-limit condition. The model fit (u-VR) shows that Pointing outperforms the Absolute task by two percent (Abs. to Poi. = 2.2, SE=0.9, CI=0.4, 4; χ2(1)=5.5, p=0.018). This result corroborates hypothesis four (H4) which simply claims that participants should perform best in the Pointing task. Figure 7.6 depicts the results qualitatively.

To investigate participants’ overall development in the three tasks across sessions, comparing also both time conditions, we fit a model (a-VR) that incorporates the complete VR data and time as an additional potential factor. Likelihood comparisons do not show any task to be significantly above overall task average. That means Pointing task performance is not higher than Absolute and Relative task performance in both decision time conditions. VR embodiment does not show conclusive support for H4. As purported by hypothesis five (H5) the model does reveal a performance increase by five percent for participants who answer without time pressure (time=5, SE=0.7, CI=1.9, 5.2; χ2(1)=41.6, p<0.001).

It also becomes evident that knowledge of spatial relations in our VR city accumulates over several visits as stated in hypothesis six (H6). We find an increase over the three sessions by about four percent. (session=3.6, SE=0.8, CI=1.9, 5.2; χ2(1)=18, p<0.001). The difference in performance from three-seconds to unlimited decision time increases over sessions by three percent as well (time with session=3.2, SE=1.5, CI=0.2, 6.1; χ2(1)=4.5, p=0.035). This interaction reveals that the accumulated spatial knowledge after three exploration sessions is mostly represented in a form that requires processing time to be accessed. The results are illustrated in Figure 7.4.

Visual comparison of task performances in each session which are detailed in Figures 7.1 to 7.3 does not give a coherent picture without model guidance. Performance is higher in the condition with no time pressure compared to the condition with a three second time limit. Performance is also higher in later experimental sessions compared to earlier experimental sessions. Interestingly, only the Pointing performance increases linearly with the number of sessions in both time conditions. When free to reason, participants show a large increase in performance in the Absolute task from the first to the second session. No such increase in performance is found for the Relative task. Under time pressure, the performance in both tasks stays at chance level. All model design matrixes and detailed results can be found in the Appendices C and D.

### VR with Belt Embodiment

As stated in hypothesis two (H2) we expect participants to perform better in the Relative spatial task asking for house-to-house orientations than in the Absolute task asking for house-to-north orientations if equipped with the north belt during the exploration period.

Performance under time pressure does not corroborate our claim. Linear mixed-model (Model s-VRwB) likelihood comparison show instead that both Absolute and Relative start out well below Pointing but draw even in the third session. The model makes evident that the difference in performance from Pointing to Absolute decreases by five percent over the three sessions (Poi. to Abs. with session=5.2, SE=1.9, CI=1.5, 9; χ2(1)=7.6, p=0.006). Similarly, Pointing to Relative decreases four percent over sessions (Poi. to Rel. with session=3.9, SE=1.9, CI=0.2, 7.6; χ2(1)=4.2, p=0.04). For a qualitative comparison see Figure 8.5. Extrapolating these results hints at both Absolute and Relative outperforming Pointing with more exploration time. The result implies that unexperienced belt wearers are not able to integrate the information the north belt provides into a quickly accessible representation. On the contrary, the belt rather seems to have led to structures misrepresenting allocentric spatial relations for unexperienced belt learners. At least when these representational structures are accessed under time pressure.

**Figure 8.**
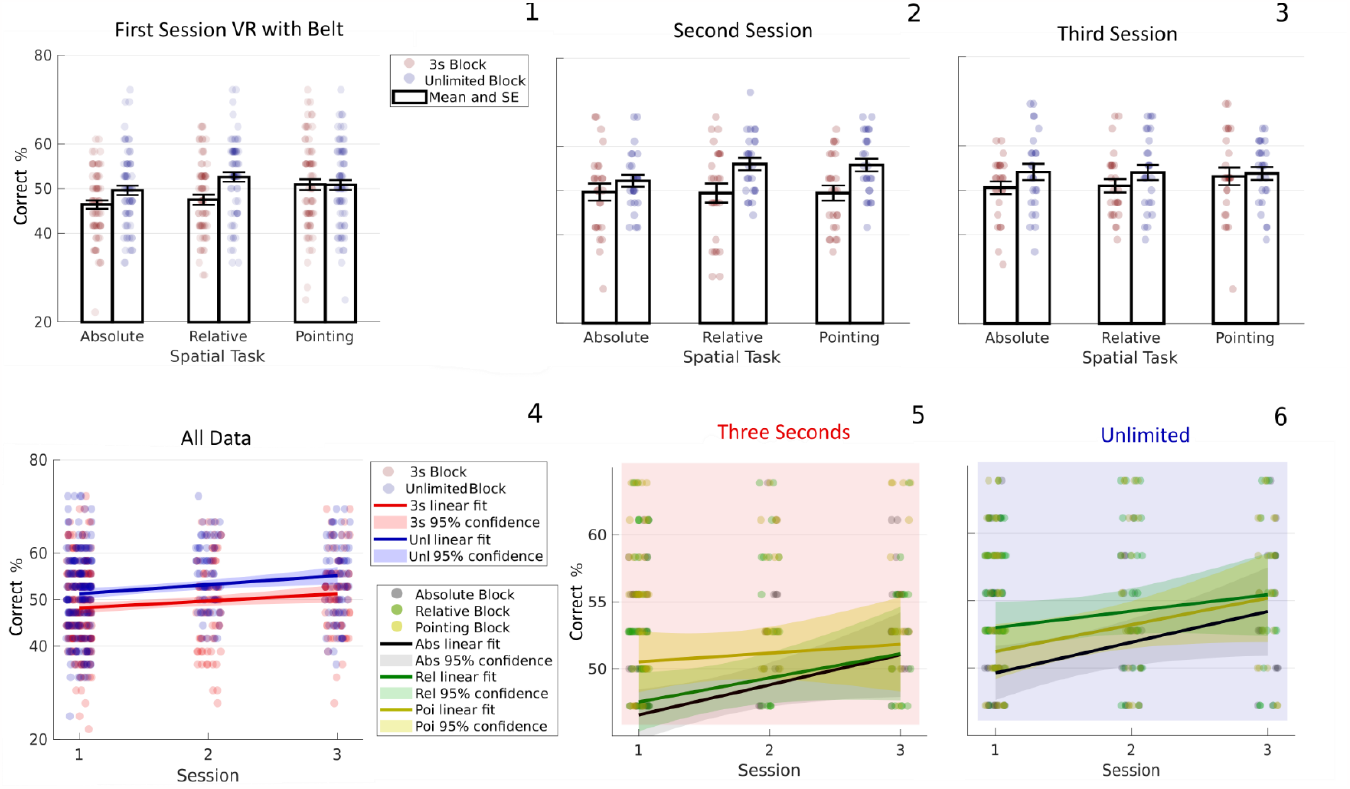
VR with Belt Embodiment. *8.1-3* (Black bars) Barplots of mean performance and standard error (SE) in all six trial blocks of VR with belt participants in their first, second and third session, respectively. (Red dots) Single participant’s performance in block with three seconds decision time per trial. (Blue dots) Single participant’s performance in block with unlimited decision time. *8.4* Correct spatial judgements increase by three percent both over repeated sessions (χ2(1)=17.23, p<0.001) and without time pressure (χ2(1)=12.4, p<0.001). Performance in the Pointing task is one percent better than overall task average (χ2(1)=7, p=0.008). (Red dots) Single participant’s performance in the three-second condition for each spatial task, respectively. (Blue dots) Single participant’s performance for unlimited decision time. (Red line) Linear mixed-effect penalized least squares fit with participant as grouping variable across all three-second blocks. (Blue line) Linear fit across all unlimited decision time blocks. (Red area) The 95% confidence interval for the three-second condition linear fit. That is, 95 of 100 intervals each from a different sample include the actual population fit. (Blue area) The 95% confidence interval for the unlimited condition. *8.5* When participants have to decide under time pressure the difference from Pointing task to Absolute task performance decreases by five percent over the three exploration sessions (χ2(1)=7.6, p=0.006). Similarly for the difference from Pointing to Relative which decreases by four percent (χ2(1)=4.2, p=0.04). (Black, Green, Yellow dots) Single participant’s performance in the Absolute, Relative and Pointing spatial task trial block, respectively, only for trials with time limit. (Black, Green, Yellow line) Linear fit across sessions in the condition with time limit, for Absolute, Relative and Pointing task, respectively. (Black, Green, Yellow area) The 95% confidence interval for the three-second condition linear fit for Absolute, Relative and Pointing, respectively. *8.6* Given unlimited decision time belt participants perform two percent better in the Relative task than in the Absolute task (χ2(1)=4.6, p=0.03). (Black, Green, Yellow dots) Single participant’s performance in the Absolute, Relative and Pointing spatial task trial block, respectively, only for trials without time pressure. (Black, Green, Yellow line) Linear fit across sessions in the free-reasoning condition, for Absolute, Relative and Pointing task, respectively. (Black, Green, Yellow area) The 95% confidence interval for the unlimited time condition linear fit for Absolute, Relative and Pointing, respectively.

Without limit on reasoning time the participants’ decisions provide evidence for H2. The best model fit (u-VRwB) reveals that Relative outperforms Absolute by two percent across sessions (Abs. to Rel.=2, SE= 0.9, CI=0.2, 3.5; χ2(1)=4.6, p=0.03). Figure 8.6 depicts the results qualitatively.

To investigate participants’ overall development in the three tasks across sessions, comparing also both time conditions, we fit a model (a-VRwB) that incorporated the complete VR with belt data and time as an additional potential factor. The model provides evidence for H4 because it shows Pointing to be significantly better than the task average (mean to Poi.=1.1, SE=0.4, CI=0.3 1.9; χ2(1)=7, p=0.008).

Furthermore, as in the other two conditions, we find evidence for H5 and H6. That is performance increases without time pressure (time=3.3, SE=0.9, CI=1.6, 5.1; χ2(1)=12.4, p<0.001) and over repeated sessions (session=3.4, SE=0.8, CI=1.8 5; χ2(1)=17.23, p<0.001). The results are illustrated in Figure 8.4.

Also, similar to the other two exploration conditions, visually assessing systematic differences in performance development from Figures 8.1 to 8.3 is barely possible. In the three-second spontaneous-decision condition participants perform worse than with reasoning time and performance increases with session. Unprecedented across exploration condition is that for decisions without time pressure, Absolute performance does not show a large increase from the first to the second session.

All model design matrixes and detailed results can be found in Appendices C and D.

### Map Embodiment

Hypothesis three (H3) predicts that the performance in the Absolute task will be better than in the Relative task after exploring the 2D city map. Linear mixed-model (Model s-Map) likelihood comparison, however, does not provide evidence for such an effect under time pressure. We instead find that participants perform about five percent better in the Relative task compared to Pointing (Poi. to Rel.=5.4, SE=1.3, CI=2.9, 7.8; χ2(1)=16.3, p<0.001). Participants also outperform Pointing in the Absolute task by four percent (Poi. to Abs.=3.9, SE=1.2, CI=1.6, 6.1; χ2(1)=10.9, p<0.001).

Note that these results are largely due to differences in long-term learning processes necessary to solve the spatial tasks. Such varying learning processes are evident as the difference in performance from Relative to Pointing increases by eight percent over the three sessions (Poi. to Rel. with session=8, SE=2.1, CI=4, 11.9; χ2(1)=14.9, p<0.001). Similarly, Absolute to Pointing increases by six percent over sessions (Poi. to Abs. with session=6.1, SE=2, CI=2.3, 10; χ2(1)=9.6, p=0.002). In statistical terms, these are two interactions of task and session. Importantly, these interactions were both unexpected. For a qualitative comparison see Figure 9.5.

**Figure 9.**
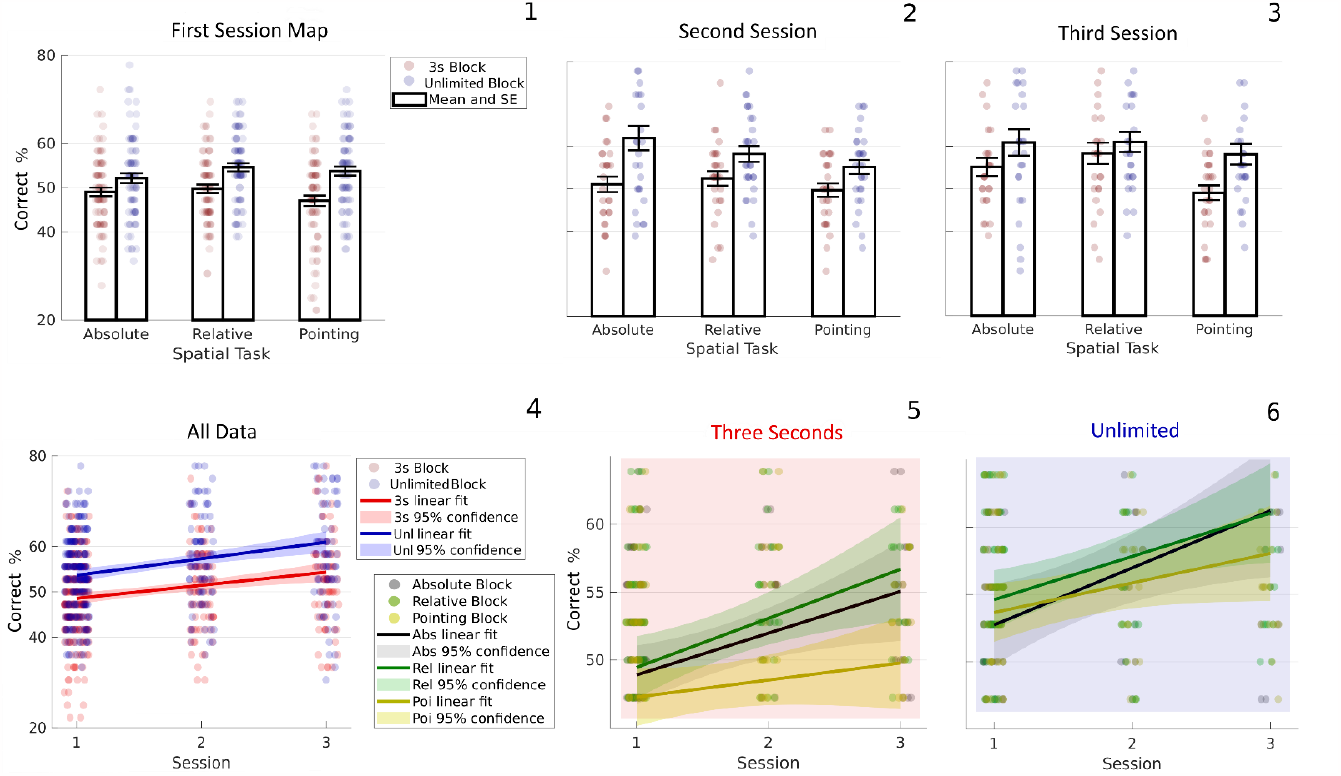
Map Embodiment. *9.1-3* (Black bars) Barplots of mean performance and standard error (SE) in all six trial blocks of map participants in their first, second and third session, respectively. (Red dots) Single participant’s performance in block with three seconds decision time per trial. (Blue dots) Single participant’s performance in block with unlimited decision time. *9.4* Participants judge allocentric spatial relations six percent more accurately after three map exploration sessions than after one (χ2(1)=26.9, p<0.001). Without time pressure participants make five percent more accurate decisions than when having to decide in three seconds (χ2(1)=31, p<0.001). Performance in the Pointing task is two percent below overall task average (χ2(1)=7.1, p=0.008). (Red dots) Single participant’s performance in the three-second condition for each spatial task, respectively. (Blue dots) Single participant’s performance for unlimited decision time. (Red line) Linear mixed-effect penalized least squares fit with participant as grouping variable across all three-second blocks. (Blue line) Linear fit across all unlimited decision time blocks. (Red area) The 95% confidence interval for the three-second condition linear fit. That is, 95 of 100 intervals each from a different sample include the actual population fit. (Blue area) The 95% confidence interval for the unlimited condition. *9.5* Compared to the Pointing task map participants under time pressure are five percent better in the Relative task (χ2(1)=16.3, p<0.001) and four percent better in the Absolute task (χ2(1)=10.9, p<0.001). The change in performance difference from the first to the third exploration session is eight percent for Relative to Pointing (χ2(1)=14.9, p<0.001) and six percent for Absolute to Pointing task performance (χ2(1)=9.6, p=0.002). (Black, Green, Yellow dots) Single participant’s performance in the Absolute, Relative and Pointing spatial task trial block, respectively, only for trials with time limit. (Black, Green, Yellow line) Linear fit across sessions in the condition with time limit, for Absolute, Relative and Pointing task, respectively. (Black, Green, Yellow area) The 95% confidence interval for the three-second condition linear fit for Absolute, Relative and Pointing, respectively. *9.6* Given unlimited reasoning time there are no significant in-between task effects. (Black, Green, Yellow dots) Single participant’s performance in the Absolute, Relative and Pointing spatial task trial block, respectively, only for trials without time pressure. (Black, Green, Yellow line) Linear fit across sessions in the free-reasoning condition, for Absolute, Relative and Pointing task, respectively. (Black, Green, Yellow area) The 95% confidence interval for the unlimited time condition linear fit for Absolute, Relative and Pointing, respectively.

For unlimited reasoning time we do not find any significant in-between task effects. Model (u-Map) results hint at Relative to be two percent above Pointing, however, this effect is not significant (Rel. to Poi.= -1.9, SE=1, CI=-3.8, 0; χ2(1)=3.7, p=0.054). Without this term in the model, a significant change of about five percent (Rel. to Abs.=4.6, SE=2, CI=0.3, 8.8; χ2(1)=4.38, p=0.036) of the difference of Absolute and Relative performance over session reduces to four percent and is, therefore, no longer significant (Rel. to Abs.=3.7, SE=2.2, CI=-.5, 7.9; χ2(1)=2.94, p=0.086; not in Appendix). Figure 9.6 depicts the results qualitatively.

To investigate participants’ overall development in the three tasks across sessions, comparing also both time conditions, we fit a model (a-Map) that incorporated the complete map data and time as an additional potential factor. The model does not provide evidence for H3, rather performance in the Pointing task is significantly below task average (mean to Poi.= -2.2, SE=0.7, CI=-3.6, -0.7; χ2(1)=7.1, p=0.008). This result speaks against H4 which purports Pointing to be the best task across all three embodiment conditions. Further likelihood comparisons do provide evidence for H5, that is performance increases without time pressure (time= 5.4, SE=0.8, CI=3.7, 7; χ2(1)=31, p<0.001).

Also, H6 holds since performance increases by about six percent over the three sessions (session=6.6, SE=1.2, CI=4.3, 8.8; χ2(1)=26.9, p<0.001). The results are illustrated in Figure 9.4.

Comparable to the VR embodiment results, visually assessing systematic differences in performance development from Figures 9.1 to 9.3 is barely possible. We see participants performing worse with limited decision time and performance increasing with session. When free to reason, participants show a strong increase in Absolute task performance from the first to the second session, similar to the VR condition. Also discernible is that performance in the Pointing task increases with session but remains below both Absolute and Relative performance in both time conditions. All in all, performance in the Relative task increases the most.

All model design matrixes and detailed results can be found in the Appendices C and D.

### VR Embodiment as Reference Condition

#### VR Versus VR with Belt

As outlined in the introduction, the embodiment condition while exploring the city in VR only, is closest to navigating an unknown large-scale environment without additional help from external devices. It can, thus, be considered a baseline or reference for acquisition of allocentric representations. The sensory augmentation belt provides the participant with egocentric information about cardinal direction, thereby making allocentric representations somewhat obsolete. In the following we directly compare VR- and VR with belt results through linear mixed models, adding exploration condition as new possible fixed effect. Note that this comparison is mostly of an exploratory nature. Only H4, H5 and H6 are supposed to hold across all three embodiment conditions during exploration. Since H5 and H6 have already been shown to fulfill this expectation we are not reporting the combined data results for these hypotheses here.

Neither model likelihood comparisons of task performance under time pressure (s-VRvB), nor modelling the combined free-reasoning results (u-VRvB) reveal any differences between allocentric representations acquired in VR with and without belt. On the contrary, combining both embodiment conditions, free-reasoning results reveal that the difference between Absolute and Relative reverses over the three sessions (Abs. to Rel. with session= -4.7, SE=1.4, CI= -7.5, - 1.8; χ2(1)=10.3, p=0.001). The interaction is similar to the one found in the VR condition and thus evidence for the robustness of the distinction between two forms of allocentric representations as purported in H1.

The polar histogram in Figure 10 depicts the orientation of participants when asked to turn towards north at the end of the exploration period in the VR city. It is apparent that participants with belt can distinguish north significantly better than participants without belt. Consequently, the fact that belt participants, after exploration, are qualitatively worse (compare Figure 8.6 and Figure 7.6) when judging the direction of north compared to VR participants without belt speaks for H2. We do not build stable representations usable in the absence of that which they represent when we do not need to. Representations are ecological.

**Figure 10.**
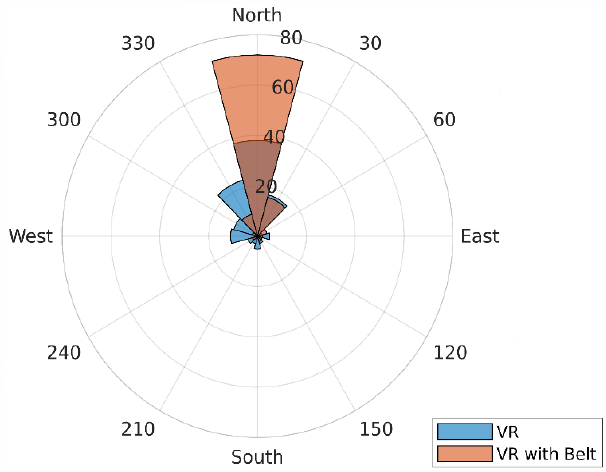
Polar coordinate histogram depicting the direction participants faced after having been asked to turn towards North. The command was given at the end of each exploration session. Each bin covers an angle of 30°. The numbers increasing radially depict the number of participants in that bin. All participants that only explored VR without belt are in blue. All participants that only explored VR wearing the sensory augmentation belt are in red. Participants with belt are about two times more often correct than those without belt.

Finally, incorporating both time conditions in one large model (aVRvBelt) provides evidence that Pointing performance is above overall task average (mean to Poi.=0.8, SE=0.3, CI=0.1, 1.4; χ2(1)=5.6, p=0.018). The stability of high Pointing performance in both VR conditions relative to the other two spatial tasks, as claimed in H4, emphasizes the impeding effect of map learning on Pointing which we will further discuss in the following section.

All model design matrices and detailed results can be found in Appendix D.

#### VR Versus Map

In the VR embodiment condition the participant has to move her legs in order to turn her body in the virtual 3D world. She also moves her head freely to look around. Both actions are necessary to explore the VR city and learn about its spatial layout. Through using a map as navigation device, a participant can gather all information necessary to solve the spatial tasks, while performing only a minimal amount of bodily movement.

In the following we directly compare VR and map results using linear mixed models through simply adding exploration condition as new possible fixed effect. Only H4, H5 and H6 are supposed to hold across all three embodiment conditions. Again, we will not report combined data results for H5 and H6 here.

Modelling (s-VRvMap) the task performance of the combined exploration conditions when the participant has to decide in three seconds creates no new insights. We refer the interested reader to the Appendix D.

Similar to the result of the comparison of VR with and without belt above, in the decision condition without time pressure the comparison between VR and map (uVRvMap) also reveals that the difference between Absolute and average task performance reverses over the three sessions (Abs. to Rel. with session= -3.2, SE=1.5, CI=-6.1, -0.2; χ2(1)=4.49, p=0.034). The finding is again evidence for the strength of the interaction effect which, if extrapolated, speaks for H1.

The largest model (a-VRvMap) likelihood comparisons marginalizing both time conditions reveal that the average task performance after map exploration is better than the performance after exploring the city in VR (exploration=2.3, SE=0.8, CI=0.7, 3.9; χ2(1)=8.2, p=0.004). Map is better than VR despite the fact that Pointing performance is below overall task average after map exploration but above average after VR exploration (mean to Poi. with explor.= -3.1, SE=0.7, CI=-4.4, -1.8; χ2(1)=20.4, p<0.001). The interaction is evidence for an effect of exploration condition on the validity of H4.

In order to find out if Pointing performance itself is worse after map than after VR exploration, and not just in relation to the other two tasks, we fit a model (p-VRvMap) with only the Pointing task data of both groups. We do not find a main effect of experimental exploration condition on Pointing performance (exploration= -0.9, SE=0.9, CI=2.6, 0.9; χ2(1)=0.87, p=0.35). We do also not find evidence that such an effect develops with longer exploration time, i.e. number of repeated sessions (explor. with session= -1.8, SE=1.9, CI=-5.6, 2; χ2(1)=0.88, p=0.35). For illustration compare Figure 7.5 and 7.6 to Figure 9.5 and 9.6.

All model design matrixes and detailed results can be found in Appendix D.

### Behavior of Participants

After three exploration sessions the general performance level in our spatial tasks reaches only about ten percent above chance level. The low performance potentially undermines the generalizability of our results. Floor effects, that means artefacts due to overall very limited acquired knowledge, might appear as confounds. That is not to say that our results are not reproducible, the statistical analysis is sound, but floor effects could make their interpretation significantly more involved.

Past studies have repeatedly found evidence for differences in spatial aptitude in humans. Some people seem to build more comprehensive representations or can use these more effectively than others (Ishikawa & Montello, 2006; Weisberg et al., 2014; Weisberg & Newcombe, 2016). If we could provide evidence for a similar distinction among our participants and show that the main results discussed above are stable across those differences in spatial learning ability, that would speak against floor effects.

As exemplified in Figure 11 participant’s performances can be approximated as normal in all task and time conditions across exploration conditions and sessions. That is, task performance does not reveal participant clusters of low and high navigation aptitude. We need to consider, however, that each experimental session provides us with six dependent variables which change their values with each session. Subgroups might split into different combinations of these variables, i.e., being good at Pointing in the first session could imply worse performance at the Relative and Absolute task in later sessions. Also, our participants might not split into subgroups all together, but rather fall into a more continuous spectrum of spatial ability.

**Figure 11.**
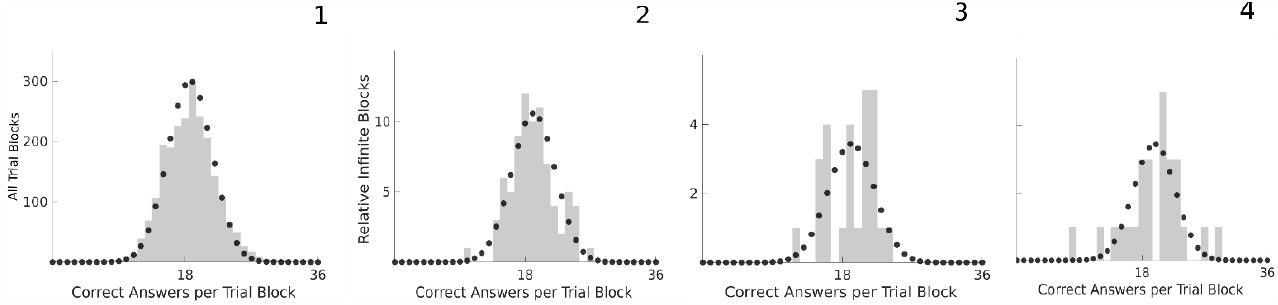
Performance Distribution. *11.1-4* Distribution of participants’ correct answers overlap with binomial distribution. (Gray bars) Histogram of correct choices summed over all trial blocks in 11.1 and a single exemplary condition trial block per session in 11.2-4. The single condition is the Relative spatial task with no time limit per trial and only for participants who explored VR. Shown are the first, the second and the third sessions, respectively. (Black dots) Binomial distribution B(p,n) with p equal to the ratio of correct trials to all n=36 trials per block.

A great advantage of our study design is the level of control. It allows analysis of a multitude of independent behavioral variables. These variables in turn might allow us to better identify individual spatial cognitive strategy and aptitude. We investigate the stability of our main results presented above in light of four behavioral factors. Percentage of houses seen, median looking distance, movement speed and self-report. For the following analysis we use the behavioral values of each participant’s first session only.

#### Percentage of Houses Seen

In Figure 12.1 to 12.3 we illustrate the percentage of houses seen or clicked for each participant in each embodiment condition. Each bin gives the number of participants that saw or clicked on that percentage of houses during one exploration session. For the VR and VR with belt conditions, we made use of the eye tracking data. We combined both VR goggle and eye tracking coordinates to compute a gaze ray and infer which VR house this ray has hit at any one moment. If the ray hit a house for at least 260ms we counted that house as seen. Details of the procedure can be found in the publication by Clay et al. (2019).

**Figure 12.**
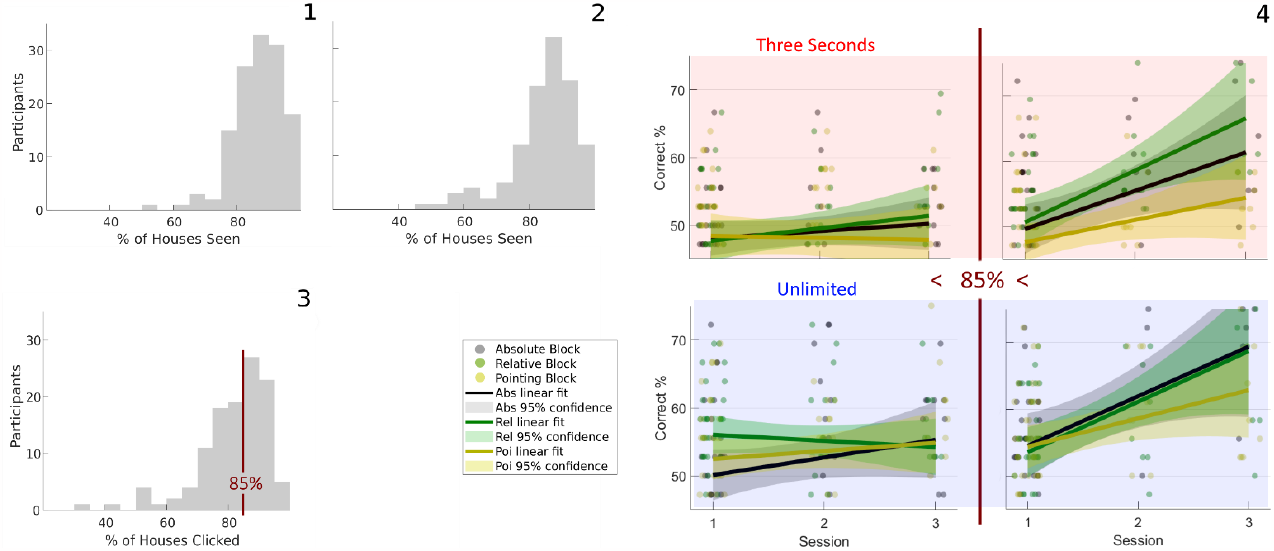
Percentage of Houses Seen. *12.1* Histogram of the percentage of houses seen during the VR experiment. Each bin covers 5%. Its height denotes the number of participants that have seen that percentage of houses during one exploration session. A house is counted as “seen”, when it has been looked at for 260ms. The total number of houses in the city is 213. *12.2* Histogram of the percentage of houses seen during the VR experiment with sensory augmentation belt. *12.3* Histogram of the percentage of houses seen during the map experiment. Each bin covers 5%. Its height denotes the number of participants that have clicked on that percentage of houses during one exploration session. After clicking on a house, the frontview of that house appears on the screen. The total number of clickable houses is 192. *12.4* The upper row depicts performances with three second decision time, the lower row with unlimited time. The left column shows the development of spatial task performance for map participants which clicked on less than 85% of houses during the first session. The right column illustrates the same for participants with more than 85% of seen houses. Under time pressure the mean percentage of correct spatial judgements rises by 16% (χ2(1)=11, p=0.001) with more houses clicked. With free reasoning time house-clicking behavior makes a difference of up to 35% (χ2(1)=15.1, p<0.001). (Black, Green, Yellow dots) Single participant’s performance in the Absolute, Relative and Pointing spatial task trial block, respectively. (Black, Green, Yellow line) Linear fit across sessions for Absolute, Relative and Pointing task, respectively. (Black, Green, Yellow area) The 95% confidence interval for the linear fit for Absolute, Relative and Pointing, respectively.

All three exploration conditions show a left-skewed unimodal distribution with similar amplitude and mode at around 85 percent. Note that each distribution is accumulated over all three sessions. For the following analysis we only use the values of the first session also for the second and third session of the same participant. We transform the distribution to a normal form using the exponential function before adding it as factor in the design matrix of its respective model. To evaluate the effect of the percentage of houses seen we simply add the factor to the full models of the limited- and unlimited decision time respectively and perform the step-down procedure described in the Methods section.

We do not find any effect in the VR and the VR with belt condition. In the map condition however, the number of houses clicked-on during exploration proves to be a highly significant indicator of performance both with (Model s-Map.h) and without time limit (u-Map.h).

Given only three seconds decision time an increase from about 50 to 100 percent of clicked houses leads to the average performance level rising with consecutive sessions by 16 percent (session with percent houses=16.3, SE=5, CI= 6.5, 26.2; χ2(1)=11, p=0.001). The behavioral factor also reveals the performance difference among Relative, Absolute and Pointing task over sessions to be itself a differential effect. The difference in performance increases with the percentage of houses clicked-on (Poi. to Rel. with session with houses=16.4, SE=4, CI= 8.4, 24.3; χ2(1)=13.7, p<0.001) (Poi. to Abs. with session with houses=11.8, SE=4, CI= 3.9, 19.7; χ2(1)=8.2, p=0.004).

Similar to the time pressure condition, with unlimited decision time performance rises faster with sessions if the participant saw more houses. The effect is the largest of any behavioral variable in our study, with a change in performance of 35 percent (session with houses=35.4, SE=8.9, CI= 17.9, 53; χ2(1)=15.1, p<0.001). The performance difference between Pointing and Relative (and Absolute, not reported) decreases when the number of clicked houses decreases (Rel. to Poi. with houses= -22.6, SE=11.2, CI= 44.6, -0.5; χ2(1)=4, p=0.04). Similarly, the difference between Relative and Absolute task performance shifts by about 25 percent (Rel. to Abs. with houses= -24.6, SE=11.2, CI= -46.6, -2.5; χ2(1)=4.8, p=0.03). Figure 12.4 illustrates the difference between participants that clicked on less- and participants that clicked on more than 85 percent of all houses during map exploration.

#### Looking Distance

The eye and head coordinates also allow us to infer the distance from which participants look at houses while they are exploring the city in the VR and the VR with belt condition. Median looking distance (weighted by the number of looking events per house, i.e. looking time) follows a log-normal distribution. We consequently use its logarithm as an additional factor in the limited and unlimited decision time models.

We find a strong effect of looking distance for belt wearers under time pressure (Model s-VRwB.d). Closely observing participants are 17 percent worse in the Relative task than in the Pointing task compared to participants who observed houses further away (Poi. to Rel. with distance (20m difference) =17.2, SE=5.5, CI= 6.4, 28; χ2(1)=9.7, p=0.002).

Similarly for the difference from Pointing to Absolute performance which increases by 14 percent for close participants (Poi. to Abs. with distance=14.3, SE=5.5, CI= 3.5, 25.1; χ2(1)=6.7, p=0.01). The complete model also features the interaction of Relative and Pointing performance with session (Poi. to Rel. with session =3.7, SE=1.9, CI= 0, 7; χ2(1)=3.93, p=0.047), as well as the interaction of Absolute to Pointing with session (Poi. to Abs. with session =5.4, SE=1.9, CI= 1.7, 9.1; χ2(1)=8.2, p=0.0042). That means the two interactions reported in the main results above are stable with respect to the behavioral factor of looking distance. The results are illustrated in Figure 13.1.

**Figure 13.**
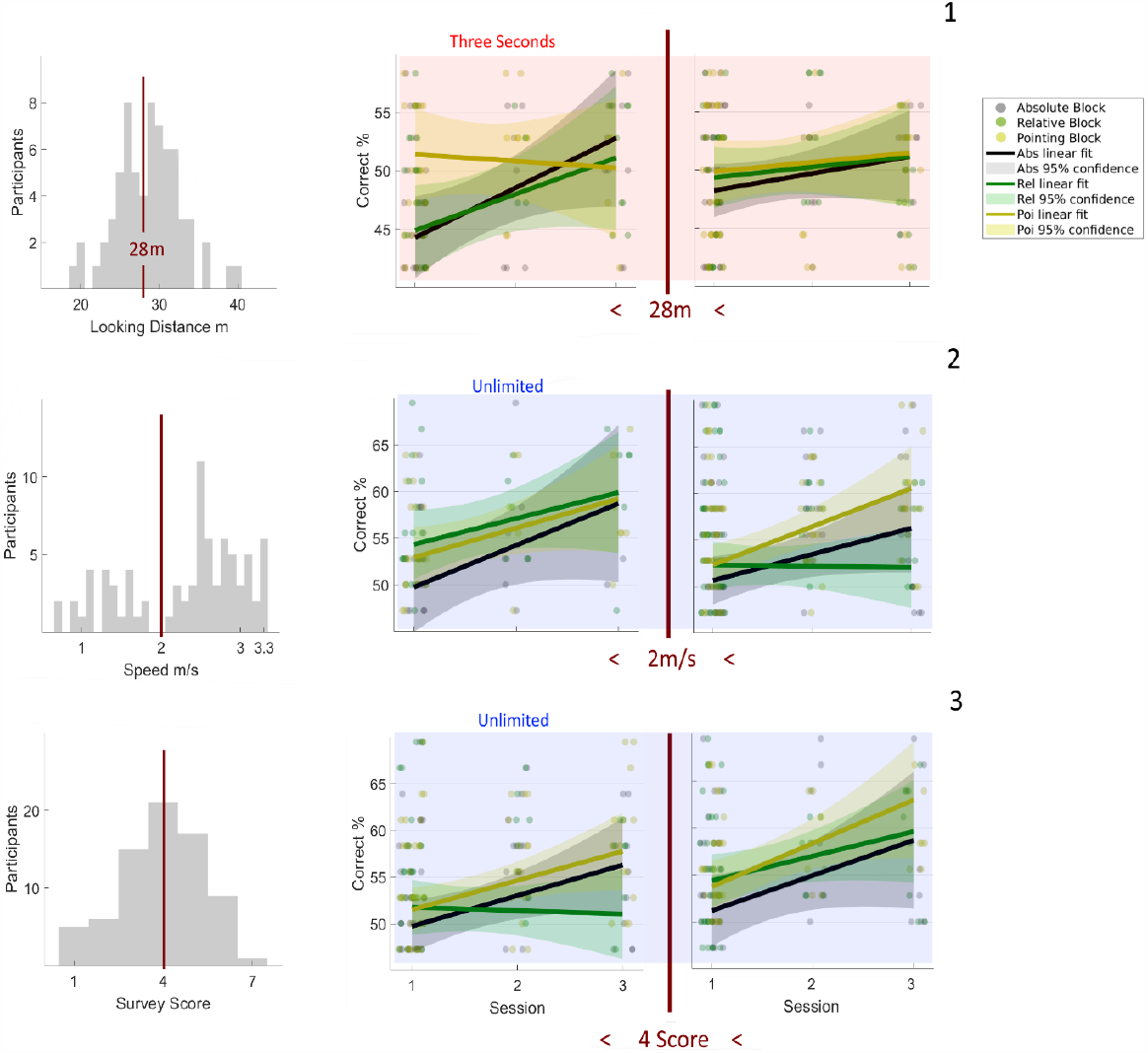
Looking Distance, Movement Speed and Self-Report. *13.1* Left panel, distribution of median distance from which participants looked at houses during their first VR exploration session with belt. Each histogram bar bins all participants with looking distance in an interval of one meter. Middle panel, development of spatial task performance under time pressure for VR with belt participants who stood closer than 28 meters when looking at houses. Right panel, same development for median looking distance greater 28 meters. Participants who explored the city while looking at houses close by are 17 percent worse at the Relative task in comparison to the Pointing task than those participants who looked at houses further away (χ2(1)=9.7, p=0.002). A similar effect appears for the Absolute task which is 14 percent worse than Pointing for participants observing near houses (χ2(1)=6.7, p=0.01). (Black, Green, Yellow dots) Single participant’s performance in the Absolute, Relative and Pointing spatial task trial block, respectively. (Black, Green, Yellow line) Linear fit across sessions for Absolute, Relative and Pointing task, respectively. (Black, Green, Yellow area) The 95% confidence interval for the linear fit for Absolute, Relative and Pointing, respectively. *13.2* Left panel, distribution of median movement speed per participant in the first session VR exploration (without belt). Each histogram bar bins all participants with median speed in an interval of 0.1 m/s. Middle panel, development of spatial task performance in the VR condition with unlimited decision time and median movement speed slower 2 m/s. Right panel, same development for participants with movement speed faster 2 m/s. The difference between Absolute and Relative task performance changes from slow to fast participants by three percent (χ2(1)=3.86, p=0.049). *13.3* Left panel, distribution of mean FRS questionnaire Likert score for the survey category for all VR participants (without belt) after their first experimental session. Each histogram bar bins all participants with a mean survey score in an interval of one centered on whole number scores. Middle panel, development of spatial task performance in the VR condition with unlimited decision time and median survey score less than four. Right panel, same development for participants with survey score greater 4. Marginal task performance increases eleven percent with increasing survey score (χ2(1)=14.7, p<0.001).

The differences in performance are largely due to Absolute and Relative being five percent below chance in the first session. We test if the closely observing participants performed so much below chance level because they did not react within the three seconds interval after which the trial is counted as wrong. Independent sample t-tests comparing the number of too late decisions do not show evidence for populations with different reaction times (Abs. close mean=2.8, SE=0.6, Abs. far mean=2.5, SE=0.3; t(68)=0.56, p=0.6) (Rel. close mean=2, SE=0.75, Rel. far mean=1.44, SE=0.27; t(68)=0.85, p=0.4).

Interestingly, we find no significant distance effects with unlimited decision time or for participants without belt.

#### Movement Speed

We further analyzed movement speed across participants in the VR and the VR with belt exploration condition. Recall that participants were able to regulate speed by moving their thumb up or down on a disc on a handheld controller. In contrast to other behavioral factors we find the distribution of median movement speed in both embodiment conditions to be bimodal (about one third of participants are slow). The bimodal form is consistent across sessions.

For the following analysis we only use the values of the first session. We perform a median split and map the participants’ velocities below and above the median to a binary code. We then use this code as an additional factor in the limited and unlimited decision time models.

Modelling the unlimited decision time condition (Model u-VR.s) reveals that the difference in performance between the Absolute and Relative task depends on the speed with which participants move (visually) through the VR city (Abs. to Rel. with speed (1.5 m/s difference) = -3.3, SE=1.7, CI= -6.6, 0; χ2(1)=3.86, p=0.049).

Importantly, the factor of speed does not interact with the effect of session on the differential learning of Relative and Absolute. Instead, adding the factor to the model increases the probability for H1, that is Absolute outperforming Relative with more experience in the VR city (Abs. to Rel. with session= -6.2, SE=2.1, CI= -10.4, -2.1; χ2(1)=8.5, p=0.004). For a visual comparison of both groups see Figure 13.2.

The median movement speed of the belt wearing participants does not inform statistically significant changes in spatial task performance.

#### Self-Report

As final behavioral factor we analyze self-report data on individual spatial aptitude and strategy. The data stems from the FRS questionnaire participants filled out in their first experimental session. The questionnaire assesses proclivity and ability of participants to use three forms of spatial knowledge. General spatial knowledge with a focus on egocentric strategies, survey knowledge and cardinal knowledge.

The distribution of general-egocentric answers shows a left-skew and was transformed with an exponential function. Self-assessed survey knowledge is normally distributed. Cardinal knowledge is perceived as being poor, that means it shows a strong right-skew and was consequently log-transformed.

Performance in the VR embodiment condition increases with self-assessed survey (Model s-VR.rs, u-VR.rs) and cardinal knowledge (s-VR.rc, u-VR.rc) in both time conditions. Survey knowledge shows the largest effect on average task performance without time pressure. Participants with the lowest self-assessed Likert score, one, perform eleven percent below participants with the highest Likert score, seven (survey=11.1, SE=2.7, CI= 5.7, 16.4; χ2(1)=14.7, p<0.001). See Figure 13.3 for an illustration of the effect. Compared to the Relative task, participants with high cardinal self-esteem are better at the Absolute task than those with low cardinal scores (Abs. to Rel. with cardinal=8.1, SE=2.3, CI= 3.5, 12.7; χ2(1)=11.8, p<0.001).

Neither factor interacts with the effect of session on the differential learning of Relative and Absolute. Instead the probability for H1 rises when including survey knowledge as additional factor in the unlimited decision time VR model (Abs. to Rel. with session= -6.4, SE=2.2, CI= -10.6, -2.1; χ2(1)=8.7, p=0.003). The same holds for cardinal knowledge as additional factor (Abs. to Rel. with session= -6.9, SE=2.2, CI= -11.2, -2.6; χ2(1)=9.9, p=0.002).

In the map condition, in addition to survey (u-Map.rs) and cardinal knowledge (s-Map.rc, u-Map.rc), general navigation ability mixed with egocentric strategies (u-Map.rg) also results in higher performance (general=8.4, SE=3.2, CI= 2.1, 14.6; χ2(1)=6.5, p=0.01). This could be an indicator for an influence of egocentric representations build up during map exploration.

In contrast to the above results, exploring in VR with belt only leads to an effect of self-assessed cardinal knowledge with limited decision time (s-VRwB.rc). Given that participants have to answer in three seconds, however, the effect is remarkably strong with an increase of ten percent in average performance over the three sessions (session with cardinal=10.1, SE=3.4, CI= 3.5, 16.7; χ2(1)=8.9, p=0.003).

That means, for participants with high affinity for cardinal directions the belt boosts building spatial representations which can be accessed quickly.

#### Limitations and Conclusion

All of the above results follow from an exploratory post-hoc analysis. For this reason, their significance needs to be interpreted with caution. Consequently, we refrain from adding more than one behavioral factor to a model at the same time although this would allow us to, for example, investigate interactions among behaviors. Even if some of our findings will turn out to be spurious in future experiments, it is remarkable that many behavioral effects increase the significance of the marginalized results presented before. While behavioral differences clearly indicate a spectrum of spatial aptitude and strategy among our participants, all significant results for, or against, our initial hypotheses remain stable. The analysis thus provides evidence against the existence of floor effects as confounds for our interpretation. All significant behavioral effects are presented with details on their respective models and results in the Appendix D.

### Item Behavior

Participant behavior indicates consistent differences in spatial navigation aptitude and strategy among individuals. The same might hold for item behavior. Here, item behavior simply refers to the different properties of items. Some of these properties can be ordered along a dimension, for example angular difference or distance between items, and thus allow for statistical analysis. In contrast to the participants’ behavioral variables discussed above, however, items vary during the course of a spatial task and among different tasks. Therefore, they also allow to identify differences among structures of spatial representations which might hold across all participants.

#### Angular Alignment

Concerning angular difference, reported elsewhere we do find performance in the Absolute task after map exploration to excel if the orientation of stimulus house and north are aligned (König et al., 2019). Alignment, here, means that the stimulus house front faces south. Thus, the participants’ egocentric perspective with respect to the item aligns with the cardinal direction of north. The map embodiment condition did not reveal alignment effects in Relative or Pointing task trials when the respective prime house faced south.

We tested the VR and VR with belt results for both north and east alignment effects. East was the direction all participants faced when starting VR city exploration. It has been shown that the direction from which one enters a new environment can become a reference direction which facilitates spatial tasks, e.g. pointing, even if that environment is large (Shelton & McNamara, 2004; He, McNamara, Bodenheimer, & Klippel, 2018).

For the analysis of each exploration condition we used repeated measures ANOVA with session, task, time condition and angular difference (0°, 30°, …, 180°) as independent variables. Neither of the two VR embodiment conditions, however, shows strong alignment effects. This lack of evidence for a dominant reference direction could be due to an interference of the task related importance of north and the starting direction of east.

#### Distance

The Relative and the Pointing task allow an investigation of the effect of distance between prime and target house on trial performance. We use simple linear regression to capture the relation of the percentage of correct participants in a trial and the distance of the trial house pair.

With unlimited decision time we find no correlation of trial performance and house pair distance in any session. When decisions have to be made in three seconds however, participants reveal a differential learning effect. Judging alignment of house fronts becomes significantly easier when both houses are close together compared to them being far apart.

In the VR condition this effect becomes strongest after three exploration sessions. In the third session participants are about ten percent worse when the correct house pair distance increases by 100m (distance= -9.16, SE=3.23, CI= -15.77, -2.55; t(34)=2.82, p=0.008, R2=0.19). VR participants with belt show a similar effect after two sessions (distance= -10.17, SE=3.5, CI= -17.32, -3.02; t(34)=-2.89, p=0.007, R2=0.2). Even map participants show a qualitative decrease of six percent for every 10cm (on screen) increase of pair distance (distance= -5.93, SE=3.08, CI= -12.24, 0.37; t(34)=-1.91, p=0.064, R2=0.1).

Remarkably, performance in the Pointing task shows no dependence on distance between house pairs after exploration in either embodiment condition. All regression model results for both time conditions are presented in Table 2. The findings are illustrated in Figure 14.

**Table 2.** House Distance Regression Results. Results of all simple linear regression models for the third session VR, third session map and second session VR with belt condition. Each model correlates trial distance of house pairs (prime and (correct) target) in the Relative or Pointing task and trial percentage of correct participants. The coefficient Distance is equal to the change in percentage of correct participants per 100 m increase in distance (except for map, there it is 10cm). Intercept denotes the putative correct percentage for zero distance between house pairs. SE is the standard error of the estimate of the coefficient and CI the confidence interval. The degree of freedom is always 34 because there are 36 trials per task block.

| Three Seconds |  | Distance<br>(Intercept) |  | SE (SE<br>Intercept) |  | CI |  | t(34) | p | R<sup>2</sup> |
| --- | --- | --- | --- | --- | --- | --- | --- | --- | --- | --- |
| VR | Relative | -9.16 | (61.46) | 3.23 | (1.65) | -15.77 | -2.55 | -2.82 | 0.008 | 0.19 |
|  | Pointing | -1.67 | (54.84) | 2.65 | (1.49) | -7.06 | -3.73 | -0.63 | 0.53 | 0.01 |
| Map | Relative | -5.93 | (65.62) | 3.08 | (1.58) | -12.24 | 0.37 | -1.91 | 0.064 | 0.1 |
|  | Pointing | 2.42 | (44.64) | 3.03 | (1.66) | -3.58 | 8.42 | 0.82 | 0.42 | 0.02 |
| Belt | Relative | -10.17 | (61.92) | 3.5 | (1.8) | -17.32 | -3.02 | -2.89 | 0.007 | 0.2 |
|  | Pointing | 2.95 | (44.1) | 2.01 | (1.13) | -1.16 | 7.05 | 1.46 | 0.15 | 0.06 |
| Unlimited |  |  |  |  |  |  |  |  |  |  |
| VR | Relative | 1.33 | (52.66) | 2.07 | (1.17) | -2.87 | 5.53 | 0.64 | 0.52 | 0.01 |
|  | Pointing | 3.08 | (55.04) | 2.54 | (1.21) | -2.07 | 8.22 | 1.21 | 0.23 | 0.04 |
| Map | Relative | -3.83 | (67.21) | 2.66 | (1.48) | -9.22 | 1.56 | -1.45 | 0.16 | 0.06 |
|  | Pointing | -0.5 | (58.85) | 2.45 | (1.15) | -5.45 | 4.45 | -0.2 | 0.84 | 0 |
| Belt | Relative | -2.42 | (59.92) | 2.1 | (1.17) | -6.71 | 1.87 | -1.15 | 0.26 | 0.04 |
|  | Pointing | 3.47 | (50.74) | 2.25 | (1.07) | -1.1 | 8.03 | 1.55 | 0.13 | 0.07 |

**Figure 14.**
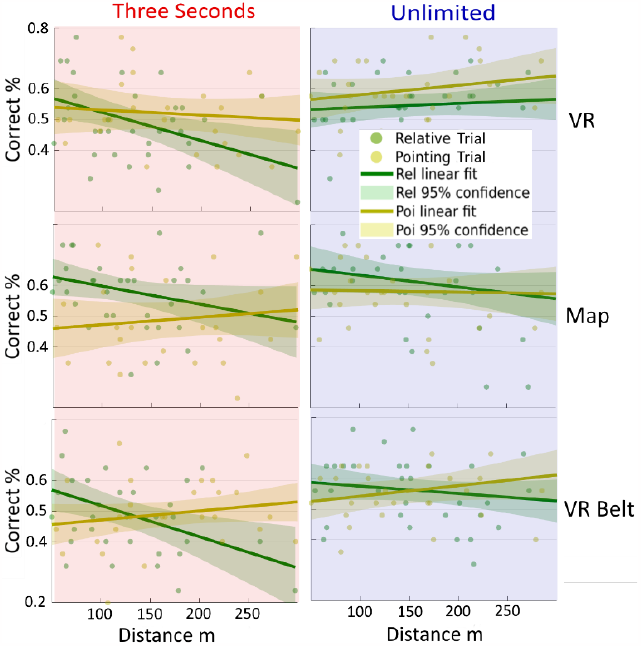
House Distance. (Green dots) Percentage of correct participants for the Relative task trial with that distance between prime and correct target house. Left column depicts performance under time pressure, right column without. First row shows VR experiment data of the third session. Second row shows third session, map experiment and third row second session of VR with belt. Distance is in meters for both VR experiments and in millimeters for map. For easier comparison only distances from 50 m to 300 m are shown. (Yellow dots) Similar for Pointing task trials with distance between prime and target house. (Green line and area) Linear fit over performance and 95% confidence interval for the Relative task trials. (Yellow line and area) Linear fit over performance and 95% confidence interval for the Pointing task trials. Only performance in the Relative task under time pressure decreases with increasing distance between houses in all embodiment conditions. Performance decreases by about ten percent for the VR condition without belt and with belt (t(34)=2.82, p=0.008, R2=0.19) (t(34)=-2.89, p=0.007, R2=0.2). Distance increase on the map leads to a decrease of six percent in spatial task performance (t(34)=-1.91, p=0.064, R2=0.1).

As already mentioned during the analysis of participant behavior above, these results follow from an exploratory post-hoc analysis and thus are to be interpreted with caution. Also, an extrapolation of the linear fit leads to performance values eventually reaching zero percent which seems not plausible. A non-linear (for example square root) decay model might describe the data more accurately. Because it is sufficient to show a distance effect and for simplicity of comparison, here we stick with the linear model.

The similarity of the distance-time-pressure findings in all three exploration conditions points towards a difference in the spatial representation accessed during the Relative task compared to the Pointing task.

## Discussion

### Spatial cognition is difficult. But not for all

As hypothesized, with accumulating exploration time average task accuracy rises above chance (H5). It nevertheless remains, in our view, surprisingly low. Comprehensive allocentric spatial representations of our VR city seem hard to acquire. In part, this might be due to a deprecated set of sensorimotor relations offered by our setup. Here the main factor is probably the lack of free walking (Chrastil & Warren, 2013; Ruddle et al., 2011) or at least the possibility to move forward through leaning forward with the upper body (Nguyen-Vo et al., 2019).

Another reason for the overall low task performance could be the size and complexity of the environment. The city spans about 450 times 500 meters and consists of over 200 individual houses arranged at all possible angles along a road network which is not grid-like.

Furthermore, the lack of task-specific instructions might have led to additional variance in explorative behavior and performance. Some participants might have performed worse than they would have with more specific instructions. However, this potential variance is, in part, welcome. Since at least the Relative task was probably unfamiliar to most participants prior to this experiment, intentional instructions might have led to unnatural exploration strategies. We decided for a compromise to address this possibility. Participants were instructed to explore freely. Through the introductory part, however, they were also aware of the spatial tasks to be completed afterwards.

As noted by Chrastil and Warren (2011), spatial intention can come in varying degrees. In a study by van Asselen et al. (2005), for example, participants underperformed who were led from one room to another with the explanation that the first room was chosen by mistake. Burte and Montello (2017) did not claim a mistake, but rather asked their “incidental” group to pay attention to the architectural appearance and design of the neighborhood through which they were led. Their study does not reveal significant differences in spatial task performance between incidental and intentional learners. Given that our participants knew that they would be tested about spatial relations, we believe our compromise between natural behavior and spatial intention to be a sensible choice. More task-specific instructions might have led to higher performance, but at the cost of generalizability.

#### Individual Differences

Tests of behavioral variability during exploration reveal mixed results. Considering the percentage of houses seen, both VR embodiment conditions (one with belt) show very similar and strongly left-skewed distributions, which means that most houses were seen by all participants. Statistical analysis did not reveal an effect of the number of houses seen on performance.

For the map condition the percentage of house fronts seen and thus the number of house symbols clicked-on makes a large difference. A possible factor might be that only a deliberate click, that is a motor action in addition to simply moving the mouse cursor over the house symbol on the map was necessary to see the house front.

The difference in clicking behavior implies a difference in spatial strategy and/or intention. The simplest explanation for the main effect of the number of houses clicked on spatial task performance might be that participants who clicked fewer times, were less motivated.

In the VR conditions the behavioral variable of movement speed might have been an indicator for spatial strategy and intention. Participants who moved more slowly through the city learned the Relative and Absolute tasks significantly better. Note, however, that the slow participants’ average performance still only peaked around 60% after three exploration sessions. The performance is low even for slowly exploring candidates.

#### Embodied Roots of Spatial Cognition

Building a comprehensive overview over large environments is generally a difficult task. That becomes even more apparent in spatial judgements under time pressure. In our three embodiment conditions, only after map exploration did participants’ judgements of spatial relations reach accuracy significantly above chance when asked to decide within three seconds. As predicted, participants generally needed to “take their time” to perform correct spatial judgements (H6).

The correlation can be interpreted as an argument against a close connection of bodily action and navigation skills. Consider, for example, competitions in robotics development, like the Amazon Picking Challenge (2017) or the ICRA Tidy Up My Room Challenge (2018). The embodied tasks tested in those challenges are of immense computational complexity, but easily completed by humans in a short span of time. Consequently, if knowledge about allocentric spatial relations is grounded in bodily action, then why does it take so long to make use of it?

In recent years, it has been shown that spatial tasks result in neural activity which resembles the allocentric and, importantly, the egocentric activity during original perception of the spatial scene (Chadwick, Jolly, Amos, Hassabis, & Spiers, 2015; Marchette, Vass, Ryan, & Epstein, 2014; Vass & Epstein, 2017). Already in 2007, Byrne, Becker and Burgess developed a comprehensive model close to these findings. The model has been refined and extended by Bicanski and Burgess (2018). Remarkably, a hallmark of both versions is the importance of “mock” motor efferents for spatial reasoning (see also Thorndyke & Hayes-Roth, 1980). Possibly, solving a task involving allocentric spatial representations in a large environment takes so much longer because it involves sensorimotor simulation or re-enactment of multiple egocentric representations. The importance of simulation in cognition has been argued before (Barsalou, 2009; Gallese & Goldman, 1998; Hesslow, 2002; Jeannerod, 2001). From a purely computational perspective it seems inefficient to involve sensorimotor relations shaped by the agent’s body when accessing allocentric relations. The findings indicate embodiment to play a major role in the development of spatial cognition in animals.

In the discussion so far, we have left out an important behavioral result, namely large individual differences in spatial aptitude. Self-reports on spatial strategies assessed by the FRS questionnaire correlate with individual differences in performance. These differences occur in both time conditions and can range up to ten percent. It is not clear how an embodied approach can encompass these strong effects of self-perceived spatial ability, which have been found in many other studies as well (Burte & Montello 2017; He et al., 2019; Ishikawa & Montello, 2006; Weisberg et al., 2014; Weisberg & Newcombe, 2016). There seem to be cognitive processes involved in spatial reasoning which develop somewhat orthogonal to the embodied root.

Indeed, the findings concerning egocentric simulation mentioned above focus mostly on how to transform the allocentric position of the goal location into an egocentric code relating directly to limb position and vice versa. It is believed that a sequence of such processes plays an integral part in creating a movement trajectory towards the goal. Not much is stated, however, about how the allocentric representation of the goal location is recreated. This process seems too fast and localized to involve sensorimotor simulation (Pfeiffer & Foster, 2013). Indeed, modeling approaches expect attractor dynamics, shaped by the hippocampus’ internal recurrent connectivity and activated by a given context, to suffice (Corneil & Gerstner, 2015; Gönner, Vitay, & Hamker, 2017). The models describe cognitive processes which rely on an internal connectivity structure for which the term embodiment seems misleading.

Due to lack of statistical power, in our study we did not check for similarities between spatial questionnaire responses and exploration behavior. Like the other behavioral variables which explain variance in task performance, some variance in spatial ability might directly relate to bodily movement patterns. Eye- and general-body-motion tracking devices can allow future studies to elucidate this relationship. Modern machine learning approaches could also be used to compare the explanatory power of bodily behavior and mobile EEG in predicting spatial ability, strategy and task performance.

### Egocentric Map Knowledge

In the VR and the VR with belt conditions, participants are better at pointing from one house to another than if they have to judge the orientation of a house relative to north or another house. These results are in line with our hypothesis that representations that allow us to find the shortest distance between two locations are important and that performance is favored by the familiarity of an action (H4).

Surprisingly, in the map condition, of all three tasks, pointing from house to house shows the worst performance in both time conditions. Note that this is not a main effect of experimental exploration condition on pointing performance. Although we find evidence that Pointing performance develops more slowly with map exploration than with VR, it is not significant. Rather, the interaction of exploration and tasks described above appears because map exploration boosts both Absolute and Relative performance while leaving the Pointing task unaffected. The selective boost of performance becomes especially apparent under time pressure.

Pointing is one of the most prominent measures of spatial expertise in research on human cognition, but the literature remains inconclusive as to the difference between spatial knowledge acquired through studying a map versus full-body exploration. One study showed no main effect of embodiment condition on pointing accuracy (Richardson et al., 1999). Others find full-body experience to favor pointing (Shelton & McNamara, 2004; Thorndyke & HayesRoth, 1980). Again, others show the opposite effect (Fields & Shelton, 2006; Moser, 1988).

In a recent study Zhang, Zherdeva and Ekstrom (2014) investigate pointing accuracy after exploring a (desktop) VR city or it’s map over multiple sessions. Importantly, they employ two different task setups when probing the ability to point to a house. Both setups have in common that participants have to rotate an arrow shown on a screen in front of them until it points to a certain building. In one setup, the screen only shows the arrow, the ground plane, and written instructions as to where participants are right now and where they should point towards. In the other setup, participants are first shown the VR city again, but without any task relevant houses. The participants walk through this version of the city until they give a signal to the experimenter that they feel “oriented.” Then the pointing trials begin, this time only showing instructions as into which direction to point, not where the participant is located. Zhang et al. find that VR exploration facilitates accuracy in the latter “oriented” setup. Map exploration on the other hand facilitates accuracy in the ground-plane setup. They also find these differences to be transient effects. After sufficient exploration time, that means with increasing level of accuracy, the differences vanish.

Our tasks are situated in between Zhang et al.’s oriented and ground-plane tasks. In addition, our interactive map allows participants to take on an egocentric perspective parallel to the VR ground plane when they are shown the picture of the housefront after clicking on the house symbol on the map. Consequently, the interaction Zhang et al. discovered might cancel a main effect of exploration condition on pointing accuracy in our study.

Their findings, however, cannot account for the boost of performance in the Absolute and Relative task after map exploration. To understand this difference between spatial knowledge acquired from a map versus spatial knowledge acquired in VR, we will first try to explain why map exploration should boost task performance at all. For this, we present two (not mutually exclusive) explanations. The first explanation emphasizes the difference in computational complexity between both embodiment conditions. The load of information an agent must process during full-body immersive navigation in order to be able to later solve the three spatial tasks is massively downscaled in the map condition. Indeed, that humans show comparable performance after VR and map exploration exemplifies the degree to which our cognitive structures are specialized to process sensorimotor information efficiently.

The second explanation for the general improvement in performance after map exploration relative to VR exploration emphasizes that map participants can perceive a large part of the spatial relations necessary to solve the tasks from one point of view, i.e. the map view. Allocentric and egocentric representations could, therefore, inform each other, a potential factor which might especially favor performance under time pressure. Such an interaction of both types of reference frames confounds analysis of allocentric spatial knowledge.

The well-documented alignment effect speaks for the existence of this egocentric confound. Learning spatial relations via map leads participants to form representations which allow them to better recall spatial relations aligned to the axes defined by the map (Evans & Pezdek, 1980; Richardson et al., 1999; Shepard & Hurwitz, 1984). Intuitively, this reads like a prime example of an allocentric frame of reference. Note, however, that the vertical north-south axis of the map aligns perfectly with the vertical bodily axis going from head to toe through the participant. Therefore, the effect can also be, at least in part, due to the alignment of real- and imagined egocentric heading direction while sitting in front of the map and facing a stimulus house that is oriented downwards on the north-up map (Shepard & Hurwitz, 1984; Richardson, et al., 1999).

Our hypothesis that map exploration should favor the Absolute task (H3) was partly informed by the alignment effect as well. Although we do not find such a general improvement, as reported elsewhere, we do find participants to excel in the Absolute task if the orientation of house and north are aligned (König et al., 2019). Note that alignment of a spatial reference system to, for example, initial heading has also been reported after full-body exploration of a large new environment (He et al., 2018; Shelton & McNamara, 2004). Thus, the map alignment effect and allocentric representation do not necessarily exclude one another.

However, while both the lower computational complexity as well as the additional egocentric reference frame can explain a boost in average performance, only the egocentric approach can account for the boost’s task selectivity, as we shall see below.

#### Visual Memory from One Perspective

So far, it remains unresolved that performance in the Absolute and Relative task excels after map exploration, while Pointing accuracy is not affected, at least not when compared to Pointing performance after VR exploration. To understand this effect, we first need to acknowledge that Pointing is fundamentally different from both the Absolute and the Relative task in our study. Only pointing necessitates knowledge of each house’s location. This difference is of special importance for the map condition.

Both, the orientation of a house relative to north and the orientation of a house relative to another house can be learned in an egocentric association process. During map exploration, participants can associate a VR house picture (later task stimulus) with its rectangular symbol on the map. They, then, only need to recall this rectangle’s orientation in order to solve the Absolute task. They do not need to compare the orientation of the memorized rectangular house symbol with the orientation of the map because, as mentioned before, their bodily orientation suffices. North is up. Similarly, for the Relative task participants can mentally compare two rectangles’ orientations by accessing a single-point-of-view representation of the map sequentially. The Absolute and Relative tasks are largely solvable by accessing one and the same egocentric frame of reference multiple times. Shelton and McNamara (2004, p. 168) corroborate this idea: “the visual overlap for same-perspective and same-orientation images may allow participants to appeal to only the visual representation and base the judgment on image matching.” In contrast to the VR and the VR with Belt experiments, no further spatial reasoning involving mental rotations is necessary.

Only when the participant has to point from one house to another does the recall of the house symbols’ orientations become insufficient. Participants then need to represent the orientation of the prime house symbol on the map *and* both the prime- and target-house symbols’ locations in order to infer the correct pointing direction. The inferring process might involve rotation of a mental vector to find the shortest distance from one location to the other. Consequently, map participants might have relied more strongly on allocentric knowledge for the Pointing task compared to the Absolute and Relative task.

Recall that a hallmark of egocentric knowledge is the ability to quickly transform it into bodily action. The differential boost in performance appears when participants have unlimited reasoning time before an answer is required and is also significant when participants have only three seconds to answer. The finding under time pressure is additional evidence for the purported distinction of an egocentric and an allocentric strategy to solve the three spatial tasks after map exploration.

The behavioral variable “percentage of houses seen” reveals two encoding subgroups. People that clicked on more (different) houses were up to 30 percent better at answering the spatial tasks than those that clicked on fewer, although both clicked on enough houses to perform comparably. Furthermore, the more-houses group shows the differential boost of performance for the Relative and the Absolute task in both time conditions. The less-houses group shows no differential performance boost. Also, accuracy in all tasks only reaches about 55 percent. The differentiation into high and low performers might be due to individual differences in spatial intention and/or spatial strategy. If we associate the number of clicks with spatial intention, participants who clicked on fewer houses might simply have paid less attention to the spatial relations on the map. A possible reason might be lack of motivation. Finally, we want to emphasize that a group distinction is an oversimplification. The variable of percentage of houses seen shows a continuous unimodal distribution. In reality, participant motivation appears as a spectrum.

In our study map learning shows evidence for a strong effect of egocentric representation on performance in some tasks asking for allocentric spatial relations. Parallel development of egocentric and allocentric representations is well documented (Burgess, 2007; Gramann, 2013). However, research systematically investigating such a differentiation during map learning is sparse (Zhang et al., 2014; Zhang, Copara, & Ekstrom, 2012). Future studies might be able to elucidate the relationship between egocentric and allocentric knowledge during map learning using a similar task design as presented here.

### Affordances Shape Allocentric Spatial Knowledge

After VR exploration, three seconds of decision time is barely enough for our average participant to reach chance performance when judging house-to-north or house-to-house alignment (note that we counted no answer as a wrong answer). In other words, after three consecutive exploration sessions, which add up to 90 minutes in our VR city, the acquired allocentric spatial representations do not yet reliably inform intuitive decisions for most participants. Some, in particular those participants with high self-assessed survey or cardinal knowledge, however, have already formed representational structures allowing quick computations.

Averaging all VR (without belt) participants in the unlimited decision time condition, we find a significant interaction between task performance and overall exploration time. With little spatial experience, Relative performance is better than Absolute, and this difference reverses with more experience. Including behavioral variables strengthens this interaction. For example, people that move more slowly through the VR city show more improvement, both in the Relative task and the Absolute task, such that the interaction remains stable.

As a first explanation for a difference in task performance one should always consider the differences in task setup. A participant needs to recall at least two houses to solve the Relative task, but only one to solve the Absolute task. Hence, the Relative task might be more difficult. On the other hand, it is not clear why the probability to be able to recall the direction towards north should be higher than the probability to have seen and memorized the orientation of the second house. In any case, the explanation seems unlikely because, as stated above, with little spatial experience participants perform better in the Relative task than in the Absolute task.

An explanation for low Absolute task performance after the first VR session could be that the participants simply did not find north. Only starting in the second session were they able to infer the cardinal direction of north during one virtual day using the movement of the sun and shadows alone. This would explain the exceptionally strong increase in performance from the first to the second session in the Absolute task.

However, this “overload” claim seems insufficient to explain why we find the same apparent jump from the first to the second session after map exploration. The rectangular map is north-up and at all times fully visible to the participant. We, therefore, do not believe the possibly confounding factor to be decisive for the differential task development in the VR condition. However, because performance development in most tasks and across all exploration conditions seemed to best fit a linear model, we did not focus our limited statistical power on the apparent non-linearity in the Absolute task. This effect remains to be elucidated by future research.

Also, if the interaction of house-to-north and house-to-house accuracy with exploration session were largely due to difficulties in finding north, the differential development should disappear for participants with high cardinal self-esteem. This is not the case. As mentioned above, including the behavioral variable of self-reported cardinal knowledge into the model heightens the probability for the interaction of Absolute and Relative performance over sessions.

Another argument against “overload” comes from the fact that extrapolating our results leads to similar findings as reported in the pilot study with residents of the city of Osnabrück (König et al., 2019). They found a reversal of task accuracy with decision time. Under time pressure, participants were better at judging house-tohouse orientations, but with more decision time they excelled when aligning houses and north. We expected to find the interaction with decision time of the pilot study in the VR condition as well (H1). We cannot reproduce the Osnabrück study’s results exactly. However, given that finding north during exploration was not too difficult for the participants and the development of the Absolute task can be approximated as linear, we can extrapolate our interaction of task and session. The Absolute task would then eventually outperform the Relative task, when deciding without time pressure, as expected.

One possible explanation for the need of extrapolation is that, while our study observed the developing structure of allocentric knowledge in a completely unfamiliar city, the pilot study probed the structure of spatial knowledge in a familiar environment. Pilot study participants had resided in the city of Osnabrück for at least a year, meaning that their spatial knowledge of the city was significantly further developed. As it stands, our study complements the pilot study’s findings since our focus lies on the development of allocentric representations.

Relating our results to the Osnabrück study has to be done with caution. That is because we cannot exclude that the high performance in the Osnabrück-north-pointing task without time pressure was due to participants having seen maps of the town beforehand. Exploration with the help of a map can lead to representations aligned with the map layout (Meilinger, Frankenstein, Watanabe, Bülthoff, & Hölscher, 2015).

For the possible confound speaks the fact that participants in the Osnabrück study performed best when house orientation and south were aligned, that means, in trials where participants were oriented towards north. Similarly, Frankenstein et al. (2012) found that participants performed best when prompted to point towards a goal location which lay in the direction of north in their hometown.

On the other hand, it is not clear why such map knowledge should not also lead to better performance when comparing house and north under time pressure. The possibly confounding factor cannot explain the interaction of Absolute and Relative task with decision time. Consequently, while map knowledge might play a role in the Osnabrück study’s findings, there remains a significant amount of variance which asks for a different explanation.

We argue that both the interaction of Absolute and Relative performance with exploration time, as well as their interaction with reaction time in the pilot study, are caused by the difference in the two tasks’ relation to embodied action. Both tasks are equivalent in that they ask for angular differences between directions of orientation only. Knowledge about relative location is irrelevant, and the action-relation of angle is rotation. The difference between the two tasks is that of house orientation and cardinal direction. What makes the difference significant is that these spatial concepts afford different actions.

#### Local Binary Code Versus Global Survey Code

Cardinal direction is important when traversing long distances in a straight line. It allows the navigator to avoid obstacles while keeping the global direction of movement constant. As long as a landmark, like a large building, remains visible during travel, it can be used in the same manner.

When the navigator is situated within an environment where landmarks are themselves part of visually separated vista spaces, their role changes (Lynch, 1960). These landmarks connect routes (Siegel & White, 1975; Montello, 1998; Chrastil, 2013). They are the start and goal locations of a journey, but they can also become obstacles on the way. Furthermore, recognizable buildings along the way can help the navigator turn at the right moment. Consequently, such local landmarks become decision points on the navigator’s journey, often associated with a change of direction, which implies whole-body rotation.

It might be ecological to code the relative orientation of these “nodes of the decision system” (Siegel & White, 1975, p. 29) in direct relation to all other already memorized landmarks (König et al., 2019). If the representational structure would be constituted by direct linkages between pairs of landmarks, that is pairs of houses, it could be learned and accessed quickly. We call this kind of representational structure a binary code. We will further refer to the landmarks the code relates to each other as nodes.

However, while the ecological value of survey knowledge (finding the shortest route) increases with the scale of the environment, information about relative orientation of landmark nodes becomes less important. In line with this reasoning, we find that when participants have to judge the relative orientation in three seconds, their performance decreases with increasing node-pair distance. The effect does not appear with unlimited decision time. The binary representational structure seems to be preferably accessed when quick judgement of rotation among local nodes is needed. Such a code might be helpful for intuitive action sequences occurring, for example, during hunting and hiding. The finding is in line with the Osnabrück study’s results which revealed Relative to outperform Absolute under time pressure (König et al., 2019).

Indeed, that connection strength should decrease as a function of distance seems a necessary property of the all-to-all linkage form of the representation. Otherwise, given any number of nodes N, it would allow access to the relative orientation of all N-squared node pairs (divided by two because the relations are symmetric and subtract N because each node is identical to itself). In other words, without a mechanism to weaken links between node pairs a linear increase in node number leads to a quadratic increase in the information carried by the binary representation.

The defining characteristic of the cognitive map, global deduction of short paths, implies knowledge of the relative location of the starting node and the goal node. That means knowledge of allocentric node location. We can refer to the underlying representation as survey code. Remarkably, pointing from node to node does not reveal a qualitative tendency for a decrease of accuracy with increasing distance. The invariance speaks for survey code to be structured differently than representation of allocentric node orientation, that means binary code.

Weisberg et al. (2014) have shown that pointing accuracy does depend on distance, even with reasoning time. In their experiment, if participants point from a building in one neighborhood to a building in another neighborhood, pointing accuracy is lower than when participants point in one neighborhood from house to house. They identify participants who do not show the distance dependence as good spatial integrators. We posit that, without time pressure, no distance dependence only holds for well-integrated environments. The explanation also fits the lack of clear distance effects in hometown pointing studies (Frankenstein et al., 2012; König et al., 2019).

If we point to a node which lies in a certain direction, e.g. north, the further away the node is located, the closer our pointing direction becomes to north. Knowledge of cardinal direction can, thus, be considered the limit case of being able to point to a node as far away as possible. That means it enables us to reach the node on the shortest (in principle) possible path. Consequently, knowledge of cardinal direction also depends on a survey-like representation.

Finally, as mentioned above, without time pressure we do not find an effect of distance on judgment of relative node orientation. Hence, the different encodings need not be strictly separate. The survey code might hold some information about the relative orientation of houses as well. Alternatively, given enough reasoning time survey- and binary code may be used together in order to solve a spatial task.

A possible test for the distinction of binary- and survey code on the neurophysiological level could be done as follows. There is ample evidence from experiments with rats that changes in local environmental cues lead to different changes in encoding than deforming the global geometric shape of the environment (for a review, see Latuske, Kornienko, Kohler, & Allen, 2018). If we identify place cells as part of a survey code, then our above analysis predicts only a small effect on these cells when manipulating the relative orientation of landmarks inside the arena. We do not expect a complete remapping. Rather, the change in landmark orientation might affect the firing-rate coding (Leutgeb, C., Leutgeb, J.K., Barnes, Moser, E.L., McNaughton, & Moser, M. B., 2005). If a change in the orientation of landmarks would have a similar effect on neuronal properties as the change in the relative location of landmarks our proposed distinction of binary- and survey code might be mistaken.

Were it not for the Absolute task, participants might not have learned the direction of north in our VR city. Nevertheless, there is evidence that survey knowledge of large environments is aggregated along (at least) one global reference direction (Shelton & McNamara, 2004; Meilinger, Riecke, & Bülthoff, 2014; He et al., 2018). Given sufficient exploration experience, alignment of node orientation and such a reference direction should become part of survey code.

In our VR condition (also with belt), we do not find alignment of house orientation and the cardinal direction of north to significantly improve task performance. The global reference direction, however, often coincides with the first perspective from which one enters a new environment (Shelton & McNamara, 2004; He et al., 2018). Since our participants always started from the same (central) position in the city facing east, we also tested for alignment effects in that direction, again without significant results. It is possible that knowledge about the direction of north and initial starting orientation east interfered destructively. In order to firmly establish the effect of a reference direction in future experiments, both task direction and starting orientation should coincide. Alternatively, one of the two should be randomized.

Even if one were to accept the distinction between binary- and survey code outlined above, one might ask if it is indeed the difference in bodily action relating to house orientation and cardinal direction of north respectively which cause the different forms of representation. The pilot study revealed an additional argument for the distinction in coding being due to the actions associated with the spatial concept coded. Recall that the Osnabrück researchers found the Relative task to outperform the Absolute task under time pressure. However, when judging the direction of streets relative to one another or aligned to north, participants were quicker in the north task (König et al., 2019). This finding speaks for the orientation of streets being primarily encoded in a structure that develops along a global reference direction, that is survey code. Indeed, while streets can also serve as landmarks, they afford different actions than the kind of landmarks we considered above as nodes. Streets do not afford rotation. On the contrary, streets afford long, straight-line movement. Therefore, the example is further evidence for the representational structure of a spatial concept depending on the embodied action it allows.

In conclusion, notwithstanding the possible confounding factors, our embodied interpretation can explain a considerable amount of the combined variance of both the Osnabrück and the VR exploration results. We will see in the following sections how it can also relate to the VR with belt results.

We are not the first to argue that spatial representations are to a large extent grounded in action possibilities. Greene and Olivia (2009) have shown that cognitive models classifying natural scenes according to global and action-related properties like depth or navigability are better than classifiers attending to local geometric scene properties. Computational models for human navigation consistently reveal the importance of pseudo-motor signals during spatial reasoning (Byrne, Becker, & Burgess, 2007; Bicanski & Burgess, 2018). Physiological evidence for the coding of navigational affordances in the human visual system has recently been put forward by Bonner and Epstein (2017). They conclude that there exists a bottom-up mechanism for perceiving potential paths for movement in one’s immediate surroundings.

### Augmentation Versus Representation

Without time pressure, as expected, participants in the VR with belt exploration condition perform significantly better in the Relative task than in the Absolute task (H2). The sensory augmentation device indicates north via vibrotactile stimulation. Therefore, naturally, when asked to turn towards north at the end of the VR and the VR with belt exploration session, respectively, participants were much more accurate with the belt. Why then, should we expect performance to increase when participants judge house-to-house alignment, rather than when they judge the relative orientation of house front and cardinal direction, after exploration?

Prior research investigating changes in spatial perception and navigation strategies in long-term belt wearers reported “automated navigation without mental reflection” (Kaspar et al., 2014). The additional sensorimotor loop provides constant information about cardinal direction in an egocentric reference frame. It, thereby, makes it unnecessary to represent north in a cognitive map. Indeed, building survey code itself becomes less ecological, at least in its former structure. The belt enables the wearer to globally move in a straight line in any direction with minimal cognitive effort.

Neurophysiological findings corroborate the claim that the belt impedes development of long-term survey knowledge. The hippocampus is a brain region important for allocentric spatial memory recall as evidenced, for example, by Maguire et al. (1998). In their study the hippocampus shows larger activity when participants have to mentally deduce a new route due to obstacles appearing in an otherwise familiar environment. That is compared to neuronal activity in the hippocampus when participants can simply follow a known path. Sabine König and colleagues found reduced activity in the hippocampus in participants who had trained with the feelSpace belt over the course of seven weeks (König, et al., 2016).

Note that, if cardinal direction is part of survey code, so is relative location. Therefore, lower performance in the Absolute task indicates lower performance in the Pointing task as well. We did not perform a statistical analysis because our original hypothesis focused on the effect of the belt comparing Relative and Absolute task. However, at least qualitatively, Pointing performance for belt wearers is worse than for those participants who explored the VR city without belt, as expected.

In addition to the decline in quality of survey code we expected the information provided by the belt to enhance the binary representation linking house-nodes. The binary representation, as posited above, is the primary form of representation for relative house orientations. We hypothesized that the additional vibrotactile experience of the direction of north while a belt participant encountered a new landmark would add to the strength of the binary linkages. However, while we did find the expected effect of Relative outperforming Absolute, Relative task performance was not better than after VR exploration without belt.

Unconscious integration of the information provided by the new sensorimotor loop into a spatial representation that can be recalled in the absence of that which it represents seems not effective after 90 minutes of wearing time. Participants might have had to consciously map the vibration around their waist to the concept of north in order to make use of it. Some direct form of understanding of the lawful change in neural activity caused by the interaction of VR north, the belt, and the participant’s body was missing. The augmentation belt had not developed into a truly a new sense, yet.

That sensorimotor relations have to be learned before a sense can be fully used as such is well known (White et al., 1970; O’Regan & Noë, 2001; von Senden, 1960). Indeed, only long-term belt wearers reported changes in spatial perception (Kaspar et al., 2014). Also, neural signatures indicating changes in sensorimotor processing in belt wearers have been recorded over several weeks (König et al., 2016).

The lack of belt training somewhat impedes our results. On the other hand, it allowed us to collect a sample large enough to investigate subgroups of spatial learning. Distance from which houses were looked at and self-reporting on knowledge of cardinal direction both correlate with the speed at which belt participants were able to acquire spatial knowledge. These findings point again to the importance of individual differences and how behavioral effects can be used to inform experimental interpretation, also of experiments involving sensory augmentation devices.

The VR with belt embodiment condition indicates that new sensorimotor relations can lead to profound changes in the development of allocentric spatial representations by changing their ecological value. Also, that judgments of the relative orientation of buildings are not affected by the belt, is further evidence for the difference between the encoding of house orientation and cardinal direction as posited above. That is evidence of the existence of two forms of allocentric representations, namely local binary and global survey code. Schumann and O’Regan (2017) have presented a sensory augmentation device indicating the direction of north via auditory signal. After manipulating the device to indicate north too early or too late during self-rotation, they observed long-term recalibration in vestibular rotation judgements. That means evidence for fast integration of the form of spatial information provided by the new sensorimotor loop. Comparing the effects of the unmanipulated device on the formation of allocentric knowledge would be a possible test for the ideas presented here.

### Development of Allocentric Spatial Knowledge

Knowledge about spatial relations among objects in a large environment is believed to build through memorizing landmarks, learning routes, and connecting those to attain survey knowledge (McNaughton et al., 2006; Siegel & White, 1975). However, the extent to which these constitute three separate processes, and whether these are the only processes involved, remains unclear (Chrastil, 2013; Ishikawa & Montello, 2006; Montello, 1998).

Montello for example, emphasizes the continuous and parallel acquisition of all three kinds of knowledge (Ishikawa & Montello, 2006; Montello, 1998). Chrastil (2013) adds the formation of topological graph knowledge as a process on the way from simple route to metric survey knowledge (see also Poucet, 1993). She further argues that, there does not exist a clear one-to-one mapping between neural activity and the purported milestones of spatial knowledge acquisition, at least not beyond the connection of place- and grid cells with the final form of survey knowledge. Rather, there exists a multitude of sub processes the interaction of which depends on higher cognitive and/or environmental factors.

The idea that graph knowledge is part of a developmental process towards metric survey knowledge is debated. Some authors believe the idea of a cognitive map, which generally stands for global survey knowledge, should be completely substituted by cognitive graphs (Meilinger, 2008; Warren, Rothman, Schnapp, & Ericson, 2017; Warren, 2019). A node in such a graph is made up of a salient landmark. To get from one node to the other, we need to recall the visual and/or motor transformations experienced while travelling between both. This sort of route knowledge accessible at a certain node is called a label and makes up the edge connecting two nodes. Survey knowledge then consists of accumulating new nodes and edges while, at the same time, strengthening the ability to access well-known and important ones (Warren, 2019).

Even in such a labeled cognitive graph, however, the nodes need to hold some kind of local reference frame relative to which the properties of the edges can be defined. Only if a landmark possesses an orientation, for example, can we speak of something being rotated relative to that landmark. An angle only exists in relation to a reference direction. The need for a reference frame brings about a dilemma. On the one hand, if the node itself makes up the local reference frame we find ourselves facing a cognitive map again, only a smaller version than before. On the other hand, if the local reference frame is itself a labelled graph, then each node in that smaller graph, again, needs its own local reference frame relative to which something can rotate and so on, leading to an infinite regress. Tobias Meilinger (2008) proposes a synthesis of cognitive graph and cognitive map to account for survey knowledge. He hypothesizes that a node does not simply represent a landmark. Rather, each node represents the landmark embedded in part of its surrounding area, i.e. the local vista space. One major claim of his theory is that grid cells only extend over such vista spaces, or nodes. In line with Meilinger’s model of survey knowledge as a network of local reference frames or mini-maps, environments with opaque borders have been shown to lead to compartmentalization of grids in humans and other animals (Derdikman et al., 2009; He & Brown, 2019).

The issue of cognitive map versus graph remains debated. We believe, however, that most authors would agree with Chrastil’s emphasis on the importance of sub-processes, or cognitive processes in between the acquisition of landmark, route, and survey representations. Our findings corroborate her emphasis and indicate a major role for the different actions associated with, or afforded by, the spatial concepts represented. We, thereby, complement past research on the effect of bodily cues on spatial integration (Chrastil & Warren, 2013; Nguyen-Vo et al., 2019; Ruddle et al., 2011; Waller et al., 2004).

Another question remaining to be answered is how does it come about that some people are consistently so much better at integrating spatial knowledge than others (Ishikawa & Montello, 2006; Weisberg et al., 2014; Burte & Montello, 2017; He et al., 2019)? The consistent individual differences in spatial aptitude point towards differences in neural structure which cannot be readily explained by sensorimotor effects alone.

Lastly, the question arises how much of the processes involved in developing survey knowledge of large-scale environments can be mapped to developing knowledge in other domains. Finding the shortest route towards a goal in an environment which cannot be “overlooked” in its entirety shows interesting parallels to general problem solving (Siegel & White, 1975; Tolman, 1948). Furthermore, recent experimental findings have found parallels between navigating environmental- and mental space relating to the activity of grid cells (Bellmund et al., 2018; Buzsáki & Moser, 2013; Constantinescu et al., 2016; Epstein et al., 2017; Schiller et al., 2015). Spatial cognition is a promising research direction to disentangle sensorimotor effects on representational structure from other, complementary factors, e.g., genetically predetermined, neurophysiological boundary conditions.

## Conclusion

Our study investigates the relationship of embodied enactivism and cognitive representation. We focus on the development of allocentric spatial knowledge. The representations underlying this knowledge are particularly interesting because they code for relations among objects which are not part of the human body.

Low accuracy when judging allocentric spatial relations even after repeated exploration of a VR city indicates that spatial cognition involving large environments is difficult. Further evidence comes from the fact that these judgments require a considerable amount of time. Other research indicates cognitive simulation to play an important part in spatial reasoning. We conclude that the difficulties posed by spatial cognition in large environments are due to its rootedness in bodily action. On the other hand, independent behavioral variables allow us to distinguish individual participants who excel at spatial reasoning. Their outstanding spatial skills imply orthogonal cognitive structures supplementing the embodied processes under investigation.

Another major finding is the evidence for two different forms of allocentric spatial representations. Although participants never see the VR city from a bird’s-eye view, a reference direction and the relative location of landmarks can still be learned to greater precision than relative orientation of landmarks, as the former two are more important for global movement on the shortest path to a goal. The resulting contrast in representational quality also resembles the difference between the bodily actions the spatial concepts afford. The spatial concept of landmark orientation, on the one hand, affords local change of direction that means, rotation. Allocentric reference direction and landmark location, on the other hand, allow for global straight-line travel. The underlying representation is the survey code also known as a cognitive map. The primary form of encoding of landmark orientation, however, is a binary structure of local landmark pairs. This distinction of two forms of allocentric representations is further corroborated by the result that the representational quality of relative landmark orientation decreases with distance between landmarks. Representational quality of relative landmark location, on the other hand, is not affected by distance.

Participants equipped with a sensory-augmentation device providing constant information about reference direction while they explore the VR city develop impaired survey knowledge. In the spatial tasks after exploration, relative orientation of landmarks is judged with greater precision than alignment of landmark and reference direction. Sensory augmentation can transform an allocentric spatial concept, like reference direction, into an egocentric experience. The transformation does not affect the ecological value the spatial concept affords for navigation, namely, moving on the globally shortest path toward the goal location. The egocentric shift does, however, make it less ecological to build the survey knowledge necessary for successful navigation without the belt. After all, the information about a global reference direction is constantly available. Consequently, cognitive resources are shifted to other processes, leaving the participant with impaired survey knowledge if she disconnects from the sensory-augmentation device. The shift in cognitive resources does not affect judgment of relative landmark orientation and, thus, substantiates the existence of two forms of allocentric spatial representations, namely global survey and local binary code.

The development of spatial-task performance after exploring the VR city by means of a 2D city map is distinct from the results after VR exploration. Map learning boosts accuracy both with and without time pressure, but only when participants have to judge reference direction or relative orientation of landmarks. We illustrate that the differential increase in accuracy is largely due to an egocentric frame of reference. This egocentric interpretation rests on the fact that the map allows for an overview of the spatial relations among landmarks in the VR city from a single point of view. No further translational motion involving the participant’s body is required. The findings suggest that embodiment during exploration affects the structure of the acquired knowledge.

In conclusion, our study illustrates that representations of spatial relations among objects which are not part of the human body are rooted to a large degree in bodily action. However, the large individual differences in spatial aptitude which do not seem to stem from bodily differences demonstrate that we cannot fully bridge the gap between allocentric spatial cognition and the embodied framework.

## Footnotes

1 It has been argued that log-transformed odds are better suited for analysis of percentage values due to floor- and ceiling effects (Jaeger, 2008). Since the results presented here all lie in a range close to 50 % and can be well approximated as normal, we employ linear regression.

2 The convergence issues occurred for both, the balanced dataset of repeatedly measured and the complete set of all participants. That is, the convergence issues were not due to imbalanced data. We also compared the mixed-model algorithms of MATLAB (R2018b) and R (lme4 v1.1-21). Both showed similar convergence issues. Convergence is in fact a common problem in mixed modelling (Eager & Roy, 2017).

