## Supplemental Appendices A-D for "Does Bodily Action Shape Spatial Representation? Evidence from Virtual Reality, Sensory Augmentation and Map Learning"

### Appendix A. Distribution of Participants and Data

#### VR Embodiment

| Overall number of participants | 82 | | |
|---|---|---|---|
| Only participated in first session | 54 | Participated in all three sessions | 28 |
| Data sets | 54 | Participated with complete data sets | 22 |
| | | Participated missing only first session data | 2 |
| | | Missing only second session data | 2 |
| | | Missing only third session data | 2 |
| | | Data Sets | 78 |
| Total number of data sets | 122 | | |
| (optimal number of data sets) | (246) | | |

*Table A1.* Distribution of participants and data sets for the VR condition. One data set includes all results in the spatial tasks but also head position and eye tracking data of the participant recorded during city exploration in one session. Optimally one participant produces one data set for each of the three experimental sessions.

#### VR with Belt Embodiment

| Overall number of participants | 70 | | |
|---|---|---|---|
| Only participated in first session | 43 | Participated in all three sessions | 27 |
| Data sets | 43 | Participated with complete data sets | 23 |
| | | Participated missing only second session data | 1 |
| | | Missing only third session data | 2 |
| | | Missing first and second session data | 1 |
| | | Data Sets | 76 |
| Total number of data sets | 119 | | |
| (optimal number of data sets) | (210) | | |

*Table A2.* Distribution of participants and data sets for the VR with belt condition. One data set includes all results in the spatial tasks but also head position and eye tracking data of the participant recorded during city exploration in one session. Optimally one participant produces one data set for each of the three experimental sessions.

#### Map Embodiment

| Overall number of participants | 74 | | |
|---|---|---|---|
| Only participated in first session | 46 | Participated in all three sessions | 28 |
| Data sets | 46 | Participated with complete data sets | 26 |
| | | Participated missing only third session data | 2 |
| | | Data Sets | 80 |
| Total number of data sets | 126 | | |
| (optimal number of data sets) | (222) | | |

*Table A3.* Distribution of participants and data sets for the map condition. One data set includes all results in the spatial tasks but also mouse (house) clicking data of the participant recorded during city exploration in one session. Optimally one participant produces one data set for each of the three experimental sessions.

#### Appendix B. Comparison of Single Session and Repeated Participants

Here we present a statistical comparison of task performance of participants that only took part in one exploration session and the performance after the first exploration session of longitudinal participants. Because the number of participants that explored the city only once is about double the number of participants that explored the city multiple times we use independent sample t-tests as statistical method of comparison. We compare each sample of task performance of the single-session group to the corresponding sample of the repeated group in their first session. We also compare the marginal samples of the performances over all task conditions to find a possible main effect of single-session- versus longitudinal participant in each embodiment condition. Such a comparison of marginals is important because a lack of single condition effects can hide a main effect.

We find only a significant difference in mean performance in the map exploration condition and there only under time pressure in the Relative task and in the Pointing task. In the Relative task the longitudinal participants perform at 53 percent while the single-session participants reach only 48 percent (p=0.009, t(72)=2.7). In the Pointing task single-session participants perform at 49 percent which is better than the average longitudinal participant who correctly chooses in 44 percent of the cases (p=0.02, t(72)=-2.3). All sample mean performances and standard errors as well as the independent sample t-test results can be found in Tables 1-3. The sample means are illustrated in Figure B.1.

The above findings do not necessarily imply that the single-session participants in the map condition are different from the longitudinal participants. Given that we had no a priori hypotheses for the results of our statistical comparisons and that we performed 18 t-tests at a significance level of five percent each, the probability to find one significant difference by chance alone is 60 percent ($1 - .95^{18}$). That means the probability to find a difference in one of our comparisons, although the samples are from the same population, is higher than not finding one.

We consequently assume that only one of the two significant differences marks a real difference in the underlying population of map participants. Since this difference, in turn, only appears when comparing one specific task condition, we still believe the general assumption reasonable (e.g., for simplicity of the statistical analysis) that the sample of participants who only explored our VR city once and the longitudinal participants both come from the same population.

| One session only | | Absolute | Relative | Pointing |
|---|---|---|---|---|
| VR | 3s | 49.5 ± 1.1 | 47.4 ± 0.9 | 48 ± 1.4 |
| | Unl | 49.7 ± 1.2 | 53.5 ± 1.1 | 52.2 ± 1 |
| Belt | 3s | 46.8 ± 1.2 | 47.5 ± 1.6 | 49.7 ± 1.4 |
| | Unl | 49.5 ± 1.4 | 54.1 ± 1.4 | 51.4 ± 1.5 |
| Map | 3s | 49.9 ± 1.3 | 47.8 ± 1.2 | 49.1 ± 1.4 |
| | Unl | 50.5 ± 1.2 | 54 ± 1.2 | 54.3 ± 1.3 |

*Table B1.* Mean performance plus/minus standard error in percent for all conditions for participants that only explored the VR city (map) once.

| Repeated | | Absolute | Relative | Pointing |
|---|---|---|---|---|
| VR | 3s | 48.2 ± 1.9 | 49.9 ± 1.9 | 48.9 ± 1.5 |
| | Unl | 50.5 ± 1.9 | 52.2 ± 2 | 53.2 ± 1.3 |
| Belt | 3s | 45.7 ± 1.5 | 47.4 ± 1.6 | 52.9 ± 2.3 |
| | Unl | 49.8 ± 1.5 | 50.2 ± 1.4 | 49.8 ± 1.6 |
| Map | 3s | 47.5 ± 1.5 | 52.9 ± 1.4 | 43.7 ± 2 |
| | Unl | 54.8 ± 2 | 55.6 ± 1.5 | 52.8 ± 1.6 |

*Table B2.* Mean performance plus/minus standard error in percent for all conditions after the first exploration session for participants that explored the VR city (map) multiple times.

| Independent t-test | | Absolute | Relative | Pointing | Total marginal |
|---|---|---|---|---|---|
| VR | 3s | t(78)=-0.65, p=0.52 | t(78)=1.34, p=0.18 | t(78)=0.42, p=0.67 | |
| | Unl | t(78)=0.37, p=0.71 | t(78)=-0.6, p=0.55 | t(78)=0.57, p=0.57 | t(478)=0.52, p=0.6 |
| Belt | 3s | t(67)=-0.58, p=0.57 | t(67)=-0.05, p=0.96 | t(67)=1.27, p=0.21 | |
| | Unl | t(67)=0.14, p=0.89 | t(67)=-1.81, p=0.08 | t(67)=-0.7, p=0.49 | t(412)=-0.57, p=0.57 |
| Map | 3s | t(72)=-1.19, p=0.24 | t(72)=2.7, p=0.009 ** | t(72)=-2.3, p=0.02 * | |
| | Unl | t(72)=1.89, p=0.06 | t(72)=0.83, p=0.41 | t(72)=-0.75, p=0.46 | t(442)=0.26, p=0.8 |

*Table B3.* Results of independent t-tests comparing the performance samples for each condition of single-session participants to the first exploration session performance of participants that explored the VR city (map) multiple times. The last column compares the total sample marginalized over all conditions to find a possible main effect of longitudinal participants.

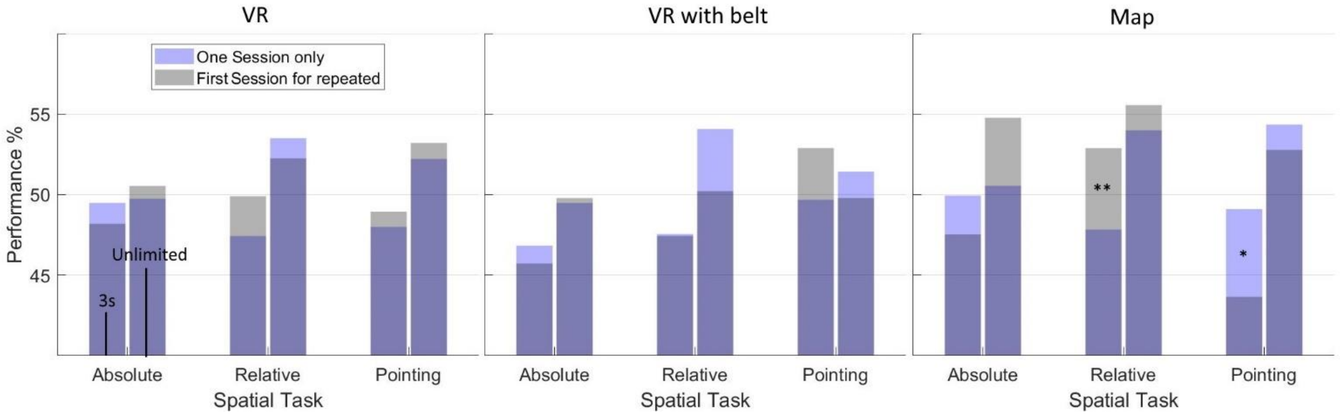

*Figure B1.* (Light blue bars) Barplots of mean performance in all six trial blocks of participants that only explored the city once. (Grey bars) Barplots of mean performance in the first session for participants who explored the city repeatedly (dark blue denotes overlap). Independent sample t-tests show that only in the Map condition comparing Relative task performance under time pressure (p=0.009 **, t(72)=2.7) and when comparing Pointing performance under time pressure (p=0.02 *, t(72)=-2.3) we find a significant difference between single-session and repeated participants in their first session.

#### Appendix C. Plot Results

##### Bar Plot Results

In this section we present the values of mean performance and standard errors in all experimental conditions depicted in the bar plots in Results (Figures 7.1-3, 8.1-3 and 9.1-3). Table C1 details the results in the VR condition (Figures 7.1-3). Table C2 contains the results in the map condition (Figures 8.1-3). Table C3 details the results in the VR condition with belt (Figures 9.1-3).

| Session 1 | Absolute | Relative | Pointing |
|---|---|---|---|
| 3s | 49.1 ± 0.9 | 48.2 ± 0.9 | 48.3 ± 1 |
| Unl | 50 ± 1 | 53.1 ± 1 | 52.5 ± 0.8 |
| Session 2 | | | |
| 3s | 49 ± 2 | 49.6 ± 1.6 | 50.1 ± 1.4 |
| Unl | 55.9 ± 2.1 | 53.1 ± 1.7 | 56.2 ± 1.4 |
| Session 3 | | | |
| 3s | 50.4 ±2 | 50.2±1.6 | 51.8±1.5 |
| Unl | 57.2 ±2.4 | 54.8±2.4 | 59.5±1.5 |

*Table C1.* Mean performance plus/minus standard error in percent in all conditions after VR exploration

| Session 1 | Absolute | Relative | Pointing |
|---|---|---|---|
| 3s | 49 ±1 | 49.7±1 | 47±1.2 |
| Unl | 52.1 ±1.1 | 54.6±0.9 | 53.8±1 |
| Session 2 | | | |
| 3s | 51.1 ±1.8 | 52.5±1.7 | 49.7±1.6 |
| Unl | 62 ±2.9 | 58.2±1.9 | 55.2±1.7 |
| Session 3 | | | |
| 3s | 55.1 ±2.1 | 58.3±2.5 | 49±1.7 |
| Unl | 60.9 ±3.1 | 61±2.4 | 58.1±2.4 |

*Table C2.* Mean performance plus/minus standard error in percent in all conditions after map exploration

| Session 1 | Absolute | Relative | Pointing |
|---|---|---|---|
| 3s | 46.4 ±0.9 | 47.5±1.1 | 50.9±1.2 |
| Unl | 49.6 ±1.1 | 52.6±1.1 | 50.8±1.1 |
| Session 2 | | | |
| 3s | 49.6 ±1.9 | 49.4±2.2 | 49.4±1.7 |
| Unl | 52.2 ±1.4 | 56±1.4 | 55.8±1.4 |
| Session 3 | | | |
| 3s | 50.6 ±1.4 | 51±1.5 | 53.1±2 |
| Unl | 54.1 ±1.8 | 54±1.7 | 53.8±1.4 |

*Table C3.* Mean performance plus/minus standard error in percent in all conditions after VR exploration with belt

##### Model Plot Results

###### Example Model

For readers not familiar with the MATLAB R2018b syntax we give one complete example of the linear mixed-effects model (LMEM) and script structure we used for fitting a single condition across sessions. The example fits the performances in the VR condition, Relative task, unlimited time trial blocks over sessions (green line in Results, Figure 7.6).

First we create a table 'RelInf_Table_VR', holding in the first column all performances in the Relative, unlimited condition 'RelInf_VR' ordered underneath each other. We start with all participants in the first session, then all participants in the second session and finally the third session. The second column holds the coding used for the predictor session 'C_Session_VR'. In the third column we store the respective participant's code number 'C_Subject_VR'. An exemplary depiction of the resulting data structure is given in Table C4.

| RelInf_Table_VR | RelUnl_VR | C_Session_VR | C_Subject_VR |
|---|---|---|---|
| Session 1 | 49 | 0 | 1 |
| | 50 | 0 | 2 |
| | 50 | 0 | 3 |
| Session 2 | 52 | .5 | 1 |
| | 53 | .5 | 3 |
| Session 3 | 54 | 1 | 1 |
| | 56 | 1 | 3 |

*Table C4.* The data structure input to the fitlme() function in MATLAB to compute the LMEM. The first column holds the performances in the Relative unlimited condition. The second column holds the coding used for the predictor session. In the third column we store the respective participant's code number.

Before feeding the table into the LMEM we define the general LMEM structure as:
LMEM_Results_RelInf_VR=fitlme(RelInf_Table_VR, 'RelInf_VR~1+C_Session_VR+(1|C_Subject_VR)')

The model structure is defined through the formula bracketed by the apostrophes (notation is similar to Wilkinson and Rogers, 1973). The term '(1|C_Subject_VR)' specifies the random effects model- and covariance matrix. The sparse number of single condition trials/observations in session two and three allows only a fit of one parameter for each subject. Consequently, the covariance matrix becomes a scalar. We decided for intercept as random effect because it leads to an overall better fit than slope. The model results are given in Table C5.

| | | | | | |
|---|---|---|---|---|---|
| Degrees of freedom (df) | #obs \| #fix \| #rand \| #cov | 132 | 2 | 82 | 2 |
| Fixed effects | β0(Intercept) ± SE \| CI | 53.01 ± 1.02 | 50.98 55.04 | | |
| | β1(Session) ± SE \| CI \| χ2(1) \| p | 1.6 ± 2.01 | -2.38 5.57 | 0.63 | 0.43 |
| Random effects | Intercept σ \| CI | 3.32 | 1.44 7.64 | | |
| | Residual variance \| CI | 8.81 | 7.52 10.33 | | |

*Table C5.* Results of the LMEM fit over sessions of the Relative task, unlimited time condition in the VR exploration condition. #obs means total number of experimental observations. #fix means number of fixed effect coefficients. #rand means total number of random effect coefficients. #cov means number of covariance parameters. SE is standard error and CI 95% confidence interval borders. $\chi2(1)$ is the likelihood ratio statistic with 1 degree of freedom difference to the model without this fixed effect. σ is the random effect standard deviation.

**VR Exploration Model Plots**

We provide details on the models we used for creating the VR exploration condition linear fit plots in Results (Figures 7.4-6). The fixed effect design matrix is identical across the single-condition-over-session plots for all experiments. This also holds for the time plots that marginalize over the tree spatial tasks (Figure 7.4). The design matrix is shown in Table C6.

Table C7 holds model results for the three second condition, marginalizing spatial tasks (Figure 7.4). Table C8, C9 and C10 show results for the Absolute -, Relative - and Pointing task, respectively. All computed for the three second condition (Figure 7.5).

Table C11 shows the model for the unlimited condition, marginalizing spatial tasks (Figure 7.4). Table C12, C13 and C14 depict results for the Absolute -, Relative - and Pointing task, respectively. Here computed for the unlimited condition (Figure 7.6).

| Intercept | Session |
|---|---|
| 1 | 0 |
| 1 | .5 |
| 1 | 1 |

*Table C6.* The fixed effect design matrix used in all plot LMEMs across experiments.

| | | | | | |
|---|---|---|---|---|---|
| Degrees of freedom (df) | #obs \| #fix \| #rand \| #cov | 396 | 2 | 164 | 3 |
| Fixed effects | β0(Intercept) ± SE \| CI | 48.53 ± 0.54 | 47.46 49.59 | | |
| | β1(Session) ± SE \| CI \| χ2(1) \| p | 2.06 ± 1.3 | -0.49 4.61 | 2.42 | 0.12 |
| Random effects | Intercept σ \| CI | 1.83 | 0.69 4.84 | | |
| | Session σ \| CI | 3.89 | 1.8 8.4 | | |
| | Residual variance \| CI | 8.01 | 7.42 8.64 | | |

*Table C7.* Results of the LMEM fit over sessions marginalizing over all three spatial tasks but only the 3s time conditions in the VR exploration condition. #obs means total number of experimental observations. #fix means number of fixed effect coefficients. #rand means total number of random effect coefficients. #cov means number of covariance parameters. SE is standard error and CI 95% confidence interval borders. χ2(1) is the likelihood ratio statistic with 1 degree of freedom difference to the model without this fixed effect. σ is the random effect standard deviation.
The LMEM formula is: 'CondResult~1+Session+(1+Session|Subject)'.

| | | | | | |
|---|---|---|---|---|---|
| Degrees of freedom (df) | #obs \| #fix \| #rand \| #cov | 132 | 2 | 82 | 2 |
| Fixed effects | β0(Intercept) ± SE \| CI | 48.99 ± 0.97 | 47.06 50.92 | | |
| | β1(Session) ± SE \| CI \| χ2(1) \| p | 1.35 ± 1.8 | -2.31 5 | 0.53 | 0.47 |
| Random effects | Intercept σ \| CI | 4.26 | 2.58 7.04 | | |
| | Residual variance \| CI | 7.86 | 6.66 9.27 | | |

*Table C8.* Results of the LMEM fit over sessions of the 3s Absolute task condition in the VR exploration condition.
#obs means total number of experimental observations. #fix means number of fixed effect coefficients. #rand means total number of random effect coefficients. #cov means number of covariance parameters. SE is standard error and CI 95% confidence interval borders. χ2(1) is the likelihood ratio statistic with 1 degree of freedom difference to the model without this fixed effect. σ is the random effect standard deviation.
The LMEM formula is: 'CondResult~1+Session+(1|Subject)'.

| | | | | | |
|---|---|---|---|---|---|
| Degrees of freedom (df) | #obs \| #fix \| #rand \| #cov | 132 | 2 | 82 | 2 |
| Fixed effects | β0(Intercept) ± SE \| CI | 48.37 ± 0.86 | 46.67 50.07 | | |
| | β1(Session) ± SE \| CI \| χ2(1) \| p | 1.14 ± 1.56 | -1.95 4.22 | 0.51 | 0.47 |
| Random effects | Intercept σ \| CI | 4.48 | 3.05 6.57 | | |
| | Residual variance \| CI | 6.45 | 5.42 7.66 | | |

*Table C9.* Results of the LMEM fit over sessions of the 3s Relative task condition in the VR exploration condition.
#obs means total number of experimental observations. #fix means number of fixed effect coefficients. #rand means total number of random effect coefficients. #cov means number of covariance parameters. SE is standard error and CI 95% confidence interval borders. χ2(1) is the likelihood ratio statistic with 1 degree of freedom difference to the model without this fixed effect. σ is the random effect standard deviation.
The LMEM formula is: 'CondResult~1+Session+(1|Subject)'.

| | | | | | |
|---|---|---|---|---|---|
| Degrees of freedom (df) | #obs \| #fix \| #rand \| #cov | 132 | 2 | 0 | 1 |
| Fixed effects | β0(Intercept) ± SE \| CI | 48.31 ± 0.91 | 46.5 50.12 | | |
| | β1(Session) ± SE \| CI \| χ2(1) \| p | 3.53 ± 1.84 | -0.12 7.18 | 3.62 | 0.057 |
| Random effects | Residual variance \| CI | 8.44 | 7.48 9.53 | | |

*Table C10***.** Results of the LMEM fit over sessions of the 3s Pointing task condition in the VR exploration condition.
#obs means total number of experimental observations. #fix means number of fixed effect coefficients. #rand means total number of random effect coefficients. #cov means number of covariance parameters. SE is standard error and CI 95% confidence interval borders. χ2(1) is the likelihood ratio statistic with 1 degree of freedom difference to the model without this fixed effect. σ is the random effect standard deviation. Here the random effect intercept σ was close to zero ($2*10^{-6}$) which leads to numerical instabilities and means one can and should ignore this effect. We therefore did not include it in the model.
The LMEM formula is: 'CondResult~1+Session'.
(We could have used a simple linear model, but results are similar and staying in LMEM analysis style is simpler).

| | | | | | |
|---|---|---|---|---|---|
| Degrees of freedom (df) | #obs \| #fix \| #rand \| #cov | 396 | 2 | 82 | 2 |
| Fixed effects | β0(Intercept) ± SE \| CI | 51.94 ± 0.54 | 50.89 52.99 | | |
| | β1(Session) ± SE \| CI \| χ2(1) \| p | 5.22 ± 1.56 | 2.14 8.29 | 9.51 | 0.002 |
| Random effects | Session σ \| CI | 5.86 | 3.81 9 | | |
| | Residual variance \| CI | 8.57 | 8 9.21 | | |

*Table C11.* Results of the LMEM fit over sessions marginalizing over all three spatial tasks but only the unlimited time conditions in the VR exploration condition. #obs means total number of experimental observations. #fix means number of fixed effect coefficients. #rand means total number of random effect coefficients. #cov means number of covariance parameters. SE is standard error and CI 95% confidence interval borders. χ2(1) is the likelihood ratio statistic with 1 degree of freedom difference to the model without this fixed effect. σ is the random effect standard deviation. Here the random effect intercept σ was close to zero (0.008) which leads to numerical instabilities and means one can and should ignore this effect. We therefore did not include it in the model.
The LMEM formula is: 'CondResult~1+Session+(-1+Session|Subject)'.

| | | | | | |
|---|---|---|---|---|---|
| Degrees of freedom (df) | #obs \| #fix \| #rand \| #cov | 132 | 2 | 82 | 2 |
| Fixed effects | β0(Intercept) ± SE \| CI | 50.24 ± 1.07 | 48.13 52.36 | | |
| | β1(Session) ± SE \| CI \| χ2(1) \| p | 7.23 ± 2.03 | 3.22 11.24 | 12.14 | 0.0005 |
| Random effects | Intercept σ \| CI | 4.69 | 2.84 7.77 | | |
| | Residual variance \| CI | 8.62 | 7.3 10.17 | | |

*Table C12.* Results of the LMEM fit over sessions of the unlimited Absolute task condition in the VR exploration condition.
#obs means total number of experimental observations. #fix means number of fixed effect coefficients. #rand means total number of random effect coefficients. #cov means number of covariance parameters. SE is standard error and CI 95% confidence interval borders. χ2(1) is the likelihood ratio statistic with 1 degree of freedom difference to the model without this fixed effect. σ is the random effect standard deviation.
The LMEM formula is: 'CondResult~1+Session+(1|Subject)'.

| | | | | | |
|---|---|---|---|---|---|
| Degrees of freedom (df) | #obs \| #fix \| #rand \| #cov | 132 | 2 | 82 | 2 |
| Fixed effects | β0(Intercept) ± SE \| CI | 53.01 ± 1.02 | 50.98 55.04 | | |
| | β1(Session) ± SE \| CI \| χ2(1) \| p | 1.6 ± 2.01 | -2.38 5.57 | 0.63 | 0.43 |
| Random effects | Intercept σ \| CI | 3.32 | 1.44 7.64 | | |
| | Residual variance \| CI | 8.81 | 7.52 10.33 | | |

*Table C13.* Results of the LMEM fit over sessions of the unlimited Relative task condition in the VR exploration condition.
#obs means total number of experimental observations. #fix means number of fixed effect coefficients. #rand means total number of random effect coefficients. #cov means number of covariance parameters. SE is standard error and CI 95% confidence interval borders. χ2(1) is the likelihood ratio statistic with 1 degree of freedom difference to the model without this fixed effect. σ is the random effect standard deviation. The LMEM formula is: 'CondResult~1+Session+(1|Subject)'.

| | | | | | |
|---|---|---|---|---|---|
| Degrees of freedom (df) | #obs \| #fix \| #rand \| #cov | 132 | 2 | 82 | 2 |
| Fixed effects | β0(Intercept) ± SE \| CI | 52.49 ± 0.8 | 50.92 54.07 | | |
| | β1(Session) ± SE \| CI \| χ2(1) \| p | 6.91 ± 1.5 | 3.93 9.89 | 19.38 | 0.00001 |
| Random effects | Intercept σ \| CI | 3.55 | 1.98 6.36 | | |
| | Residual variance \| CI | 6.38 | 5.31 7.66 | | |

*Table C14.* Results of the LMEM fit over sessions of the unlimited Pointing task condition in the VR exploration condition.
#obs means total number of experimental observations. #fix means number of fixed effect coefficients. #rand means total number of random effect coefficients. #cov means number of covariance parameters. SE is standard error and CI 95% confidence interval borders. χ2(1) is the likelihood ratio statistic with 1 degree of freedom difference to the model without this fixed effect. σ is the random effect standard deviation. The LMEM formula is: 'CondResult~1+Session+(1|Subject)'.

**Map Exploration Model Plots**

We provide details on the models we used for creating the map exploration condition linear fit plots in Results (Figures 8.4-6). Table C15 holds model results for the three second condition, marginalizing spatial tasks (Figure 8.4). Table C16, C17 and C18 show results for the Absolute -, Relative - and Pointing task, respectively. All computed for the three second condition (Figure 8.5).

Table C19 shows the model for the unlimited condition, marginalizing spatial tasks (Figure 8.4). Table C20, C21 and C22 depict results for the Absolute -, Relative - and Pointing task, respectively. Here computed for the unlimited condition (Figure 8.6).

| | | | | | |
|---|---|---|---|---|---|
| Degrees of freedom (df) | #obs \| #fix \| #rand \| #cov | 384 | 2 | 148 | 3 |
| Fixed effects | β0(Intercept) ± SE \| CI | 48.55 ± 0.63 | 47.32 49.78 | | |
| | β1(Session) ± SE \| CI \| χ2(1) \| p | 5.79 ± 1.62 | 2.61 8.97 | 11.06 | 0.00088 |
| Random effects | Intercept σ \| CI | 2.21 | 0.91 5.39 | | |
| | Session σ \| CI | 5.75 | 3.52 9.4 | | |

| | | | |
|---|---|---|---|
| | Residual variance \| CI | 8.82 | 8.14<br>9.56 |

*Table C15.* Results of the LMEM fit over sessions marginalizing over all three spatial tasks but only the 3s time conditions in the VR exploration condition. #obs means total number of experimental observations. #fix means number of fixed effect coefficients. #rand means total number of random effect coefficients. #cov means number of covariance parameters. SE is standard error and CI 95% confidence interval borders. χ2(1) is the likelihood ratio statistic with 1 degree of freedom difference to the model without this fixed effect. σ is the random effect standard deviation.
The LMEM formula is: 'CondResult~1+Session+(1+Session |Subject)'.

| | | | | | |
|---|---|---|---|---|---|
| Degrees of freedom (df) | #obs \| #fix \| #rand \| #cov | 128 | 2 | 74 | 2 |
| Fixed effects | β0(Intercept) ± SE \| CI | 48.9<br>± 1.04 | 46.85<br>50.95 | | |
| | β1(Session) ± SE \| CI \| χ2(1) \| p | 6.17<br>± 1.97 | 2.27<br>10.06 | 9.04 | 0.0026 |
| Random effects | Intercept σ \| CI | 3.26 | 1.27<br>8.38 | | |
| | Residual variance \| CI | 8.59 | 7.26<br>10.17 | | |

*Table C16.* Results of the LMEM fit over sessions of the 3s Absolute task condition in the map exploration condition.
#obs means total number of experimental observations. #fix means number of fixed effect coefficients. #rand means total number of random effect coefficients. #cov means number of covariance parameters. SE is standard error and CI 95% confidence interval borders. χ2(1) is the likelihood ratio statistic with 1 degree of freedom difference to the model without this fixed effect. σ is the random effect standard deviation
The LMEM formula is: 'CondResult~1+Session+(1|Subject)'.

| | | | | | |
|---|---|---|---|---|---|
| Degrees of freedom (df) | #obs \| #fix \| #rand \| #cov | 128 | 2 | 74 | 2 |
| Fixed effects | β0(Intercept) ± SE \| CI | 49.45<br>± 1.05 | 47.37<br>51.54 | | |
| | β1(Session) ± SE \| CI \| χ2(1) \| p | 7.24<br>± 1.82 | 3.64<br>10.84 | 14.96 | 0.00011 |
| Random effects | Intercept σ \| CI | 5.45 | 3.8<br>7.83 | | |
| | Residual variance \| CI | 7.5 | 6.33<br>8.87 | | |

*Table C17.* Results of the LMEM fit over sessions of the 3s Relative task condition in the map exploration condition.
#obs means total number of experimental observations. #fix means number of fixed effect coefficients. #rand means total number of random effect coefficients. #cov means number of covariance parameters. SE is standard error and CI 95% confidence interval borders. χ2(1) is the likelihood ratio statistic with 1 degree of freedom difference to the model without this fixed effect. σ is the random effect standard deviation.
The LMEM formula is: 'CondResult~1+Session+(1|Subject)'.

| | | | | | |
|---|---|---|---|---|---|
| Degrees of freedom (df) | #obs \| #fix \| #rand \| #cov | 128 | 2 | 74 | 2 |
| Fixed effects | β0(Intercept) ± SE \| CI | 47.27<br>± 1.06 | 45.16<br>49.37 | | |
| | β1(Session) ± SE \| CI \| χ2(1) \| p | 2.52<br>± 2.08 | -1.61<br>6.64 | 3.62 | 0.057 |
| Random effects | Intercept σ \| CI | 1.32 | 0<br>525.94 | | |

| | | | |
|---|---|---|---|
| | Residual variance \| CI | 9.38 | 7.93<br>11.1 |

*Table C18.* Results of the LMEM fit over sessions of the 3s Pointing task condition in the map exploration condition.
#obs means total number of experimental observations. #fix means number of fixed effect coefficients. #rand means total number of random effect coefficients. #cov means number of covariance parameters. SE is standard error and CI 95% confidence interval borders. χ2(1) is the likelihood ratio statistic with 1 degree of freedom difference to the model without this fixed effect. σ is the random effect standard deviation. Here the random effect intercept σ was very close to zero which leads to numerical instabilities and means one can and should ignore this effect. We therefore did not include. We therefore did not include it in the model.
The LMEM formula is: 'CondResult~1+Session'.
(We could have used a simple linear model, but results are similar and staying in LMEM analysis style is simpler).

| Degrees of freedom (df) | #obs \| #fix \| #rand \| #cov | 384 | 2 | 148 | 3 |
|---|---|---|---|---|---|
| Fixed effects | β0(Intercept) ± SE \| CI | 53.65<br>± 0.66 | 52.36<br>54.94 | | |
| | β1(Session) ± SE \| CI \| χ2(1) \| p | 7.31<br>± 2.21 | 2.97<br>11.64 | 9.61 | 0.0019 |
| Random effects | Intercept σ \| CI | 2.94 | 1.66<br>5.21 | | |
| | Session σ \| CI | 9.71 | 6.64<br>14.21 | | |
| | Residual variance \| CI | 8.63 | 7.98<br>9.34 | | |

*Table C19.* Results of the LMEM fit over sessions marginalizing over all three spatial tasks but only the unlimited time conditions in the map exploration condition. #obs means total number of experimental observations. #fix means number of fixed effect coefficients. #rand means total number of random effect coefficients. #cov means number of covariance parameters. SE is standard error and CI 95% confidence interval borders. χ2(1) is the likelihood ratio statistic with 1 degree of freedom difference to the model without this fixed effect. σ is the random effect standard deviation.
The LMEM formula is: 'CondResult~1+Session+(1+Session|Subject)'.

| Degrees of freedom (df) | #obs \| #fix \| #rand \| #cov | 128 | 2 | 74 | 2 |
|---|---|---|---|---|---|
| Fixed effects | β0(Intercept) ± SE \| CI | 52.73<br>± 1.36 | 50.03<br>55.44 | | |
| | β1(Session) ± SE \| CI \| χ2(1) \| p | 8.49<br>± 2.07 | 4.38<br>12.59 | 15.7 | 0.00007 |
| Random effects | Intercept σ \| CI | 8.63 | 6.76<br>11.03 | | |
| | Residual variance \| CI | 8.23 | 6.95<br>9.74 | | |

*Table C20.* Results of the LMEM fit over sessions of the unlimited Absolute task condition in the map exploration condition.
#obs means total number of experimental observations. #fix means number of fixed effect coefficients. #rand means total number of random effect coefficients. #cov means number of covariance parameters. SE is standard error and CI 95% confidence interval borders. χ2(1) is the likelihood ratio statistic with 1 degree of freedom difference to the model without this fixed effect. σ is the random effect standard deviation.
The LMEM formula is: 'CondResult~1+Session+(1|Subject)'.

| Degrees of freedom (df) | #obs \| #fix \| #rand \| #cov | 128 | 2 | 74 | 2 |
|---|---|---|---|---|---|
| Fixed effects | β0(Intercept) ± SE \| CI | 54.61<br>± 1.04 | 52.56<br>56.66 | | |

| | | | | | |
|---|---|---|---|---|---|
| | β1(Session) ± SE \| CI \| χ2(1) \| p | 6.44<br>± 1.88 | 2.72<br>10.16 | 11.18 | 0.00083 |
| Random effects | Intercept σ \| CI | 4.54 | 2.83<br>7.31 | | |
| | Residual variance \| CI | 7.94 | 6.73<br>9.37 | | |

*Table C21.* Results of the LMEM fit over sessions of the unlimited Relative task condition in the map exploration condition.
#obs means total number of experimental observations. #fix means number of fixed effect coefficients. #rand means total number of random effect coefficients. #cov means number of covariance parameters. SE is standard error and CI 95% confidence interval borders. χ2(1) is the likelihood ratio statistic with 1 degree of freedom difference to the model without this fixed effect. σ is the random effect standard deviation.
The LMEM formula is: 'CondResult~1+Session+(1|Subject)'.

| Degrees of freedom (df) | #obs \| #fix \| #rand \| #cov | 128 | 2 | 74 | 2 |
|---|---|---|---|---|---|
| Fixed effects | β0(Intercept) ± SE \| CI | 53.66<br>± 1.07 | 51.54<br>55.78 | | |
| | β1(Session) ± SE \| CI \| χ2(1) \| p | 4.4<br>± 2 | 0.43<br>8.37 | 4.64 | 0.031 |
| Random effects | Intercept σ \| CI | 3.93 | 1.98<br>7.81 | | |
| | Residual variance \| CI | 8.64 | 7.32<br>10.21 | | |

*Table C22.* Results of the LMEM fit over sessions of the unlimited Pointing task condition in the map exploration condition.
#obs means total number of experimental observations. #fix means number of fixed effect coefficients. #rand means total number of random effect coefficients. #cov means number of covariance parameters. SE is standard error and CI 95% confidence interval borders. χ2(1) is the likelihood ratio statistic with 1 degree of freedom difference to the model without this fixed effect. σ is the random effect standard deviation.
The LMEM formula is: 'CondResult~1+Session+(1|Subject)'.

**VR with Belt Exploration Model Plots**

We provide details on the models we used for creating the VR with belt exploration condition linear fit plots in Results (Figures 9.4-6). Table C23 holds model results for the three second condition, marginalizing spatial tasks (Figure 9.4). Table C24, C25 and C26 show results for the Absolute -, Relative - and Pointing task, respectively. All computed for the three second condition (Figure 9.5).

Table C27 shows the model for the unlimited condition, marginalizing spatial tasks (Figure 9.4). Table C28, C29 and C30 depict results for the Absolute -, Relative - and Pointing task, respectively. Here computed for the unlimited condition (Figure 9.6).

| Degrees of freedom (df) | #obs \| #fix \| #rand \| #cov | 357 | 2 | 70 | 2 |
|---|---|---|---|---|---|
| Fixed effects | β0(Intercept) ± SE \| CI | 48.19<br>± 0.67 | 46.88<br>49.51 | | |
| | β1(Session) ± SE \| CI \| χ2(1) \| p | 3.07<br>± 1.22 | 0.67<br>5.46 | 6.25 | 0.012 |
| Random effects | Intercept σ \| CI | 2.65 | 1.5<br>4.7 | | |
| | Residual variance \| CI | 8.75 | 8.07<br>9.49 | | |

*Table C23.* Results of the LMEM fit over sessions marginalizing over all three spatial tasks but only the 3s time conditions in the VR with belt exploration condition. #obs means total number of experimental observations. #fix means number of fixed effect coefficients. #rand means total number of random effect coefficients. #cov means number of covariance parameters. SE is standard error and CI 95% confidence interval borders. χ2(1) is the likelihood ratio statistic with 1 degree of freedom difference to the model without this fixed effect. σ is the random effect standard deviation. Here the random effect session σ was close to zero (0.01) which leads to numerical instabilities and means one can and should ignore this effect. We therefore did not include it in the model.
The LMEM formula is: 'CondResult~1+Session+(1|Subject)'.

| | | | | | |
|---|---|---|---|---|---|
| Degrees of freedom (df) | #obs \| #fix \| #rand \| #cov | 119 | 2 | 70 | 2 |
| Fixed effects | β0(Intercept) ± SE \| CI | 46.56 ± 0.93 | 44.7 48.4 | | |
| | β1(Session) ± SE \| CI \| χ2(1) \| p | 4.49 ± 1.7 | 1.13 7.86 | 6.73 | 0.0095 |
| Random effects | Intercept σ \| CI | 3.71 | 2.05 6.71 | | |
| | Residual variance \| CI | 7.03 | 5.89 8.41 | | |

*Table C24.* Results of the LMEM fit over sessions of the 3s Absolute task condition in the VR with belt exploration condition.
#obs means total number of experimental observations. #fix means number of fixed effect coefficients. #rand means total number of random effect coefficients. #cov means number of covariance parameters. SE is standard error and CI 95% confidence interval borders. χ2(1) is the likelihood ratio statistic with 1 degree of freedom difference to the model without this fixed effect. σ is the random effect standard deviation
The LMEM formula is: 'CondResult~1+Session+(1|Subject)'.

| | | | | | |
|---|---|---|---|---|---|
| Degrees of freedom (df) | #obs \| #fix \| #rand \| #cov | 119 | 2 | 70 | 2 |
| Fixed effects | β0(Intercept) ± SE \| CI | 47.52 ± 1.05 | 45.35 49.68 | | |
| | β1(Session) ± SE \| CI \| χ2(1) \| p | 3.61 ± 1.95 | -0.25 7.47 | 3.35 | 0.067 |
| Random effects | Intercept σ \| CI | 4.79 | 2.69 8.54 | | |
| | Residual variance \| CI | 7.99 | 6.57 9.71 | | |

*Table C25.* Results of the LMEM fit over sessions of the 3s Relative task condition in the VR with belt exploration condition.
#obs means total number of experimental observations. #fix means number of fixed effect coefficients. #rand means total number of random effect coefficients. #cov means number of covariance parameters. SE is standard error and CI 95% confidence interval borders. χ2(1) is the likelihood ratio statistic with 1 degree of freedom difference to the model without this fixed effect. σ is the random effect standard deviation.
The LMEM formula is: 'CondResult~1+Session+(1|Subject)'.

| | | | | | |
|---|---|---|---|---|---|
| Degrees of freedom (df) | #obs \| #fix \| #rand \| #cov | 119 | 2 | 70 | 2 |
| Fixed effects | β0(Intercept) ± SE \| CI | 50.52 ± 1.14 | 48.27 52.78 | | |
| | β1(Session) ± SE \| CI \| χ2(1) \| p | 1.31 ± 2.14 | -2.94 5.55 | 0.37 | 0.55 |
| Random effects | Intercept σ \| CI | 3.46 | 1.33 9.02 | | |
| | Residual variance \| CI | 9.1 | 7.65 10.83 | | |

*Table C26.* Results of the LMEM fit over sessions of the 3s Pointing task condition in the VR with belt exploration condition.
#obs means total number of experimental observations. #fix means number of fixed effect coefficients. #rand means total number of random effect coefficients. #cov means number of covariance parameters. SE is standard error and CI 95% confidence interval borders. χ2(1) is the likelihood ratio statistic with 1 degree of freedom difference to the model without this fixed effect. σ is the random effect standard deviation.
Here the random effect intercept σ was very close to zero which leads to numerical instabilities and means one should (and can) ignore this effect. We therefore did not include it in the model.
The LMEM formula is: 'CondResult~1+Session'.
(We could have used a simple linear model, but results are basically identical and staying in LMEM analysis style was considered less confusing).

| | | | | | |
|---|---|---|---|---|---|
| Degrees of freedom (df) | #obs \| #fix \| #rand \| #cov | 357 | 2 | 70 | 2 |
| Fixed effects | β0(Intercept) ± SE \| CI | 51.26 ± 0.69 | 49.91 52.62 | | |
| | β1(Session) ± SE \| CI \| χ2(1) \| p | 3.89 ± 1.12 | 1.68 6.09 | 8.87 | 0.0029 |
| Random effects | Intercept σ \| CI | 3.8 | 2.7 5.34 | | |
| | Residual variance \| CI | 7.64 | 7.03 8.3 | | |

*Table C27.* Results of the LMEM fit over sessions marginalizing over all three spatial tasks but only the unlimited time conditions in the VR with belt exploration condition. #obs means total number of experimental observations. #fix means number of fixed effect coefficients. #rand means total number of random effect coefficients. #cov means number of covariance parameters. SE is standard error and CI 95% confidence interval borders. χ2(1) is the likelihood ratio statistic with 1 degree of freedom difference to the model without this fixed effect. σ is the random effect standard deviation. Here the random effect session σ was close to zero (0.01) which leads to numerical instabilities and means one can and should ignore this effect. We therefore did not include it in the model.
The LMEM formula is: 'CondResult~1+Session+(1|Subject)'.

| | | | | | |
|---|---|---|---|---|---|
| Degrees of freedom (df) | #obs \| #fix \| #rand \| #cov | 119 | 2 | 70 | 2 |
| Fixed effects | β0(Intercept) ± SE \| CI | 49.62 ± 0.98 | 47.67 51.57 | | |
| | β1(Session) ± SE \| CI \| χ2(1) \| p | 4.56 ± 1.74 | 1.11 8.01 | 6.62 | 0.01 |
| Random effects | Intercept σ \| CI | 4.46 | 2.58 7.71 | | |
| | Residual variance \| CI | 7.09 | 5.82 8.64 | | |

*Table C28.* Results of the LMEM fit over sessions of the unlimited Absolute task condition in the VR with belt exploration condition.
#obs means total number of experimental observations. #fix means number of fixed effect coefficients. #rand means total number of random effect coefficients. #cov means number of covariance parameters. SE is standard error and CI 95% confidence interval borders. χ2(1) is the likelihood ratio statistic with 1 degree of freedom difference to the model without this fixed effect. σ is the random effect standard deviation.
The LMEM formula is: 'CondResult~1+Session+(1|Subject)'.

| | | | | | |
|---|---|---|---|---|---|
| Degrees of freedom (df) | #obs \| #fix \| #rand \| #cov | 119 | 2 | 70 | 2 |
| Fixed effects | β0(Intercept) ± SE \| CI | 53 ± 0.98 | 51.06 54.92 | | |
| | β1(Session) ± SE \| CI \| χ2(1) \| p | 2.44 ± 1.8 | -1.13 6.01 | 1.73 | 0.19 |
| Random effects | Intercept σ \| CI | 3.55 | 1.5 8.42 | | |
| | Residual variance \| CI | 7.54 | 6.18 9.2 | | |

*Table A29.* Results of the LMEM fit over sessions of the unlimited Relative task condition in the VR with belt exploration condition.
#obs means total number of experimental observations. #fix means number of fixed effect coefficients. #rand means total number of random effect coefficients. #cov means number of covariance parameters. SE is standard error and CI 95% confidence interval borders. χ2(1) is the likelihood ratio statistic with 1 degree of freedom difference to the model without this fixed effect. σ is the random effect standard deviation.
The LMEM formula is: 'CondResult~1+Session+(1|Subject)'.

| | | | | | |
|---|---|---|---|---|---|
| Degrees of freedom (df) | #obs \| #fix \| #rand \| #cov | 119 | 2 | 70 | 2 |
| Fixed effects | β0(Intercept) ± SE \| CI | 51.22 ± 1.01 | 49.22 53.22 | | |
| | β1(Session) ± SE \| CI \| χ2(1) \| p | 3.97 ± 1.83 | 0.34 7.6 | 4.47 | 0.034 |
| Random effects | Intercept σ \| CI | 4.05 | 1.76 9.31 | | |
| | Residual variance \| CI | 7.59 | 6.08 9.46 | | |

*Table C30.* Results of the LMEM fit over sessions of the unlimited Pointing task condition in the VR with belt exploration condition. #obs means total number of experimental observations. #fix means number of fixed effect coefficients. #rand means total number of random effect coefficients. #cov means number of covariance parameters. SE is standard error and CI 95% confidence interval borders. χ2(1) is the likelihood ratio statistic with 1 degree of freedom difference to the model without this fixed effect. σ is the random effect standard deviation. The LMEM formula is: 'CondResult~1+Session+(1|Subject)'.

#### Appendix D. Best-fit Model Results

In this section of the Appendix we provide detailed results of all best-fit models. That are the models selected according to the step-down procedure described in Methods. We also provide information on the maximum models, before step-down. For an introductory example of how to set up a linear mixed effect model (LMEM) in MATLAB, please see above.

##### Best-fit VR Models

We provide details on all linear models used for the quantitative analysis of the VR exploration condition as presented in Results. Table D1 and D2 hold model results and design matrix, respectively, of the fit over session, task and time. Table D3 and D4 depict results and design matrix of the fit over session, task and three seconds decision time. Table D5 and D6 contain results and design matrix of the fit over session, task and unlimited decision time.

| | | | | | |
|---|---|---|---|---|---|
| Degrees of freedom (df) | #obs \| #fix \| #rand \| #cov | 792 | 4 | 164 | 3 |
| Fixed effects | β0(Intercept) ± SE \| CI | 52.01 ± 0.48 | 51.08 52.94 | | |
| | β1(Session) ± SE \| CI \| χ2(1) \| p | 3.57 ± 0.83 | 1.94 5.2 | 18.02 | 0.000022 |
| | β2(Time) ± SE \| CI \| χ2(1) \| p | 5 ± 0.67 | 1.94 5.2 | 41.13 | 1.4*10⁻10 |
| | β1*β2 ± SE \| CI \| χ2(1) \| p | 3.16 ± 1.49 | 0.23 6.08 | 4.45 | 0.035 |
| Random effects | Intercept σ \| CI | 2.52 | 1.85 3.43 | | |
| | Time σ \| CI | 0.79 | 0 321.05 | | |
| | Residual variance \| CI | 8.35 | 7.91 8.8 | | |

*Table D1.* Model a-VR.
Results of the best LMEM fit of the VR exploration condition over session, task and time. #obs means total number of experimental observations. #fix means number of fixed effect coefficients. #rand means total number of random effect coefficients. #cov means number of covariance parameters. SE is standard error and CI 95% confidence interval borders. χ2(1) is the likelihood ratio statistic with 1 degree of freedom difference to the model without this fixed effect. σ is the random effect standard deviation. Here the random effect session σ was close to zero (0.02) which leads to numerical instabilities and means one can and should ignore this effect. It is therefore not included in the best fit model.
The best fit LMEM formula is: 'CondResult~1+Time+Session+Time*Session+(1+Time|Subject)'.
In short: 'CondResult~Time*Session+(1+Time|Subject)'.

| Condition | β0(Intercept) | β1(Session) | β2(Time) | β3(Mean to Rel) | β4(Mean to Poi) | β1(Session)* β2(Time) |
|---|---|---|---|---|---|---|
| S1 Abs 3s | 1 | -.5 | -.5 | -1 | -1 | .25 |
| S1 Rel 3s | 1 | -.5 | -.5 | 1 | 0 | .25 |
| S1 Poi 3s | 1 | -.5 | -.5 | 0 | 1 | .25 |
| S1 Abs Unl | 1 | -.5 | .5 | -1 | -1 | -.25 |
| S1 Rel Unl | 1 | -.5 | .5 | 1 | 0 | -.25 |
| S1 Poi Unl | 1 | -.5 | .5 | 0 | 1 | -.25 |
| S2 Abs 3s | 1 | 0 | -.5 | -1 | -1 | 0 |
| S2 Rel 3s | 1 | 0 | -.5 | 1 | 0 | 0 |
| S2 Poi 3s | 1 | 0 | -.5 | 0 | 1 | 0 |
| S2 Abs Unl | 1 | 0 | .5 | -1 | -1 | 0 |
| S2 Rel Unl | 1 | 0 | .5 | 1 | 0 | 0 |
| S2 Poi Unl | 1 | 0 | .5 | 0 | 1 | 0 |
| S3 Abs 3s | 1 | .5 | -.5 | -1 | -1 | -.25 |
| S3 Rel 3s | 1 | .5 | -.5 | 1 | 0 | -.25 |
| S3 Poi 3s | 1 | .5 | -.5 | 0 | 1 | -.25 |
| S3 Abs Unl | 1 | .5 | .5 | -1 | -1 | .25 |
| S3 Rel Unl | 1 | .5 | .5 | 1 | 0 | .25 |
| S3 Poi Unl | 1 | .5 | .5 | 0 | 1 | .25 |

*Table D2.* Model a-VR Matrix.
The fixed effect design matrix used to fit the VR exploration condition over session, task and time. The grey columns denote all main or simple effect predictors and their matrix coding, which dropped out in the process of model selection. Note that the maximum model before step down selection included all predictors depicted here as well as all their interactions as fixed effects. The maximum model also included all fixed effect terms except interactions as random effects.
The maximum LMEM formula is:
'CondResult~Time*Session*MeanRel+Time*Session*MeanPoi+(1+Time+Session+MeanRel+MeanPoi | Subject)'.

| | | | | | |
|---|---|---|---|---|---|
| Degrees of freedom (df) | #obs \| #fix \| #rand \| #cov | 396 | 4 | 164 | 3 |
| Fixed effects | β0(Intercept) ± SE \| CI | 49.55 ± 0.58 | 48.41 50.69 | | |
| | β1(Session) ± SE \| CI \| χ2(1) \| p | 2.03 ± 1.09 | -0.12 4.18 | 3.4 | 0.065 (ns) |
| Random effects | Intercept σ \| CI | 2.62 | 1.72 3.99 | | |
| | Residual variance \| CI | 8.03 | 7.44 8.66 | | |

*Table D3.* Model s-VR.
Results of the LMEM fit including the last, not significant term of the VR exploration condition over session and task only for the 3 second time condition. #obs means total number of experimental observations. #fix means number of fixed effect coefficients. #rand means total number of random effect coefficients. #cov means number of covariance parameters. SE is standard error and CI 95% confidence interval borders. $\chi 2(1)$ is the likelihood ratio statistic with 1 degree of freedom difference to the model without this fixed effect. σ is the random effect standard deviation. Here the random effect session σ was close to zero (0.008) which leads to numerical instabilities and means one can and should ignore this effect. We therefore did not include it in the model.
The LMEM formula is: 'CondResult~1+Session+(1|Subject)'.

| Condition | β0(Intercept) | β1(Session) | β4(Abs to Rel) | β3(Abs to Poi) |
|---|---|---|---|---|
| S1 Abs 3s | 1 | -.5 | 0 | 0 |
| S1 Rel 3s | 1 | -.5 | 1 | 0 |
| S1 Poi 3s | 1 | -.5 | 0 | 1 |
| S2 Abs 3s | 1 | 0 | 0 | 0 |
| S2 Rel 3s | 1 | 0 | 1 | 0 |
| S2 Poi 3s | 1 | 0 | 0 | 1 |
| S3 Abs 3s | 1 | .5 | 0 | 0 |
| S3 Rel 3s | 1 | .5 | 1 | 0 |
| S3 Poi 3s | 1 | .5 | 0 | 1 |

*Table D4.* Model s-VR Matrix.
The fixed effect design matrix of the LMEM fit including the last, not significant term of the VR exploration condition over session and task only for the 3 second time condition. The grey columns denote all main or simple effect predictors and their matrix coding, which dropped out in the process of model selection. Note that the maximum model before step down selection included all predictors depicted here as well as all their interactions as fixed effects. The maximum model also included all fixed effect terms except interactions as random effects.
The maximum LMEM formula is: 'CondResult~Session*AbsRel+Session*AbsPoi+(1+Session+AbsRel+AbsPoi | Subject)'.

| | | | | | |
|---|---|---|---|---|---|
| Degrees of freedom (df) | #obs \| #fix \| #rand \| #cov | 396 | 4 | 164 | 3 |
| Fixed effects | β0(Intercept) ± SE \| CI | 53.82 ± 0.68 | 52.48 55.16 | | |
| | β1(Session) ± SE \| CI \| χ2(1) \| p | 7.14 ± 1.35 | 4.49 9.79 | 27.07 | $2*10^{-7}$ |
| | β3(Abs to Poi) ± SE \| CI \| χ2(1) \| p | 2.2 ± 0.93 | 0.37 4 | 5.54 | 0.018 |
| | β1*β4(Abs to Rel) ± SE \| CI \| χ2(1) \| p | -5.6 ± 2.1 | -9.69 -1.51 | 7.18 | 0.0074 |
| Random effects | Intercept σ \| CI | 2.65 | 1.69 4.16 | | |
| | Residual variance \| CI | 8.51 | 7.89 9.18 | | |

*Table D5.* Model u-VR.
Results of the best LMEM fit of the VR exploration condition over sessions and tasks for the unlimited time condition results. #obs means total number of experimental observations. #fix means number of fixed effect coefficients. #rand means total number of random effect coefficients. #cov means number of covariance parameters. SE is standard error and CI 95% confidence interval borders. χ2(1) is the likelihood ratio statistic with 1 degree of freedom difference to the model without this fixed effect. σ is the random effect standard deviation. Here both random effects session σ and absolute to pointing task difference σ were close to zero (0.008, 0.01) which leads to numerical instabilities and means one can and should ignore these effects. We therefore did not include them in the model.
The best fit LMEM formula is: 'CondResult~1+Session+AbsPoi+AbsRel:Session+(1|Subject)'.
Note 'AbsRel:Session' means only the interaction of 'AbsRel' and 'Session', while 'AbsRel*Session' would also include both 'AbsRel' and 'Session' in addition to their interaction.

| Condition | β0(Intercept) | β1(Session) | β3(Abs to Poi) | β4(Abs to Rel) | β1(Session)* β4(Abs to Rel) |
|---|---|---|---|---|---|
| S1 Abs Unl | 1 | -.5 | 0 | 0 | 0 |
| S1 Rel Unl | 1 | -.5 | 0 | 1 | -.5 |
| S1 Poi Unl | 1 | -.5 | 1 | 0 | 0 |
| S2 Abs Unl | 1 | 0 | 0 | 0 | 0 |
| S2 Rel Unl | 1 | 0 | 0 | 1 | 0 |
| S2 Poi Unl | 1 | 0 | 1 | 0 | 0 |
| S3 Abs Unl | 1 | .5 | 0 | 0 | 0 |
| S3 Rel Unl | 1 | .5 | 0 | 1 | .5 |
| S3 Poi Unl | 1 | .5 | 1 | 0 | 0 |

*Table D6.* Model u-VR Matrix.
The fixed effect design matrix of the best fit LMEM fit of the VR exploration condition over session and task only for the unlimited time condition results. The grey columns denote all main or simple effect predictors and their matrix coding, which dropped out in the process of model selection. Note that the maximum model before step down selection included all predictors depicted here as well as all their interactions as fixed effects. The maximum model also included all fixed effect terms except interactions as random effects.
The maximum LMEM formula is: 'CondResult ~ Session*AbsRel+Session*AbsPoi+(1+Session+AbsRel+AbsPoi | Subject)'.

#### Best-fit Map Models

We provide details on all linear models used for the quantitative analysis of the map exploration condition as presented in Results. Table D7 and D8 hold model results and design matrix, respectively, of the fit over session, task and time. Table D9 and D10 depict results and design matrix of the fit over session, task and three seconds decision time. Table D11 and D12 contain results and design matrix of the fit over session, task and unlimited decision time.

| Degrees of freedom (df) | #obs \| #fix \| #rand \| #cov | 768 | 4 | 296 | 5 |
|---|---|---|---|---|---|
| Fixed effects | β0(Intercept) ± SE \| CI | 55.11 ± 0.7 | 53.73 56.48 | | |
| | β1(Session) ± SE \| CI \| χ2(1) \| p | 6.55 ± 1.16 | 4.27 8.83 | 26.87 | $2.2*10^{-7}$ |
| | β2(Time) ± SE \| CI \| χ2(1) \| p | 5.35 ± 0.84 | 3.71 7 | 30.99 | $2.6*10^{-8}$ |
| | β3(Mean to Poi) ± SE \| CI \| χ2(1) \| p | -2.15 ± 0.74 | -3.59 -0.7 | 7.08 | 0.0079 |
| Random effects | Intercept σ \| CI | 3.74 | 2.94 4.77 | | |
| | Session σ \| CI | 4.71 | 3.03 7.31 | | |
| | Time σ \| CI | 4.53 | 3.09 6.66 | | |
| | Mean to Poi σ \| CI | 2.65 | 1.15 6.13 | | |
| | Residual variance \| CI | 8.52 | 8 9.06 | | |

*Table D7.* Model a-Map.
Results of the best LMEM fit of the map exploration condition over session, task and time. #obs means total number of experimental observations. #fix means number of fixed effect coefficients. #rand means total number of random effect coefficients. #cov means number of covariance parameters. SE is standard error and CI 95% confidence interval borders. χ2(1) is the likelihood ratio statistic with 1 degree of freedom difference to the model without this fixed effect. σ is the random effect standard deviation.
The best fit LMEM formula is: 'CondResult ~ 1+Time+Session+MeanPoi+(1+Time+Session+MeanPoi | Subject)'.

| Condition | β0(Intercept) | β1(Session) | β2(Time) | β3(Mean to Poi) | β4(Mean to Abs) |
|---|---|---|---|---|---|
| S1 Abs 3s | 1 | -.5 | -.5 | 0 | 1 |
| S1 Rel 3s | 1 | -.5 | -.5 | -1 | -1 |
| S1 Poi 3s | 1 | -.5 | -.5 | 1 | 0 |
| S1 Abs Unl | 1 | -.5 | .5 | 0 | 1 |
| S1 Rel Unl | 1 | -.5 | .5 | -1 | -1 |
| S1 Poi Unl | 1 | -.5 | .5 | 1 | 0 |
| S2 Abs 3s | 1 | 0 | -.5 | 0 | 1 |
| S2 Rel 3s | 1 | 0 | -.5 | -1 | -1 |
| S2 Poi 3s | 1 | 0 | -.5 | 1 | 0 |
| S2 Abs Unl | 1 | 0 | .5 | 0 | 1 |
| S2 Rel Unl | 1 | 0 | .5 | -1 | -1 |
| S2 Poi Unl | 1 | 0 | .5 | 1 | 0 |
| S3 Abs 3s | 1 | .5 | -.5 | 0 | 1 |
| S3 Rel 3s | 1 | .5 | -.5 | -1 | -1 |
| S3 Poi 3s | 1 | .5 | -.5 | 1 | 0 |
| S3 Abs Unl | 1 | .5 | .5 | 0 | 1 |
| S3 Rel Unl | 1 | .5 | .5 | -1 | -1 |
| S3 Poi Unl | 1 | .5 | .5 | 1 | 0 |

*Table D8.* Model a-Map Matrix.
The fixed effect design matrix used to fit the map exploration condition over session, task and time. The grey columns denote all main or simple effect predictors and their matrix coding, which dropped out in the process of model selection. Note that the maximum model before step down selection included all predictors depicted here as well as all their interactions as fixed effects. The maximum model also included all fixed effect terms except interactions as random effects.
The maximum LMEM formula is:
'CondResult ~ Time*Session*MeanPoi+Time*Session*MeanAbs+(1+Time+Session+MeanPoi+MeanAbs | Subject)'.

| Degrees of freedom (df) | #obs \| #fix \| #rand \| #cov | 384 | 5 | 222 | 4 |
|---|---|---|---|---|---|
| Fixed effects | β0(Intercept) ± SE \| CI | 48.09 ± 0.85 | 46.43 49.75 | | |
| | β1(Poi to Abs) ± SE \| CI \| χ2(1) \| p | 3.86 ± 1.15 | 1.59 6.13 | 10.94 | 0.00094 |
| | β2(Poi to Rel) ± SE \| CI \| χ2(1) \| p | 5.38 ± 1.25 | 2.93 7.83 | 16.34 | 0.000053 |
| | β1*β3(Session) ± SE \| CI \| χ2(1) \| p | 6.14 ± 1.96 | 2.29 9.98 | 9.59 | 0.002 |
| | β2*β3(Session) ± SE \| CI \| χ2(1) \| p | 7.97 ± 2.01 | 4.01 11.9 | 14.88 | 0.00011 |
| Random effects | Intercept σ \| CI | 2.91 | 1.75 4.85 | | |
| | Poi to Abs | 1.42 | 0 91.29 | | |
| | Poi to Rel | 3.56 | 1.68 7.55 | | |

| | | | |
|---|---|---|---|
| | Residual variance \| CI | 8.59 | 7.86<br>9.38 |

*Table D9.* Model s-Map.
Results of the best fit LMEM of the map exploration condition over session and task only for the 3 second time condition. #obs means total number of experimental observations. #fix means number of fixed effect coefficients. #rand means total number of random effect coefficients. #cov means number of covariance parameters. SE is standard error and CI 95% confidence interval borders. $\chi2(1)$ is the likelihood ratio statistic with 1 degree of freedom difference to the model without this fixed effect. σ is the random effect standard deviation.
The best fit LMEM formula is: 'CondResult~1+PoiAbs+PoiRel+PoiAbs:Session+PoiRel:Session+(1+PoiAbs+PoiRel | Subject)'.

| Condition | β0(Intercept) | β1(Poi to Abs) | β2(Poi to Rel) | β3(Session) | β1(Poi to Abs)* β3(Session) | β2(Poi to Rel)* β3(Session) |
|---|---|---|---|---|---|---|
| S1 Abs 3s | 1 | 1 | 0 | -.5 | .-5 | 0 |
| S1 Rel 3s | 1 | 0 | 1 | -.5 | 0 | -.5 |
| S1 Poi 3s | 1 | 0 | 0 | -.5 | 0 | 0 |
| S2 Abs 3s | 1 | 1 | 0 | 0 | 0 | 0 |
| S2 Rel 3s | 1 | 0 | 1 | 0 | 0 | 0 |
| S2 Poi 3s | 1 | 0 | 0 | 0 | 0 | 0 |
| S3 Abs 3s | 1 | 1 | 0 | .5 | .5 | 0 |
| S3 Rel 3s | 1 | 0 | 1 | .5 | 0 | .5 |
| S3 Poi 3s | 1 | 0 | 0 | .5 | 0 | 0 |

*Table D10.* Model s-Map Matrix.
The fixed effect design matrix of the best fit LMEM fit of the map exploration condition over session and task only for the 3 second time condition results. The grey columns denote all main or simple effect predictors and their matrix coding, which dropped out in the process of model selection. Note that the maximum model before step down selection included all predictors depicted here as well as all their interactions as fixed effects. The maximum model also included all fixed effect terms except interactions as random effects.
The maximum LMEM formula is: 'CondResult~Session*PoiAbs+Session*PoiRel+(1+Session+PoiAbs+PoiRel | Subject)'.

| | | | | | |
|---|---|---|---|---|---|
| Degrees of freedom (df) | #obs \| #fix \| #rand \| #cov | 384 | 4 | 148 | 3 |
| Fixed effects | β0(Intercept) ± SE \| CI | 57.65<br>± 0.93 | 55.83<br>59.47 | | |
| | β1(Session) ± SE \| CI \| χ2(1) \| p | 5.21<br>± 1.6 | 2.04<br>8.38 | 10.14 | 0.0015 |
| | β2(Rel to Poi) ± SE \| CI \| χ2(1) \| p | -1.86<br>± 0.96 | -3.76<br>0 | 3.7 | 0.054 (ns) |
| | β1*β3(Rel to Abs) ± SE \| CI \| χ2(1) \| p | 4.59<br>± 2.18 | 0.3<br>8.8 | 4.38 | 0.036 |
| Random effects | Intercept σ \| CI | 5.12 | 4<br>6.52 | | |
| | Session σ \| CI | 4.3 | 1.89<br>9.8 | | |
| | Residual variance \| CI | 8.71 | 8.03<br>9.45 | | |

*Table D11.* Model u-Map.
Results of the LMEM fit including the two last, not significant terms (β1*β3 interaction is only significant because of β2) of the map exploration condition over session and task only for the unlimited time condition. #obs means total number of experimental observations. #fix means number of fixed effect coefficients. #rand means total number of random effect coefficients. #cov means number of covariance parameters. SE is standard error and CI 95% confidence interval borders. $\chi2(1)$ is the likelihood ratio statistic with 1 degree of freedom difference to the model

without this fixed effect. σ is the random effect standard deviation. Random effect Relative to Pointing task difference σ was close to zero (0.007) which leads to numerical instabilities and means one can and should ignore this effect. We therefore did not include it in the model. The best fit LMEM formula is: 'CondResult~1+Session+RelPoi+RelAbs:Session+(1+Session | Subject)'.

| Condition | β0(Intercept) | β1(Session) | β2(Rel to Poi) | β3(Rel to Abs) | β1(Session)* β3(Rel to Abs) |
|---|---|---|---|---|---|
| S1 Abs Unl | 1 | -.5 | 0 | 1 | .-5 |
| S1 Rel Unl | 1 | -.5 | 0 | 0 | 0 |
| S1 Poi Unl | 1 | -.5 | 1 | 0 | 0 |
| S2 Abs Unl | 1 | 0 | 0 | 1 | 0 |
| S2 Rel Unl | 1 | 0 | 0 | 0 | 0 |
| S2 Poi Unl | 1 | 0 | 1 | 0 | 0 |
| S3 Abs Unl | 1 | .5 | 0 | 1 | .5 |
| S3 Rel Unl | 1 | .5 | 0 | 0 | 0 |
| S3 Poi Unl | 1 | .5 | 1 | 0 | 0 |

*Table D12.* Model u-Map Matrix.
The fixed effect design matrix of the LMEM fit including the two last, not significant terms of the map exploration condition over session and task, only for the unlimited time condition results. The grey columns denote all main or simple effect predictors and their matrix coding, which dropped out in the process of model selection. Note that the maximum model before step down selection included all predictors depicted here as well as all their interactions as fixed effects. The maximum model also included all fixed effect terms except interactions as random effects. The maximum LMEM formula is: 'CondResult~Session*RelPoi+Session*RelAbs+(1+Session+RelPoi+RelAbs | Subject)'.

#### Best-fit VR with Belt Models

We provide details on all linear models used for the quantitative analysis of the VR with belt exploration condition as presented in Results. Table D13 and D14 hold model results and design matrix, respectively, of the fit over session, task and time. Table D15 and D16 depict results and design matrix of the fit over session, task and three seconds decision time. Table D17 and D18 contain results and design matrix of the fit over session, task and unlimited decision time.

| | | | | | |
|---|---|---|---|---|---|
| Degrees of freedom (df) | #obs \| #fix \| #rand \| #cov | 714 | 4 | 210 | 4 |
| Fixed effects | β0(Intercept) ± SE \| CI | 51.43 ± 0.45 | 50.54 52.32 | | |
| | β1(Session) ± SE \| CI \| χ2(1) \| p | 3.4 ± 0.81 | 1.8 4.99 | 17.23 | 0.000055 |
| | β2(Time) ± SE \| CI \| χ2(1) \| p | 3.3 ± 0.89 | 1.55 5.05 | 12.43 | 0.00042 |
| | β3(Mean to Poi) ± SE \| CI \| χ2(1) \| p | 1.11 ± 0.41 | 0.31 1.91 | 7 | 0.0082 |
| Random effects | Intercept σ \| CI | 2.17 | 1.43 3.27 | | |
| | Time σ \| CI | 5.11 | 3.48 7.5 | | |
| | Mean to Poi σ \| CI | 1.31 | 0.47 3.64 | | |
| | Residual variance \| CI | 8.07 | 7.59 8.57 | | |

*Table D13.* Model a-VRwB.
Results of the best LMEM fit of the VR with belt exploration condition over session, task and time. #obs means total number of experimental observations. #fix means number of fixed effect coefficients. #rand means total number of random effect coefficients. #cov means number of covariance parameters. SE is standard error and CI 95% confidence interval borders. χ2(1) is the likelihood ratio statistic with 1 degree of

freedom difference to the model without this fixed effect. σ is the random effect standard deviation. Random effect session σ was close to zero (0.009) which leads to numerical instabilities and means one can and should ignore this effect. We therefore did not include it in the model. The best fit LMEM formula is: 'CondResult~1+Time+Session+MeanPoi+(1+Time+MeanPoi | Subject)'.

| Condition | β0(Intercept) | β1(Session) | β2(Time) | β3(Mean to Poi) | β4(Mean to Rel) |
|---|---|---|---|---|---|
| S1 Abs 3s | 1 | -.5 | -.5 | -1 | -1 |
| S1 Rel 3s | 1 | -.5 | -.5 | 0 | 1 |
| S1 Poi 3s | 1 | -.5 | -.5 | 1 | 0 |
| S1 Abs Unl | 1 | -.5 | .5 | -1 | -1 |
| S1 Rel Unl | 1 | -.5 | .5 | 0 | 1 |
| S1 Poi Unl | 1 | -.5 | .5 | 1 | 0 |
| S2 Abs 3s | 1 | 0 | -.5 | -1 | -1 |
| S2 Rel 3s | 1 | 0 | -.5 | 0 | 1 |
| S2 Poi 3s | 1 | 0 | -.5 | 1 | 0 |
| S2 Abs Unl | 1 | 0 | .5 | -1 | -1 |
| S2 Rel Unl | 1 | 0 | .5 | 0 | 1 |
| S2 Poi Unl | 1 | 0 | .5 | 1 | 0 |
| S3 Abs 3s | 1 | .5 | -.5 | -1 | -1 |
| S3 Rel 3s | 1 | .5 | -.5 | 0 | 1 |
| S3 Poi 3s | 1 | .5 | -.5 | 1 | 0 |
| S3 Abs Unl | 1 | .5 | .5 | -1 | -1 |
| S3 Rel Unl | 1 | .5 | .5 | 0 | 1 |
| S3 Poi Unl | 1 | .5 | .5 | 1 | 0 |

*Table D14.* Model a-VRwB Matrix.
The fixed effect design matrix used to fit the map exploration condition over session, task and time. The grey columns denote all main or simple effect predictors and their matrix coding, which dropped out in the process of model selection. Note that the maximum model before step down selection included all predictors depicted here as well as all their interactions as fixed effects. The maximum model also included all fixed effect terms except interactions as random effects.
The maximum LMEM formula is:
'CondResult ~ Time*Session*MeanRel+Time*Session*MeanRel+(1+Time+Session+MeanRel+MeanPoi | Subject)'.

| | | | | | |
|---|---|---|---|---|---|
| Degrees of freedom (df) | #obs \| #fix \| #rand \| #cov | 357 | 3 | 70 | 2 |
| Fixed effects | β0(Intercept) ± SE \| CI | 49.72 ± 0.62 | 48.51 50.93 | | |
| | β1(Poi to Abs) ± SE \| CI \| χ2(1) \| p *β3(Session) | 5.24 ± 1.89 | 1.53 8.96 | 7.63 | 0.0057 |
| | β2(Poi to Rel) ± SE \| CI \| χ2(1) \| p *β3(Session) | 3.88 ± 1.89 | 0.17 7.6 | 4.21 | 0.04 |
| Random effects | Intercept σ \| CI | 2.7 | 1.55 4.7 | | |
| | Residual variance \| CI | 8.68 | 8 9.42 | | |

**Table D15.** Model s-VRwB.
Results of the best LMEM fit of the VR with belt exploration condition over sessions and tasks for the 3 second time condition. #obs means total number of experimental observations. #fix means number of fixed effect coefficients. #rand means total number of random effect coefficients. #cov means number of covariance parameters. SE is standard error and CI 95% confidence interval borders. χ2(1) is the likelihood ratio statistic with 1 degree of freedom difference to the model without this fixed effect. σ is the random effect standard deviation. Here the

random effect session σ was close to zero (0.02) which leads to numerical instabilities and means one can and should ignore this effect. We therefore did not include it in the model.
The best fit LMEM formula is: ‘Condition~1+PoiAbs:Session+PoiRel:Session+(1|Subject)’.
Different from ‘PoiAbs*Session’, ‘PoiAbs:Session’ only includes the interaction in the model.

| Condition | β0(Intercept) | β1(Poi to Abs) | β2(Poi to Rel) | β3(Session) | β1(Poi to Abs)* β3(Session) | β2(Poi to Rel)* β3(Session) |
|---|---|---|---|---|---|---|
| S1 Abs 3s | 1 | 1 | 0 | -.5 | .-5 | 0 |
| S1 Rel 3s | 1 | 0 | 1 | -.5 | 0 | -.5 |
| S1 Poi 3s | 1 | 0 | 0 | -.5 | 0 | 0 |
| S2 Abs 3s | 1 | 1 | 0 | 0 | 0 | 0 |
| S2 Rel 3s | 1 | 0 | 1 | 0 | 0 | 0 |
| S2 Poi 3s | 1 | 0 | 0 | 0 | 0 | 0 |
| S3 Abs 3s | 1 | 1 | 0 | .5 | .5 | 0 |
| S3 Rel 3s | 1 | 0 | 1 | .5 | 0 | .5 |
| S3 Poi 3s | 1 | 0 | 0 | .5 | 0 | 0 |

*Table D16.* Model s-VRwB Matrix.
The fixed effect design matrix of the best fit LMEM fit of the VRwB exploration condition over session and task only for the 3 second time condition results. The grey columns denote all main or simple effect predictors and their matrix coding, which dropped out in the process of model selection. Note that the maximum model before step down selection included all predictors depicted here as well as all their interactions as fixed effects. The maximum model also included all fixed effect terms except interactions as random effects.
The maximum LMEM formula is: ‘CondResult~Session*PoiAbs+Session*PoiRel+(1+Session+PoiAbs+PoiRel | Subject)’.

| | | | | | |
|---|---|---|---|---|---|
| Degrees of freedom (df) | #obs \| #fix \| #rand \| #cov | 357 | 3 | 70 | 2 |
| Fixed effects | β0(Intercept) ± SE \| CI | 52.6 ± 0.75 | 51.13 54.07 | | |
| | β1(Session) ± SE \| CI \| χ2(1) \| p | 3.89 ± 1.11 | 1.7 6.08 | 11.96 | 0.00054 |
| | β2(Abs to Rel) ± SE \| CI \| χ2(1) \| p | 1.83 ± 0.85 | 0.16 3.51 | 4.6 | 0.032 |
| Random effects | Intercept σ \| CI | 3.84 | 2.74 5.37 | | |
| | Residual variance \| CI | 7.58 | 6.98 8.23 | | |

*Table D17.* Model u-VRwB.
Results of the best LMEM fit of the VR with belt exploration condition over sessions and tasks only for the unlimited time condition. #obs means total number of experimental observations. #fix means number of fixed effect coefficients. #rand means total number of random effect coefficients. #cov means number of covariance parameters. SE is standard error and CI 95% confidence interval borders. χ2(1) is the likelihood ratio statistic with 1 degree of freedom difference to the model without this fixed effect. σ is the random effect standard deviation. Here both the random effects session σ and difference from Absolute to Relative task σ were close to zero (0.01, 0.02) which leads to numerical instabilities and means one can and should ignore these effects. We therefore did not include them in the model.
The best fit LMEM formula is: ‘Condition~1+Session+AbsRel+(1|Subject)’.

| Condition | β0(Intercept) | β1(Session) | β2(Abs to Rel) | β3(Abs to Poi) |
|---|---|---|---|---|
| S1 Abs Unl | 1 | -.5 | 1 | 0 |
| S1 Rel Unl | 1 | -.5 | 0 | 0 |
| S1 Poi Unl | 1 | -.5 | 0 | 1 |
| S2 Abs Unl | 1 | 0 | 1 | 0 |
| S2 Rel Unl | 1 | 0 | 0 | 0 |
| S2 Poi Unl | 1 | 0 | 0 | 1 |
| S3 Abs Unl | 1 | .5 | 1 | 0 |
| S3 Rel Unl | 1 | .5 | 0 | 0 |
| S3 Poi Unl | 1 | .5 | 0 | 1 |

*Table D18.* Model u-VRwB Matrix.
The fixed effect design matrix of the best fit LMEM fit of the VR with belt exploration condition over session and task only for the unlimited time condition results. The grey columns denote all main or simple effect predictors and their matrix coding, which dropped out in the process of model selection. Note that the maximum model before step down selection included all predictors depicted here as well as all their interactions as fixed effects. The maximum model also included all fixed effect terms except interactions as random effects.
The maximum LMEM formula is: 'CondResult ~ Session*AbsRel+Session*AbsPoi+(1+Session+AbsRel+AbsPoi | Subject)'.

#### Best-fit VR-as-Reference Models

##### Best-fit VR Versus Map Models

We provide details on all linear models used for the quantitative analysis of the comparison of VR and map exploration condition as presented in Results. Table D19 and D20 hold model results and design matrix, respectively, of the fit over experimental exploration, session, task and time. Table D21 and D22 show the results and design matrix, fitted on exploration, session and time for only the Pointing task.

Table D23 and D24 depict results and design matrix of the fit over exploration, session, task and three seconds decision time. Table D25 and D26 contain results and design matrix of the fit over exploration, session, task and unlimited decision time.

| Degrees of freedom (df) | #obs \| #fix \| #rand \| #cov | 1560 | 10 | 468 | 4 |
|---|---|---|---|---|---|
| Fixed effects | β0(Intercept) ± SE \| CI | 53.15 ± 0.4 | 52.4 53.9 | | |
| | β1(Exploration) ± SE \| CI \| χ2(1) \| p | 2.29 ± 0.79 | 0.73 3.85 | 8.16 | 0.0043 |
| | β2(Session) ± SE \| CI \| χ2(1) \| p | 4.96 ± 0.7 | 3.58 6.33 | 43.48 | $4.3*10^{-11}$ |
| | β3(Time) ± SE \| CI \| χ2(1) \| p | 4.82 ± 0.52 | 3.81 5.84 | 65.9 | $4.4*10^{-16}$ |
| | β1*β4(Mean to Rel) ± SE \| CI \| χ2(1) \| p | 2.13 ± 0.68 | 0.8 3.45 | 9.82 | 0.002 |
| | β1*β5(Mean to Poi) ± SE \| CI \| χ2(1) \| p | -3.07 ± 0.68 | -4.4 -1.75 | 20.44 | $6.2*10^{-6}$ |
| | β1*β2 ± SE \| CI \| χ2(1) \| p | 2.79 ± 1.4 | 0.04 5.54 | 3.9 | 0.048 |
| | β1*β2*β4 ± SE \| CI \| χ2(1) \| p | 3.26 ± 1.52 | 0.28 6.25 | 4.6 | 0.032 |
| | β1*β2*β5 ± SE \| CI \| χ2(1) \| p | -4.39 ± 1.52 | -7.37 -1.41 | 8.31 | 0.004 |

| | | | | | |
|---|---|---|---|---|---|
| | β2*β3*β4 ± SE \| CI \| χ2(1) \| p | -2.8<br>± 1.18 | -5.12<br>-0.49 | 5.61 | 0.018 |
| Random effects | Intercept σ \| CI | 3.15 | 2.62<br>3.8 | | |
| | Session σ \| CI | 3.16 | 1.89<br>5.3 | | |
| | Time σ \| CI | 3.25 | 2.2<br>4.8 | | |
| | Residual variance \| CI | 8.48 | 8.16<br>8.83 | | |

*Table D19.* Model a-VRvMap.
Results of the best LMEM fit of both, VR and map exploration condition, over session, task and time . #obs means total number of experimental observations. #fix means number of fixed effect coefficients. #rand means total number of random effect coefficients. #cov means number of covariance parameters. SE is standard error and CI 95% confidence interval borders. χ2(1) is the likelihood ratio statistic with 1 degree of freedom difference to the model without this fixed effect. σ is the random effect standard deviation.
The best fit LMEM formula is:
'Condition~1+ Exploration +Session+Time+ Exploration:MeanRel+ Exploration:MeanPoi+Time:Session+ Exploration:Session:MeanRel+ Exploration:Session:MeanPoi+Session:Time:MeanRel+(1+Session+Time|Subject)'.
Different from 'Exploration *MeanRel', 'Exploration:MeanRel' only includes the interaction in the model.

| Condition | β0 (Int.) | β1 (Expl.) | β2 (Sess.) | β3 (Time) | β4(Mean to Rel) | β5(Mean to Poi) | β1* β4 | β1* β5 | β1* β2 | β1* β2* β4 | β1* β2* β5 | β2* β3* β4 |
|---|---|---|---|---|---|---|---|---|---|---|---|---|
| S1 Abs 3s VR | 1 | -.5 | -.5 | -.5 | -1 | -1 | .5 | .5 | .25 | -.25 | -.25 | -.25 |
| S1 Rel 3s VR | 1 | -.5 | -.5 | -.5 | 1 | 0 | -.5 | 0 | .25 | .25 | 0 | .25 |
| S1 Poi 3s VR | 1 | -.5 | -.5 | -.5 | 0 | 1 | 0 | -.5 | .25 | 0 | .25 | 0 |
| S1 Abs Unl VR | 1 | -.5 | -.5 | .5 | -1 | -1 | .5 | .5 | .25 | -.25 | -.25 | .25 |
| S1 Rel Unl VR | 1 | -.5 | -.5 | .5 | 1 | 0 | -.5 | 0 | .25 | .25 | 0 | -.25 |
| S1 Poi Unl VR | 1 | -.5 | -.5 | .5 | 0 | 1 | 0 | -.5 | .25 | 0 | .25 | 0 |
| S2 Abs 3s VR | 1 | -.5 | 0 | -.5 | -1 | -1 | .5 | .5 | 0 | 0 | 0 | 0 |
| S2 Rel 3s VR | 1 | -.5 | 0 | -.5 | 1 | 0 | -.5 | 0 | 0 | 0 | 0 | 0 |
| S2 Poi 3s VR | 1 | -.5 | 0 | -.5 | 0 | 1 | 0 | -.5 | 0 | 0 | 0 | 0 |
| S2 Abs Unl VR | 1 | -.5 | 0 | .5 | -1 | -1 | .5 | .5 | 0 | 0 | 0 | 0 |
| S2 Rel Unl VR | 1 | -.5 | 0 | .5 | 1 | 0 | -.5 | 0 | 0 | 0 | 0 | 0 |
| S2 Poi Unl VR | 1 | -.5 | 0 | .5 | 0 | 1 | 0 | -.5 | 0 | 0 | 0 | 0 |
| S3 Abs 3s VR | 1 | -.5 | .5 | -.5 | -1 | -1 | .5 | .5 | -.25 | .25 | .25 | .25 |
| S3 Rel 3s VR | 1 | -.5 | .5 | -.5 | 1 | 0 | -.5 | 0 | -.25 | -.25 | 0 | -.25 |
| S3 Poi 3s VR | 1 | -.5 | .5 | -.5 | 0 | 1 | 0 | -.5 | -.25 | 0 | -.25 | 0 |
| S3 Abs Unl VR | 1 | -.5 | .5 | .5 | -1 | -1 | .5 | .5 | -.25 | .25 | .25 | -.25 |
| S3 Rel Unl VR | 1 | -.5 | .5 | .5 | 1 | 0 | -.5 | 0 | -.25 | -.25 | 0 | .25 |
| S3 Poi Unl | 1 | -.5 | .5 | .5 | 0 | 1 | 0 | -.5 | -.25 | 0 | -.25 | 0 |

| | | | | | | | | | | | | |
|---|---|---|---|---|---|---|---|---|---|---|---|---|
| VR | | | | | | | | | | | | |
| S1 Abs 3s Map | 1 | .5 | -.5 | -.5 | -1 | -1 | -.5 | -.5 | -.25 | .25 | .25 | -.25 |
| ... | | | | | | | | | | | | |

*Table D20.* Model a-VRvMap Matrix.
The fixed effect design matrix used to fit both, VR and map exploration condition, over session, task and time. The grey columns denote all main or simple effect predictors and their matrix coding, which dropped out in the process of model selection. Note that the maximum model before step down selection included all predictors depicted here as well as all their interactions as fixed effects. The maximum model also included all fixed effect terms except interactions as random effects.
The maximum LMEM formula is:
'CondResult ~ Expl*Time*Session*MeanRel+Expl*Time*Session*MeanPoi+(1+Time+Session+MeanRel+MeanPoi | Subject)'.

| | | | | | |
|---|---|---|---|---|---|
| Degrees of freedom (df) | #obs \| #fix \| #rand \| #cov | 520 | 5 | 468 | 4 |
| Fixed effects | β0(Intercept) ± SE \| CI | 52.6 ± 0.44 | 51.73 53.47 | | |
| | β1(Session) ± SE \| CI \| χ2(1) \| p | 4.31 ± 0.96 | 2.42 6.2 | 18.17 | 0.00005 |
| | β2(Time) ± SE \| CI \| χ2(1) \| p | 5.9 ± 0.85 | 4.24 7.57 | 39.42 | $3.4*10^{-10}$ |
| | β3(Explore) ± SE \| CI \| χ2(1) \| p | -0.85 ± 0.88 | -2.58 0.88 | 0.87 | 0.35 (ns) |
| | β1*β3 ± SE \| CI \| χ2(1) \| p | -1.83 ± 1.93 | -5.61 1.96 | 0.88 | 0.35 (ns) |
| Random effects | Intercept σ \| CI | 1.85 | 0.85 4.03 | | |
| | Session σ \| CI | 3.53 | 1.16 10.76 | | |
| | time σ \| CI | 5.42 | 3.24 9.08 | | |
| | Residual variance \| CI | 7.96 | 7.14 8.87 | | |

*Table D21.* Model p-VRvMap.
Results of the LMEM fit including the two last, not significant terms (visual inspection allows both β3 and β1*β3 as effects), of the VR and map exploration condition over session and time only for the pointing task. #obs means total number of experimental observations. #fix means number of fixed effect coefficients. #rand means total number of random effect coefficients. #cov means number of covariance parameters. SE is standard error and CI 95% confidence interval borders. χ2(1) is the likelihood ratio statistic with 1 degree of freedom difference to the model without this fixed effect. σ is the random effect standard deviation.
The fitted LMEM formula is: 'CondResult~1+Session+Time+ Exploration + Exploration:Session+(1+Session+Time | Subject)'.

| Condition | β0(Intercept) | β1(Session) | β2(Time) | β3(Exploration) | β1(Session)* β3(Exploration) |
|---|---|---|---|---|---|
| S1 Poi 3s VR | 1 | -.5 | -.5 | -.5 | .25 |
| S1 Poi Unl VR | 1 | -.5 | .5 | -.5 | .25 |
| S2 Poi 3s VR | 1 | 0 | -.5 | -.5 | 0 |
| S2 Poi Unl VR | 1 | 0 | .5 | -.5 | 0 |
| S3 Poi 3s VR | 1 | .5 | -.5 | -.5 | -.25 |

| | | | | | |
|---|---|---|---|---|---|
| S3 Poi Unl VR | 1 | .5 | .5 | -.5 | -.25 |
| S1 Poi 3s Map | 1 | -.5 | .-5 | .5 | -.25 |
| ... | | | | | |

*Table D22.* Model p-VRvMap Matrix.
The fixed effect design matrix of the LMEM fit including the two last, not significant terms, of the VR and map exploration condition over session and time only for the pointing task results. The grey columns denote all main or simple effect predictors and their matrix coding, which dropped out in the process of model selection. Note that the maximum model before step down selection included all predictors depicted here as well as all their interactions as fixed effects. The maximum model also included all fixed effect terms except interactions as random effects. The maximum LMEM formula is: 'CondResult~Session*Time* Exploration +(1+Session+Time | Subject)'.

| Degrees of freedom (df) | #obs \| #fix \| #rand \| #cov | 780 | 7 | 468 | 4 |
|---|---|---|---|---|---|
| Fixed effects | β0(Intercept) ± SE \| CI | 51.02 ± 0.49 | 50.05 51.98 | | |
| | β1(Session) ± SE \| CI \| χ2(1) \| p | 3.92 ± 0.85 | 2.25 5.6 | 19.47 | 0.00001 |
| | β2(Explore) ± SE \| CI \| χ2(1) \| p | 3.51 ± 1.01 | 1.54 5.49 | 12 | 0.0005 |
| | β3(AbsPoi) ± SE \| CI \| χ2(1) \| p | -1.52 ± 0.64 | -2.79 -0.26 | 5.36 | 0.021 |
| | β1*β2 ± SE \| CI \| χ2(1) \| p | 5.87 ± 2 | 1.93 9.8 | 8.26 | 0.004 |
| | β2*β3 ± SE \| CI \| χ2(1) \| p | -4.81 ± 1.44 | -7.63 -1.99 | 10.11 | 0.001 |
| | β1*β2* β3 ± SE \| CI \| χ2(1) \| p | -6.35 ± 3.18 | -12.59 -0.11 | 3.91 | 0.048 |
| Random effects | Intercept σ \| CI | 2.89 | 2.14 3.9 | | |
| | Session σ \| CI | 2.64 | 0.73 9.63 | | |
| | Abs to Poi σ \| CI | 1.39 | 0.06 34.36 | | |
| | Residual variance \| CI | 8.31 | 7.78 8.87 | | |

*Table D23***.** Model s-VRvMap.
Results of the best LMEM fit of the VR and the map over sessions and tasks for the 3 second time condition. #obs means total number of experimental observations. #fix means number of fixed effect coefficients. #rand means total number of random effect coefficients. #cov means number of covariance parameters. SE is standard error and CI 95% confidence interval borders. χ2(1) is the likelihood ratio statistic with 1 degree of freedom difference to the model without this fixed effect. σ is the random effect standard deviation.
The best fit LMEM formula is:
'Condition~1+Session+ Exploration +AbsPoi+ Session: Exploration + Exploration:AbsPoi+Session:Exploration:AbsPoi+ (1+Session|Subject)'.

| Condition | β0 (Inter.) | β1 (Session) | β2 (Expl.) | β3(Abs to Poi) | β4(Abs to Rel) | β2*β3 | β1*β2 | β1*β2*β3 |
|---|---|---|---|---|---|---|---|---|
| S1 Abs Unl VR | 1 | -.5 | -.5 | 0 | 0 | 0 | ... | ... |
| S1 Rel Unl VR | 1 | -.5 | -.5 | 0 | 1 | ... | | |

| | | | | | |
|---|---|---|---|---|---|
| S1 Poi Unl VR | 1 | -.5 | -.5 | 1 | 0 |
| S2 Abs Unl VR | 1 | 0 | -.5 | 0 | 0 |
| S2 Rel Unl VR | 1 | 0 | -.5 | 0 | 1 |
| S2 Poi Unl VR | 1 | 0 | -.5 | 1 | 0 |
| S3 Abs Unl VR | 1 | .5 | -.5 | 0 | 0 |
| S3 Rel Unl VR | 1 | .5 | -.5 | 0 | 1 |
| S3 Poi Unl VR | 1 | .5 | -.5 | 1 | 0 |
| S1 Abs Unl Map | 1 | -.5 | .5 | 0 | 0 |
| ... | | | | | |

*Table D24.* Model s-VRvMap Matrix.
The fixed effect design matrix used to fit both, VR and map exploration condition, over session, task and the 3 second time condition results. The grey columns denote all main or simple effect predictors and their matrix coding, which dropped out in the process of model selection. Note that the maximum model before step down selection included all predictors depicted here as well as all their interactions as fixed effects. The maximum model also included all fixed effect terms except interactions as random effects.
The maximum LMEM formula is:
'CondResult ~ Expl*Session*AbsPoi+Expl*Session*AbsRel+(1+Session+AbsRel+AbsPoi | Subject)'.

| Degrees of freedom (df) | #obs \| #fix \| #rand \| #cov | 780 | 5 | 312 | 3 |
|---|---|---|---|---|---|
| Fixed effects | β0(Intercept) ± SE \| CI | 55.78 ± 0.53 | 54.74 56.81 | | |
| | β1(Session) ± SE \| CI \| χ2(1) \| p | 7.01 ± 1.04 | 4.97 9.04 | 42.28 | $7.89*10^{-11}$ |
| | β2(Exploration) ± SE \| CI \| χ2(1) \| p | 3.16 ± 1.04 | 1.12 5.21 | 9.02 | 0.003 |
| | β2*β3(Abs to Poi) ± SE \| CI \| χ2(1) \| p | -3.1 ± 1.32 | -5.69 -0.51 | 5.5 | 0.019 |
| | β1*β4(Abs to Rel) ± SE \| CI \| χ2(1) \| p | -3.15 ± 1.48 | -6.06 -0.24 | 4.49 | 0.034 |
| Random effects | Intercept σ \| CI | 4.08 | 3.34 5 | | |
| | Session σ \| CI | 2.34 | 0.46 11.9 | | |
| | Residual variance \| CI | 8.69 | 8.2 9.2 | | |

*Table D25.* Model u-VRvMap.
Results of the best LMEM fit of the VR and the map over sessions and tasks for the unlimited time condition. #obs means total number of experimental observations. #fix means number of fixed effect coefficients. #rand means total number of random effect coefficients. #cov means number of covariance parameters. SE is standard error and CI 95% confidence interval borders. χ2(1) is the likelihood ratio statistic with 1 degree of freedom difference to the model without this fixed effect. σ is the random effect standard deviation.
The best fit LMEM formula is:
'Condition~1+Session+ Exploration +Session:AbsRel+ Exploration:AbsPoi+(1+Session|Subject)'.

| Condition | β0 (Interc.) | β1 (Session) | β2 (Expl.) | β3(Abs to Poi) | β4(Abs to Rel) | β1*β4 | β2*β3 |
|---|---|---|---|---|---|---|---|
| S1 Abs Unl VR | 1 | -.5 | -.5 | 0 | 0 | 0 | … |
| S1 Rel Unl VR | 1 | -.5 | -.5 | 0 | 1 | … | |
| S1 Poi Unl VR | 1 | -.5 | -.5 | 1 | 0 | | |
| S2 Abs Unl VR | 1 | 0 | -.5 | 0 | 0 | | |
| S2 Rel Unl VR | 1 | 0 | -.5 | 0 | 1 | | |
| S2 Poi Unl VR | 1 | 0 | -.5 | 1 | 0 | | |
| S3 Abs Unl VR | 1 | .5 | -.5 | 0 | 0 | | |
| S3 Rel Unl VR | 1 | .5 | -.5 | 0 | 1 | | |
| S3 Poi Unl VR | 1 | .5 | -.5 | 1 | 0 | | |
| S1 Abs Unl Map | 1 | -.5 | .5 | 0 | 0 | | |
| ... | | | | | | | |

*Table D26.* Model u-VRvMap Matrix.
The fixed effect design matrix used to fit both, VR and map exploration condition, over session, task and the unlimited time condition results. The grey columns denote all main or simple effect predictors and their matrix coding, which dropped out in the process of model selection. Note that the maximum model before step down selection included all predictors depicted here as well as all their interactions as fixed effects. The maximum model also included all fixed effect terms except interactions as random effects.
The maximum LMEM formula is:
'CondResult ~ Expl*Session*AbsRel+Expl*Session*AbsPoi+(1+Session+AbsRel+AbsPoi| Subject)'.

**Best-fit VR Versus VR with Belt Models**

We provide details on all linear models used for the quantitative analysis of the comparison of VR and VR with belt exploration condition as presented in Results. Table D27 and D28 hold model results and design matrix, respectively, of the fit over experimental exploration, session, task and time. Table D29 and D30 depict results and design matrix of the fit over exploration, session, task and three seconds decision time. Table D31 and D32 contain results and design matrix of the fit over exploration, session, task and unlimited decision time.

| Degrees of freedom (df) | #obs \| #fix \| #rand \| #cov | 1506 | 7 | 760 | 5 |
|---|---|---|---|---|---|
| Fixed effects | β0(Intercept) ± SE \| CI | 51.74 ± 0.33 | 51.1 52.4 | | |
| | β1(Session) ± SE \| CI \| χ2(1) \| p | 3.48 ± 0.57 | 2.36 4.6 | 36.39 | $1.6*10^{-9}$ |
| | β2(Time) ± SE \| CI \| χ2(1) \| p | 3.76 ± 0.52 | 2.75 4.78 | 44.15 | $3*10^{-11}$ |
| | β3(Mean to Abs) ± SE \| CI \| χ2(1) \| p | -0.89 ± 0.34 | -1.55 -0.22 | 6.84 | 0.0089 |
| | β4(Mean to Poi) ± SE \| CI \| χ2(1) \| p | 0.75 ± 0.31 | 0.14 1.35 | 5.59 | 0.018 |
| | β1*β2*β3 ± SE \| CI \| χ2(1) \| p | 2.64 ± 1.13 | 0.41 4.86 | 5.4 | 0.02 |
| | β2*β4*β5(Exploration) ± SE \| CI \| χ2(1) \| p | -2.12 ± 1.01 | -4.11 -0.14 | 4.39 | 0.036 |

| | | | |
|---|---|---|---|
| Random effects | Intercept σ \| CI | 2.45 | 1.94<br>3.09 |
| | Time σ \| CI | 3.49 | 2.41<br>5.07 |
| | Mean to Abs σ \| CI | 1.89 | 1.28<br>2.8 |
| | Mean to Poi σ \| CI | 1.1 | 0.37<br>3.29 |
| | Residual variance \| CI | 8.01 | 7.67<br>8.37 |

*Table D27.* Model a-VRvB.
Results of the best LMEM fit of both, VR and VR with belt exploration condition, over session, task and time. #obs means total number of experimental observations. #fix means number of fixed effect coefficients. #rand means total number of random effect coefficients. #cov means number of covariance parameters. SE is standard error and CI 95% confidence interval borders. $\chi2(1)$ is the likelihood ratio statistic with 1 degree of freedom difference to the model without this fixed effect. σ is the random effect standard deviation. Random effect session σ was close to zero (0.02) which leads to numerical instabilities and means one can and should ignore this effect. We therefore did not include it in the model.
The best fit LMEM formula is:
'Condition~1+Session+Time+MeanAbs+MeanPoi+
Session:Time:MeanAbs+Time:MeanPoi:Exploration + (1+Time+MeanAbs+MeanPoi|Subject)'.
Different from 'Session*Time*MeanAbs', 'Session:Exploration:MeanAbs' only includes the interaction.

| Condition | β0 (Inter.) | β1 (Session) | β2 (Time) | β3(Mean to Abs) | β4(Mean to Poi) | β5(Expl.) | β1*β2* β3 | β2*β4* β5 | β2*β4* β5 |
|---|---|---|---|---|---|---|---|---|---|
| S1 Abs 3s | 1 | -.5 | -.5 | 1 | 0 | -.5 | .25 | ... | ... |
| S1 Rel 3s | 1 | -.5 | -.5 | -1 | -1 | -.5 | ... | | |
| S1 Poi 3s | 1 | -.5 | -.5 | 0 | 1 | -.5 | | | |
| S1 Abs Unl | 1 | -.5 | .5 | 1 | 0 | -.5 | | | |
| S1 Rel Unl | 1 | -.5 | .5 | -1 | -1 | -.5 | | | |
| S1 Poi Unl | 1 | -.5 | .5 | 0 | 1 | -.5 | | | |
| S2 Abs 3s | 1 | 0 | -.5 | 1 | 0 | -.5 | | | |
| S2 Rel 3s | 1 | 0 | -.5 | -1 | -1 | -.5 | | | |
| S2 Poi 3s | 1 | 0 | -.5 | 0 | 1 | -.5 | | | |
| S2 Abs Unl | 1 | 0 | .5 | 1 | 0 | -.5 | | | |
| S2 Rel Unl | 1 | 0 | .5 | -1 | -1 | -.5 | | | |
| S2 Poi Unl | 1 | 0 | .5 | 0 | 1 | -.5 | | | |
| S3 Abs 3s | 1 | .5 | -.5 | 1 | 0 | -.5 | | | |
| S3 Rel 3s | 1 | .5 | -.5 | -1 | -1 | -.5 | | | |
| S3 Poi 3s | 1 | .5 | -.5 | 0 | 1 | -.5 | | | |
| S3 Abs Unl | 1 | .5 | .5 | 1 | 0 | -.5 | | | |
| S3 Rel Unl | 1 | .5 | .5 | -1 | -1 | -.5 | | | |
| S3 Poi Unl | 1 | .5 | .5 | 0 | 1 | -.5 | | | |
| S1 Abs 3s | 1 | -.5 | -.5 | 1 | -0 | .5 | | | |
| ... | | | | | | | | | |

*Table D28.* Model a-VRvB Matrix.
The fixed effect design matrix used to fit both, VR and VR with belt exploration condition, over session, task and time. The grey columns denote all main or simple effect predictors and their matrix coding, which dropped out in the process of model selection. Note that the maximum

model before step down selection included all predictors depicted here as well as all their interactions as fixed effects. The maximum model also included all fixed effect terms except interactions as random effects.
The maximum LMEM formula is:
'CondResult ~ Expl*Time*Session*MeanAbs+Expl*Time*Session*MeanPoi+(1+Time+Session+MeanAbs+MeanPoi | Subject)'.

| | | | | | |
|---|---|---|---|---|---|
| Degrees of freedom (df) | #obs \| #fix \| #rand \| #cov | 753 | 3 | 304 | 3 |
| Fixed effects | β0(Intercept) ± SE \| CI | 49.15 ± 0.48 | 48.2 50.1 | | |
| | β1(Session) ± SE \| CI \| χ2(1) \| p | 2.51 ± 0.81 | 0.92 4.1 | 9.54 | 0.002 |
| | β2(Abs to Poi) ± SE \| CI \| χ2(1) \| p | 1.39 ± 0.67 | 0.07 2.7 | 4.14 | 0.042 |
| Random effects | Intercept σ \| CI | 2.62 | 1.8 3.7 | | |
| | Abs to Poi | 2.18 | 0.6 8.15 | | |
| | Residual variance \| CI | 8.28 | 7.78 8.78 | | |

*Table D29.* Model s-VRvB.
Results of the best LMEM fit of the VR and the VR with belt exploration condition over sessions and tasks for the 3 second time condition. #obs means total number of experimental observations. #fix means number of fixed effect coefficients. #rand means total number of random effect coefficients. #cov means number of covariance parameters. SE is standard error and CI 95% confidence interval borders. χ2(1) is the likelihood ratio statistic with 1 degree of freedom difference to the model without this fixed effect. σ is the random effect standard deviation. Here the random effect session σ was close to zero (0.01) which leads to numerical instabilities and means one can and should ignore this effect. We therefore did not include it in the model.
The best fit LMEM formula is: 'Condition~1+Session+AbsPoi+(1+AbsPoi|Subject)'.

| Condition | β0 (Intercept) | β1 (Session) | β2(Abs to Poi) | β3(Abs to Rel) | β4 (Explor.) |
|---|---|---|---|---|---|
| S1 Abs Unl VR | 1 | -.5 | 0 | 0 | -.5 |
| S1 Rel Unl VR | 1 | -.5 | 0 | 1 | -.5 |
| S1 Poi Unl VR | 1 | -.5 | 1 | 0 | -.5 |
| S2 Abs Unl VR | 1 | 0 | 0 | 0 | -.5 |
| S2 Rel Unl VR | 1 | 0 | 0 | 1 | -.5 |
| S2 Poi Unl VR | 1 | 0 | 1 | 0 | -.5 |
| S3 Abs Unl VR | 1 | .5 | 0 | 0 | -.5 |
| S3 Rel Unl VR | 1 | .5 | 0 | 1 | -.5 |
| S3 Poi Unl VR | 1 | .5 | 1 | 0 | -.5 |
| S1 Abs Unl VR with belt | 1 | -.5 | 0 | 0 | .5 |
| ... | | | | | |

*Table D30.* Model s-VRvB Matrix.

The fixed effect design matrix used to fit both, VR and VR with belt exploration condition, over session, task and the 3 second time condition results. The grey columns denote all main or simple effect predictors and their matrix coding, which dropped out in the process of model selection. Note that the maximum model before step down selection included all predictors depicted here as well as all their interactions as fixed effects. The maximum model also included all fixed effect terms except interactions as random effects.
The maximum LMEM formula is:
'CondResult ~ Expl*Session*AbsPoi+Expl*Session*AbsRel+(1+Session+AbsPoi+AbsRel | Subject)'.

| | | | | | |
|---|---|---|---|---|---|
| Degrees of freedom (df) | #obs \| #fix \| #rand \| #cov | 753 | 4 | 304 | 3 |
| Fixed effects | β0(Intercept) ± SE \| CI | 53.41 ± 0.51 | 52.41 54.4 | | |
| | β1(Session) ± SE \| CI \| χ2(1) \| p | 6.06 ± 0.94 | 4.22 7.9 | 40.59 | $1.9*10^{-10}$ |
| | β2(Abs to Poi) ± SE \| CI \| χ2(1) \| p | 1.37 ± 0.64 | 0.11 2.64 | 4.53 | 0.033 |
| | β1*β3(Abs to Rel) ± SE \| CI \| χ2(1) \| p | -4.66 ± 1.44 | -7.5 -1.83 | 10.34 | 0.001 |
| Random effects | Intercept σ \| CI | 3.2 | 2.43 4.2 | | |
| | Abs to Poi σ \| CI | 1.67 | 0.24 11.43 | | |
| | Residual variance \| CI | 8.08 | 7.61 8.57 | | |

*Table D31***.** Model u-VRvB.
Results of the best LMEM fit of the VR and the VR with belt exploration condition over sessions and tasks for the unlimited time condition. #obs means total number of experimental observations. #fix means number of fixed effect coefficients. #rand means total number of random effect coefficients. #cov means number of covariance parameters. SE is standard error and CI 95% confidence interval borders. χ2(1) is the likelihood ratio statistic with 1 degree of freedom difference to the model without this fixed effect. σ is the random effect standard deviation. Here the random effects session σ (0.02) which leads to numerical instabilities and means one can and should ignore these effects. We therefore did not include them in the model.
The best fit LMEM formula is: 'Condition~1+Session+AbsPoi+Session:AbsRel+(1|Subject)'.

| Condition | β0 (Interc.) | β1 (Session) | β2(Abs to Poi) | β3(Abs to Rel) | β4 (Explor.) | β1*β3 |
|---|---|---|---|---|---|---|
| S1 Abs Unl VR | 1 | -.5 | 0 | 0 | -.5 | 0 |
| S1 Rel Unl VR | 1 | -.5 | 0 | 1 | -.5 | -.5 |
| S1 Poi Unl VR | 1 | -.5 | 1 | 0 | -.5 | 0 |
| S2 Abs Unl VR | 1 | 0 | 0 | 0 | -.5 | 0 |
| S2 Rel Unl VR | 1 | 0 | 0 | 1 | -.5 | 0 |
| S2 Poi Unl VR | 1 | 0 | 1 | 0 | -.5 | 0 |
| S3 Abs Unl VR | 1 | .5 | 0 | 0 | -.5 | 0 |
| S3 Rel Unl VR | 1 | .5 | 0 | 1 | -.5 | .5 |
| S3 Poi Unl VR | 1 | .5 | 1 | 0 | -.5 | 0 |
| S1 Abs Unl VR with belt | 1 | -.5 | 0 | 0 | .5 | 0 |

| ... |
|---|

*Table D32.* Model u-VRvB Matrix.
The fixed effect design matrix used to fit both, VR and VR with belt exploration condition, over session, task and the unlimited time condition results. The grey columns denote all main or simple effect predictors and their matrix coding, which dropped out in the process of model selection. Note that the maximum model before step down selection included all predictors depicted here as well as all their interactions as fixed effects. The maximum model also included all fixed effect terms except interactions as random effects.
The maximum LMEM formula is:
'CondResult ~ Expl*Session*AbsRel+Expl*Session*AbsPoi+(1+Session+AbsRel+AbsPoi | Subject)'.

#### Best-fit Behavioral Models

We provide details on the linear models used for evaluating the effect of independent behavioral variables on performance in all spatial tasks and time conditions and for each exploration condition. More specifically we only provide details for those models that show a significant effect for the specific behavioral variable. That is, we report those best-fit models that still include the behavioral factor after step down.

The design matrix for each model before step-down was identical to the model for that specific condition without the behavioral factor. Each behavioral variable was either directly normalized and centered or it was first transformed to a normal distribution and then that distribution was normalized and centered before being added to the design matrix. For a simplified example see Table D33.

| Condition | β0(Intercept) | β1(Task) | β2(Behavior) | β1(Task)* β2(Behavior) |
|---|---|---|---|---|
| Task 1 Participant 1 | 1 | 0 | ≈ -.5 | 0 |
| Task 2 Participant 1 | 1 | 1 | ≈ -.5 | ≈ -.5 |
| Task 1 Participant 2 | 1 | 0 | ≈ -.43 | 0 |
| Task 2 Participant 2 | 1 | 1 | ≈ -.43 | ≈ -.43 |
| … | | | | |
| Task 1 Participant N | 1 | 0 | ≈ .5 | 0 |
| Task 2 Participant N | 1 | 1 | ≈ .5 | ≈ .5 |

*Table D33.* Simple example of a normalized and centered behavioral factor being added to a design matrix.

##### Best-fit Map Model Including Percentage of Houses Seen

We provide details on the linear model used for evaluating the effect of the independent behavioral variable "percentage of houses seen" as presented in Results section 3.5.1.1. The design matrix was identical to that used for the s-Map model (u-Map model below), only the behavioral factor was added. Table D34 and D35 hold the model results and transformation of the behavioral variable.

| | | | | | |
|---|---|---|---|---|---|
| Degrees of freedom (df) | #obs (#miss) \| #fix \| #rand \| #cov | 348 (36) | 8 | 210 | 4 |
| Fixed effects | β0(Intercept) ± SE \| CI | 48.47 ± 0.84 | 46.8 50.13 | | |
| | β1(Poi to Abs) ± SE \| CI \| χ2(1) \| p | 4.01 ± 1.24 | 1.56 6.45 | 10.19 | 0.0014 |
| | β2(Poi to Rel) ± SE \| CI \| χ2(1) \| p | 6.29 ± 1.24 | 3.84 8.74 | 20.88 | $5*10^{-6}$ |
| | β1*β3(Session) ± SE \| CI \| χ2(1) \| p | 7.0 ± 2.07 | 2.91 11.09 | 11.01 | 0.0009 |
| | β2*β3(Session) ± SE \| CI \| χ2(1) \| p | 10.62 ± 2.08 | 6.53 14.72 | 23.63 | $1*10^{-6}$ |
| | β1*β3* β4(Percent) ± SE \| CI \| χ2(1) \| p | 11.79 ± 4.01 | 3.89 19.68 | 8.21 | 0.0041 |
| | β2* β3* β4 ± SE \| CI \| χ2(1) \| p | 16.36 ± 4.03 | 8.43 24.29 | 13.73 | 0.00021 |
| | β3* β4 ± SE \| CI \| χ2(1) \| p | 16.32 ± 4.99 | 6.49 26.15 | 11.01 | 0.0012 |

| | | | |
|---|---|---|---|
| Random effects | Intercept σ \| CI | 2.53 | 1.33<br>4.81 |
| | Poi to Abs \| CI | 2.24 | 0.38<br>13.08 |
| | Poi to Rel \| CI | 2.33 | 0.49<br>11.03 |
| | Residual variance \| CI | 8.36 | 7.59<br>9.2 |
| Transform function | exponential | | |

*Table D34.* Model s-Map.h.
Results of the best fit LMEM of the map exploration condition over session and task and percent of houses seen only for the 3 second time condition. #obs means total number of experimental observations. # miss is the number of missing observations due to participants excluded missing behavioral data. #fix means number of fixed effect coefficients. #rand means total number of random effect coefficients. #cov means number of covariance parameters. SE is standard error and CI 95% confidence interval borders. $\chi 2(1)$ is the likelihood ratio statistic with 1 degree of freedom difference to the model without this fixed effect. σ is the random effect standard deviation.
The best fit LMEM formula is:
'CondResult~1+PoiAbs+PoiRel+PoiAbs:Session+PoiRel:Session+PoiAbs:Percent+PoiRel:Percent+Session:Percent+(1+PoiAbs+PoiRel | Subject)'.

| | | | | | |
|---|---|---|---|---|---|
| Degrees of freedom (df) | #obs (#miss) \| #fix \| #rand \| #cov | 348<br>(18) | 6 | 210 | 4 |
| Fixed effects | β0(Intercept) ± SE \| CI | 57.51<br>± 0.8 | 55.97<br>59.04 | | |
| | β1(Session) ± SE \| CI \| χ2(1) \| p | 8.05<br>± 1.46 | 5.16<br>10.94 | 25.93 | $4*10^{-7}$ |
| | β2(Percent) ± SE \| CI \| χ2(1) \| p | 13.68<br>± 3.53 | 6.73<br>20.62 | 13.46 | 0.00024 |
| | β1*β2 ± SE \| CI \| χ2(1) \| p | 35.43<br>± 8.93 | 17.86<br>53 | 15.12 | 0.0001 |
| | β2*β3(Rel to Abs) ± SE \| CI \| χ2(1) \| p | -24.57<br>± 11.21 | -46.62<br>-2.51 | 4.75 | 0.029 |
| | β2*β4(Rel to Poi) ± SE \| CI \| χ2(1) \| p | -22.59<br>± 11.21 | -44.64<br>-0.54 | 4.03 | 0.044 |
| Random effects | Intercept σ \| CI | 4.13 | 3.05<br>5.58 | | |
| | Session σ \| CI | 2.87 | 0.52<br>15.78 | | |
| | Residual variance \| CI | 8.77 | 8.04<br>9.56 | | |
| Transform function | exponential | | | | |

*Table D35.* Model u-Map.h.
Results of the best fit LMEM of the map exploration condition over session and task and percent of houses seen only for the unlimited time condition. #obs means total number of experimental observations. #miss is the number of missing observations due to participants excluded missing behavioral data. #fix means number of fixed effect coefficients. #rand means total number of random effect coefficients. #cov means number of covariance parameters. SE is standard error and CI 95% confidence interval borders. $\chi 2(1)$ is the likelihood ratio statistic with 1 degree of freedom difference to the model without this fixed effect. σ is the random effect standard deviation.
The best fit LMEM formula is: 'CondResult~1+Session+Percent+Session:Percent+RelAbs:Percent:Session+RelPoi:Percent:Session+(1+Session | Subject)'.

**Best-fit VR Model Including Movement Speed**

We provide details on the linear model used for evaluating the effect of the independent behavioral variable "movement speed" as presented in Results section 3.5.1.3. The design matrix was identical to that used for the u-VR model, only the behavioral factor was added. Table D36 holds the model results and transformation of the behavioral variable.

| | | | | | |
|---|---|---|---|---|---|
| Degrees of freedom (df) | #obs (#miss) \| #fix \| #rand \| #cov | 384 (12) | 5 | 80 | 2 |
| Fixed effects | β0(Intercept) ± SE \| CI | 54.13 ± 0.71 | 52.73 55.53 | | |
| | β1(Session) ± SE \| CI \| χ2(1) \| p | 7.44 ± 1.37 | 4.74 10.15 | 28.17 | $1*10^{-7}$ |
| | β2(Abs to Poi) ± SE \| CI \| χ2(1) \| p | 2.14 ± 0.95 | 0.26 4.01 | 5 | 0.025 |
| | β1*β3(Abs to Rel) ± SE \| CI \| χ2(1) \| p | -6.24 ± 2.12 | -10.41 -2.06 | 8.5 | 0.0035 |
| | β3*β4(Speed) ± SE \| CI \| χ2(1) \| p | -3.29 ± 1.66 | -6.57 -0.01 | 3.86 | 0.049 |
| Random effects | Intercept σ \| CI | 2.63 | 1.66 4.18 | | |
| | Residual variance \| CI | 8.48 | 7.85 9.16 | | |
| Transform function | Speed > 2 = .5<br>Speed < 2 = -.5 | | | | |

**Table D36.** Model u-VR.s.
Results of the best fit LMEM of the VR exploration condition over session and task and movement speed only for the unlimited time condition. #obs means total number of experimental observations. # miss is the number of missing observations due to participants excluded missing behavioral data. #fix means number of fixed effect coefficients. #rand means total number of random effect coefficients. #cov means number of covariance parameters. SE is standard error and CI 95% confidence interval borders. χ2(1) is the likelihood ratio statistic with 1 degree of freedom difference to the model without this fixed effect. σ is the random effect standard deviation.
The best fit LMEM formula is: 'CondResult~1+Session+AbsPoi+AbsRel:Session+AbsRel:Speed+(1 | Subject)'.

**Best-fit VR with Belt Model Including Looking Distance**

We provide details on the linear model used for evaluating the effect of the independent behavioral variable "looking distance" as presented in Results section 3.5.1.2. The design matrix was identical to that used for the s-VRwB model, only the behavioral factor was added. Table D37 holds the model results and transformation of the behavioral variable.

| | | | | | |
|---|---|---|---|---|---|
| Degrees of freedom (df) | #obs (#miss) \| #fix \| #rand \| #cov | 354 (3) | 5 | 69 | 2 |
| Fixed effects | β0(Intercept) ± SE \| CI | 49.92 ± 0.6 | 48.73 51.25 | | |
| | β1(Poi to Abs) ± SE \| CI \| χ2(1) \| p *β3(Session) | 5.38 ± 1.87 | 1.7 9.1 | 8.2 | 0.0042 |
| | β2(Poi to Rel) ± SE \| CI \| χ2(1) \| p *β3(Session) | 3.71 ± 1.87 | 0.04 7.39 | 3.93 | 0.047 |
| | β1*β4(Distance) ± SE \| CI \| χ2(1) \| p | 14.28 ± 5.49 | 3.47 25.08 | 6.68 | 0.0097 |
| | β2*β4(Distance) ± SE \| CI \| χ2(1) \| p | 17.21 ± 5.49 | 6.4 28.01 | 9.67 | 0.0018 |
| Random effects | Intercept σ \| CI | 2.58 | 1.51 4.43 | | |
| | Residual variance \| CI | 8.54 | 7.87 9.25 | | |

| | |
|---|---|
| Transform function | none |

**Table D37.** Model s-VRwB.d.
Results of the best LMEM fit of the VR with belt exploration condition over sessions and tasks and looking distance for the 3 second time condition. #obs means total number of experimental observations. #miss is the number of missing observations due to participants excluded missing behavioral data. #fix means number of fixed effect coefficients. #rand means total number of random effect coefficients. #cov means number of covariance parameters. SE is standard error and CI 95% confidence interval borders. χ2(1) is the likelihood ratio statistic with 1 degree of freedom difference to the model without this fixed effect. σ is the random effect standard deviation.
The best fit LMEM formula is: 'Condition~1+PoiAbs:Session+PoiRel:Session+PoiAbs:Distance+PoiRel:Distance+(1|Subject)'.

**Best-fit Models Including Self-report**

We provide details on the linear models used for evaluating the effect of the independent behavioral variable "self-report" as presented in Results section 3.5.1.4. Self-report splits into three factors, namely general-egocentric, survey and cardinal. We provide those models in which one of the factors showed a significant effect. The order is general, survey and cardinal and three seconds to unlimited for each.

*Best-fit VR Model Including Self-report Survey*

| | | | | | |
|---|---|---|---|---|---|
| Degrees of freedom (df) | #obs (#miss) \| #fix \| #rand \| #cov | 348 (48) | 3 | 74 | 2 |
| Fixed effects | β0(Intercept) ± SE \| CI | 48.92 ± 0.48 | 47.96 49.87 | | |
| | β1(Survey) ± SE \| CI \| χ2(1) \| p | 5.45 ± 2.52 | 0.48 10.41 | 4.40 | 0.035 |
| | β1* β2(Session)* β3(Abs to Poi) ± SE \| CI \| χ2(1) \| p | 21.78 | 7.18 36.38 | 8.44 | 0.0036 |
| Random effects | Survey σ \| CI | 8.4 | 4.44 15.9 | | |
| | Residual variance \| CI | 8.12 | 7.51 8.77 | | |
| Transform function | none | | | | |

*Table D38.* Model s-VR.rs.
Results of the best fit LMEM of the VR exploration condition over session and task and self-report survey only for the three seconds time condition. #obs means total number of experimental observations. #miss is the number of missing observations due to participants excluded missing behavioral data. #fix means number of fixed effect coefficients. #rand means total number of random effect coefficients. #cov means number of covariance parameters. SE is standard error and CI 95% confidence interval borders. χ2(1) is the likelihood ratio statistic with 1 degree of freedom difference to the model without this fixed effect. σ is the random effect standard deviation.
The LMEM formula is: 'CondResult~1+Survey+Survey:Session:AbsPoi+(Survey|Subject)'.

| | | | | | |
|---|---|---|---|---|---|
| Degrees of freedom (df) | #obs (#miss) \| #fix \| #rand \| #cov | 348 (48) | 6 | 74 | 2 |
| Fixed effects | β0(Intercept) ± SE \| CI | 53.14 ± 0.65 | 51.85 54.43 | | |
| | β1(Session) ± SE \| CI \| χ2(1) \| p | 6.37 ± 1.38 | 3.65 9.09 | 20.32 | $7*10^{-6}$ |
| | β2(Abs to Poi) ± SE \| CI \| χ2(1) \| p | 2.81 ± 0.97 | 0.89 4.73 | 8.18 | 0.0042 |
| | β3(Survey) ± SE \| CI \| χ2(1) \| p | 11.07 ± 2.70 | 5.74 16.39 | 14.69 | 0.00012 |
| | β1*β4(Abs to Rel) ± SE \| CI \| χ2(1) \| p | -6.39 ± 2.15 | -10.64 -2.14 | 8.65 | 0.0032 |

| | | | | | |
|---|---|---|---|---|---|
| | β1*β3 ± SE \| CI \| χ2(1) \| p | 12.36 ± 5.0 | 2.51 22.21 | 6.01 | 0.014 |
| Random effects | Survey σ \| CI | 7.56 | 3.86 14.83 | | |
| | Residual variance \| CI | 8.31 | 7.69 8.97 | | |
| Transform function | none | | | | |

*Table D39.* Model u-VR.rs.
Results of the best fit LMEM of the VR exploration condition over session and task and self-report survey only for the unlimited time condition. #obs means total number of experimental observations. #miss is the number of missing observations due to participants excluded missing behavioral data. #fix means number of fixed effect coefficients. #rand means total number of random effect coefficients. #cov means number of covariance parameters. SE is standard error and CI 95% confidence interval borders. χ2(1) is the likelihood ratio statistic with 1 degree of freedom difference to the model without this fixed effect. σ is the random effect standard deviation.
The best fit LMEM formula is: 'CondResult~1+Survey+Session+AbsPoi+AbsRel:Session+AbsRel:Survey+(Survey | Subject)'.

*Best-fit VR Model Including Self-report Cardinal*

| Degrees of freedom (df) | #obs (#miss) \| #fix \| #rand \| #cov | 348 (48) | 2 | 74 | 2 |
|---|---|---|---|---|---|
| Fixed effects | β0(Intercept) ± SE \| CI | 48.91 ± 0.50 | 47.91 49.91 | | |
| | β1(Cardinal) ± SE \| CI \| χ2(1) \| p | 3.39 ± 1.57 | 0.29 6.49 | 4.43 | 0.035 |
| Random effects | Cardinal σ \| CI | 6.18 | 3.28 11.67 | | |
| | Residual variance \| CI | 8.17 | 7.54 8.85 | | |
| Transform function | natural logarithm | | | | |

*Table D40.* Model s-VR.rc.
Results of the best fit LMEM of the VR exploration condition over session and task and self-report cardinal only for the three seconds time condition. #obs means total number of experimental observations. #miss is the number of missing observations due to participants excluded missing behavioral data. #fix means number of fixed effect coefficients. #rand means total number of random effect coefficients. #cov means number of covariance parameters. SE is standard error and CI 95% confidence interval borders. χ2(1) is the likelihood ratio statistic with 1 degree of freedom difference to the model without this fixed effect. σ is the random effect standard deviation.
The LMEM formula is: 'CondResult~1+Cardinal+(Cardinal|Subject)'.

| Degrees of freedom (df) | #obs (#miss) \| #fix \| #rand \| #cov | 348 (48) | 5 | 74 | 2 |
|---|---|---|---|---|---|
| Fixed effects | β0(Intercept) ± SE \| CI | 53.40 ± 0.69 | 52.02 54.77 | | |
| | β1(Session) ± SE \| CI \| χ2(1) \| p | 6.57 ± 1.41 | 3.79 9.35 | 20.85 | $5*10^{-6}$ |
| | β2(Abs to Poi) ± SE \| CI \| χ2(1) \| p | 2.82 ± 0.98 | 0.89 4.76 | 8.16 | 0.0042 |
| | β1*β3(Abs to Rel) ± SE \| CI \| χ2(1) \| p | -6.89 ± 2.17 | -11.17 -2.61 | 9.87 | 0.0016 |
| | β3*β4(Cardinal) ± SE \| CI \| χ2(1) \| p | 8.10 ± 2.32 | 3.54 12.67 | 11.82 | 0.00058 |

| | | | |
|---|---|---|---|
| Random effects | Intercept σ \| CI | 2.16 | 1.14<br>4.09 |
| | Residual variance \| CI | 8.36 | 7.71<br>9.06 |
| Transform function | natural logarithm | | |

*Table D41.* Model u-VR.rc.
Results of the best fit LMEM of the VR exploration condition over session and task and self-report cardinal only for the unlimited time condition. #obs means total number of experimental observations. #miss is the number of missing observations due to participants excluded missing behavioral data. #fix means number of fixed effect coefficients. #rand means total number of random effect coefficients. #cov means number of covariance parameters. SE is standard error and CI 95% confidence interval borders. $\chi2(1)$ is the likelihood ratio statistic with 1 degree of freedom difference to the model without this fixed effect. σ is the random effect standard deviation.
The best fit LMEM formula is: 'CondResult~1+Session+AbsPoi+AbsRel:Session+AbsRel:Cardinal+(1 | Subject)'.

*Best-fit Map Model Including Self-report General*

| | | | | | |
|---|---|---|---|---|---|
| Degrees of freedom (df) | #obs (#miss) \| #fix \| #rand \| #cov | 366<br>(18) | 3 | 144 | 3 |
| Fixed effects | β0(Intercept) ± SE \| CI | 57.06<br>± 0.87 | 55.35<br>58.78 | | |
| | β1(Session) ± SE \| CI \| χ2(1) \| p | 6.88<br>± 1.5 | 3.92<br>9.83 | 19.03 | 0.000013 |
| | β2(General) ± SE \| CI \| χ2(1) \| p | 8.36<br>± 3.17 | 2.12<br>14.59 | 6.49 | 0.01 |
| Random effects | Intercept σ \| CI | 4.79 | 3.68<br>6.24 | | |
| | Session σ \| CI | 4.46 | 1.98<br>10.04 | | |
| | Residual variance \| CI | 8.83 | 8.12<br>9.6 | | |
| Transform function | exponential | | | | |

*Table D42.* Model u-Map.rg.
Results of the best fit LMEM of the map exploration condition over session and task and self-report general only for the unlimited time condition. #obs means total number of experimental observations. #miss is the number of missing observations due to participants excluded missing behavioral data. #fix means number of fixed effect coefficients. #rand means total number of random effect coefficients. #cov means number of covariance parameters. SE is standard error and CI 95% confidence interval borders. $\chi2(1)$ is the likelihood ratio statistic with 1 degree of freedom difference to the model without this fixed effect. σ is the random effect standard deviation.
The best fit LMEM formula is: 'CondResult~1+Session+General+(1+Session | Subject)'.

*Best-fit Map Model Including Self-report Survey*

| | | | | | |
|---|---|---|---|---|---|
| Degrees of freedom (df) | #obs (#miss) \| #fix \| #rand \| #cov | 366<br>(18) | 4 | 144 | 3 |
| Fixed effects | β0(Intercept) ± SE \| CI | 57.59<br>± 0.87 | 55.88<br>59.31 | | |
| | β1(Session) ± SE \| CI \| χ2(1) \| p | 7.57<br>± 1.5 | 4.60<br>10.54 | 22.51 | $2*10^{-6}$ |
| | β2(Survey) ± SE \| CI \| χ2(1) \| p | 11.31<br>± 3.59 | 4.24<br>18.38 | 9.08 | 0.0025 |
| | β1*β2 ± SE \| CI \| χ2(1) \| p | 12.26<br>± 6.13 | 0.19<br>24.32 | 3.89 | 0.048 |

| | | | |
|---|---|---|---|
| Random effects | Intercept σ \| CI | 4.66 | 3.56<br>6.11 |
| | Session σ \| CI | 4.10 | 1.62<br>10.35 |
| | Residual variance \| CI | 8.84 | 8.13<br>9.61 |
| Transform function | none | | |

*Table D43.* Model u-Map.rs.
Results of the best fit LMEM of the map exploration condition over session and task and self-report survey only for the unlimited time condition. #obs means total number of experimental observations. #miss is the number of missing observations due to participants excluded missing behavioral data. #fix means number of fixed effect coefficients. #rand means total number of random effect coefficients. #cov means number of covariance parameters. SE is standard error and CI 95% confidence interval borders. χ2(1) is the likelihood ratio statistic with 1 degree of freedom difference to the model without this fixed effect. σ is the random effect standard deviation.
The best fit LMEM formula is: 'CondResult~1+Session+Survey+Session:Survey+(1+Session | Subject)'.

*Best-fit Map Models Including Self-report Cardinal*

| Degrees of freedom (df) | #obs (#miss) \| #fix \| #rand \| #cov | 366 (18) | 6 | 288 | 5 |
|---|---|---|---|---|---|
| Fixed effects | β0(Intercept) ± SE \| CI | 48.04<br>± 0.8 | 46.45<br>49.63 | | |
| | β1(Poi to Abs) ± SE \| CI \| χ2(1) \| p | 3.61<br>± 1.18 | 1.29<br>5.93 | 9.2 | 0.0024 |
| | β2(Poi to Rel) ± SE \| CI \| χ2(1) \| p | 5.28<br>± 1.21 | 2.89<br>7.68 | 16.07 | $6*10^{5}$ |
| | β1*β3(Session) ± SE \| CI \| χ2(1) \| p | 5.56<br>± 2.07 | 1.48<br>9.65 | 6.96 | 0.0083 |
| | β2*β3(Session) ± SE \| CI \| χ2(1) \| p | 7.61<br>± 2.09 | 3.49<br>11.74 | 12.09 | 0.0005 |
| | β2*β4(Cardinal) ± SE \| CI \| χ2(1) \| p | 6.76<br>± 2.8 | 1.25<br>12.27 | 5.32 | 0.021 |
| Random effects | Intercept σ \| CI | 2.46 | 1.28<br>4.71 | | |
| | Session \| CI | 6.89 | 4.46<br>10.64 | | |
| | Poi to Abs \| CI | 3.18 | 1.3<br>7.75 | | |
| | Poi to Rel \| CI | 3.71 | 1.95<br>7.09 | | |
| | Residual variance \| CI | 8.59 | 7.86<br>9.38 | | |
| Transform function | natural logarithm | | | | |

*Table D44.* Model s-Map.rc.
Results of the best fit LMEM of the map exploration condition over session and task and self-report cardinal only for the three seconds time condition. #obs means total number of experimental observations. #miss is the number of missing observations due to participants excluded missing behavioral data. #fix means number of fixed effect coefficients. #rand means total number of random effect coefficients. #cov means number of covariance parameters. SE is standard error and CI 95% confidence interval borders. χ2(1) is the likelihood ratio statistic with 1 degree of freedom difference to the model without this fixed effect. σ is the random effect standard deviation.
The best fit LMEM formula is: 'CondResult~1+PoiAbs+PoiRel+PoiAbs:Session+PoiRel:Session+PoiRel:Cardinal+(1+PoiAbs+PoiRel+Session | Subject)'.

<table>
<tr><td>Degrees of freedom (df)</td><td>#obs (#miss) | #fix | #rand | #cov</td><td>366<br>(18)</td><td>4</td><td>144</td><td>3</td></tr>
<tr><td>Fixed effects</td><td>$\beta_0$(Intercept) ± SE | CI</td><td>57.56<br>± 0.8</td><td>55.97<br>59.14</td><td></td><td></td></tr>
<tr><td></td><td>$\beta_1$(Session) ± SE | CI | $\chi^2(1)$ | p</td><td>7.52<br>± 1.37</td><td>4.82<br>10.23</td><td>25.93</td><td>$4*10^{-7}$</td></tr>
<tr><td></td><td>$\beta_2$(Cardinal) ± SE | CI | $\chi^2(1)$ | p</td><td>11.97<br>± 2.41</td><td>7.23<br>16.71</td><td>21.38</td><td>$4*10^{-6}$</td></tr>
<tr><td></td><td>$\beta_1*\beta_2$ ± SE | CI | $\chi^2(1)$ | p</td><td>15.36<br>± 4.05</td><td>7.39<br>23.33</td><td>13.22</td><td>0.00027</td></tr>
<tr><td>Random effects</td><td>Intercept σ | CI</td><td>4.26</td><td>3.21<br>5.67</td><td></td><td></td></tr>
<tr><td></td><td>Session σ | CI</td><td>2.54</td><td>0.31<br>20.67</td><td></td><td></td></tr>
<tr><td></td><td>Residual variance | CI</td><td>8.81</td><td>8.1<br>9.58</td><td></td><td></td></tr>
<tr><td>Transform function</td><td colspan="5">natural logarithm</td></tr>
</table>

*Table D45.* Model u-Map.rc.
Results of the best fit LMEM of the map exploration condition over session and task and self-report cardinal only for the unlimited time condition. #obs means total number of experimental observations. #miss is the number of missing observations due to participants excluded missing behavioral data. #fix means number of fixed effect coefficients. #rand means total number of random effect coefficients. #cov means number of covariance parameters. SE is standard error and CI 95% confidence interval borders. $\chi^2(1)$ is the likelihood ratio statistic with 1 degree of freedom difference to the model without this fixed effect. σ is the random effect standard deviation.
The best fit LMEM formula is: 'CondResult~1+Session+Cardinal+Session:Cardinal+(1+Session | Subject)'.

**Best-fit VR with Belt Model Including Self-report Cardinal**

<table>
<tr><td>Degrees of freedom (df)</td><td>#obs (#miss) | #fix | #rand | #cov</td><td>336<br>(21)</td><td>4</td><td>67</td><td>2</td></tr>
<tr><td>Fixed effects</td><td>$\beta_0$(Intercept) ± SE | CI</td><td>49.6<br>± 0.64</td><td>48.33<br>50.87</td><td></td><td></td></tr>
<tr><td></td><td>$\beta_1$(Poi to Abs) ± SE | CI | $\chi^2(1)$ | p<br>*$\beta_3$(Session)</td><td>5.54<br>± 1.93</td><td>1.73<br>9.35</td><td>8.1</td><td>0.0044</td></tr>
<tr><td></td><td>$\beta_2$(Poi to Rel) ± SE | CI | $\chi^2(1)$ | p<br>*$\beta_3$(Session)</td><td>3.73<br>± 1.93</td><td>-0.07<br>7.54</td><td>3.69</td><td>0.054 (ns)</td></tr>
<tr><td></td><td>$\beta_3*\beta_4$(Cardinal) ± SE | CI | $\chi^2(1)$ | p</td><td>10.11<br>± 3.36</td><td>3.49<br>16.74</td><td>8.89</td><td>0.0028</td></tr>
<tr><td>Random effects</td><td>Intercept σ | CI</td><td>2.77</td><td>1.64<br>4.7</td><td></td><td></td></tr>
<tr><td></td><td>Residual variance | CI</td><td>8.59</td><td>7.9<br>9.34</td><td></td><td></td></tr>
<tr><td>Transform function</td><td colspan="5">natural logarithm</td></tr>
</table>

**Table D46.** Model s-VRwB.rc.
Results of the best LMEM fit of the VR with belt exploration condition over sessions and tasks for the 3 second time condition. #obs means total number of experimental observations. #miss is the number of missing observations due to participants excluded missing behavioral data. #fix means number of fixed effect coefficients. #rand means total number of random effect coefficients. #cov means number of covariance parameters. SE is standard error and CI 95% confidence interval borders. $\chi^2(1)$ is the likelihood ratio statistic with 1 degree of freedom difference to the model without this fixed effect. σ is the random effect standard deviation.
The best fit LMEM formula is: 'Condition~1+PoiAbs:Session+PoiRel:Session+Session:Cardinal+(1|Subject)
